# Genetic variation in heat tolerance of the coral *Platygyra daedalea* indicates potential for adaptation to ocean warming

**DOI:** 10.1101/2020.10.13.337089

**Authors:** Holland Elder, Virginia Weis, Jose Montalvo-Proano, Veronique J.L Mocellin, Andrew H. Baird, Eli Meyer, Line K. Bay

**Author notes:** These authors contributed equally and share last authorship (Eli Meyer, Line K. Bay).

## Abstract

Climate change induced increases in global ocean temperature represent the greatest threat to the persistence of reef ecosystems and most coral populations are projected to experience temperatures above their current bleaching thresholds annually by 2050. Adaptation to higher temperatures is necessary if corals are to persist in a warming future. While many aspects of heat stress have been well studied, few data are available for predicting the capacity for adaptive cross-generational responses in corals. Consistent sets of heat tolerant genomic markers that reliably predict thermal tolerance have yet to be identified. To address this knowledge gap, we quantified the heritability and genetic variation associated with heat tolerance in *Platygyra daedalea* from the Great Barrier Reef. We tracked the survival of quantitative genetic crosses of larvae in a heat tolerance selection experiment. We also identified allelic shifts in heat-selected survivors compared with paired, non-selected controls. The narrow-sense heritability of survival under heat stress was 0.66 and a total of 1,069 single nucleotide polymorphisms (SNPs) were associated with different survival probabilities. While 148 SNPs were shared between several experimental crosses, no common SNPs were identified for all crosses suggesting that specific combinations of many markers are responsible for heat tolerance. However, we found two regions that overlap with previously identified loci associated with heat tolerance in Persian Gulf populations of *P. daedalea* reinforcing the importance of these markers for heat tolerance. These results illustrate the importance of high heritability and the complexity of the genomic architecture underpinning host heat tolerance. These findings suggest that this *P. daedalea* population has the genetic prerequisites for adaptation to increasing temperatures. This study also provides knowledge for the development of high throughput genomic tools to screen for variation within and across populations to enhance adaptation through assisted gene flow and assisted migration.

## Introduction

Rising ocean temperatures induced by anthropogenic climate change have increased the frequency and severity of coral bleaching events worldwide to the point that it is recognized as the primary threat to coral reefs (Hutchings et al., 2019; Souter et al., 2020). Bleaching is the process in which dinoflagellate symbionts (Family Symbiodiniaceae) are lost or expelled from host corals (Lajeunesse et al., 2018). As symbiont photosynthesis provides the majority of host nutritional requirements, mass bleaching caused by marine heat waves can result in rapid and widespread coral mortality (Glynn, 1993). Global and regional bleaching events have resulted in a substantial loss of coral cover worldwide over the last few decades (Hughes et al., 2017, 2018). Over the last five years alone, 30% of reefs in the Great Barrier Reef (GBR) showed decreases in coral cover (Hughes et al., 2019; Hughes et al., 2018; Long Term Reef Monitoring Program, 2019).

For coral reef ecosystems to persist in an ocean that will continue to warm for the foreseeable future, corals will need to rapidly become more tolerant or resilient to higher temperatures. Consequently, there is an urgent need to understand the adaptive capacity of corals to this threat across ocean basins and populations to inform conventional management approaches. This knowledge is also required for restoration and adaptation programs using assisted gene flow and assisted migration to enhance the tolerance of local populations (van Oppen et al., 2015) as adaptive capacity can help determine which populations should be used to stock nursery and breeding programs. Working knowledge of the genomic regions that influence heat tolerance, the extent to which coral host genetics contributes to heat tolerance, and its cross generational persistence in a population will be essential to predict corals’ responses to climate change into the future.

Mechanisms of thermal and bleaching tolerance are diverse, and both hosts and symbionts are known to contribute to the holobiont phenotype. For example, different symbiont species can confer different thermal tolerances in the same host species (Fuller et al., 2020). In addition, the presence of low background densities of heat tolerant symbiont species can aid in recovery from heat stress (Bay et al., 2016; Berkelmans & van Oppen, 2006; Manzello et al., 2019; Oliver & Palumbi, 2011; Silverstein et al., 2014). Many studies have also documented genetic variation in coral host heat tolerance that supports the potential for adaptation (Bay et al., 2017; Dixon et al., 2015; Howells et al., 2016; Kenkel et al., 2013; Matz et al., 2020; Meyer et al., 2009; Woolsey et al., 2014). Differential gene expression has been associated with heat tolerance in a number of coral species (e.g., Barshis et al., 2013; Kenkel et al., 2016; Ruiz-jones & Palumbi, 2017; Traylor-Knowles et al., 2017). While some studies have identified genomic markers in coding and non-coding genes that are associated with a heat tolerant phenotype, yet no consensus set of ‘candidate genes’ has emerged (Bay & Palumbi, 2014; Dixon et al., 2015; Dziedzic et al., 2019; Fuller et al., 2020; Jin et al., 2016) and the genetic architecture underpinning heat tolerance remains largely unknown.

Coral heat tolerance is likely a complex polygenic trait controlled by many alleles of small effect which complicates efforts to identify loci in part because of the statistical challenge of distinguishing signal from noise (Fuller et al., 2020). Polygenicity is well documented in diverse traits in agricultural species, model organisms, and humans (reviewed in Sella & Barton, 2019) but remains difficult to detect because most genotyping methods are biased towards detecting loci of large effect (Wellenreuther & Hansson, 2016). Although methodological challenges remain, genome-wide association studies have yielded insight into the genomic basis of key performance traits in a number of species. For example, in plants, allele frequency analysis has identified markers associated with traits such as yield, drought tolerance and heat tolerance (Singh et al., 2017; Venuprasad et al., 2009; Vikram et al., 2012; Wang et al., 2019). Consequently, great interest remains in identifying and developing genomic marker sets for conservation, restoration and adaptation applications (Baums et al., 2019).

Even in the absence of predictive markers, knowledge of trait heritability can inform breeding programs (Falconer & Mackay, 1996; Lynch & Walsh, 1998) and the development of predictive models (Gienapp et al., 2013). If variation in a fitness-related phenotype is heritable and not otherwise constrained, then selection can act to further propagate it within populations (Charmantier & Garant, 2005). Narrow-sense heritability (*h*^2^) describes the total phenotypic variance in a trait that can be attributed to parental or additive genetics and is used to determine the genetic contribution to traits such as heat tolerance (Falconer & Mackay, 1996; Lynch & Walsh, 1998). To model the persistence of a trait in a population with the breeder’s equation, estimates of narrow-sense heritability are essential for calculating selection differentials (Falconer & Mackay, 1996; Lynch & Walsh, 1998). Specifically, estimates of narrow-sense heritability of heat tolerance in corals will allow us to calculate potential rates of adaptation under various selection scenarios.

Current estimates of narrow-sense heritability suggest that the genetic variation to support adaptation of increased heat tolerance is present in coral populations in many different regions. For example, a heritability value of 0.89 was found for bleaching resistance in *Orbicella faveolata* in the Caribbean (Dziedzic et al., 2019) and values between 0.48 and 0.74 for heat tolerance in *Platygyra daedalea* in the Persian Gulf (Kirk et al., 2018). Heritability was similarly high for two *Acropora* species from the GBR: (0.87) for heat tolerance in the larvae of the coral *A. millepora* (Dixon et al., 2015) and (0.93) for survival during heat stress in the larvae of *A. spathulata* (Quigley et al., 2020). However, heritability can vary substantially between species, as well as between populations of conspecifics, and is influenced by the environment in which it is measured (Falconer & Mackay, 1996; Lynch & Walsh, 1998). Therefore, a heritability estimate from the Persian Gulf should not be used to estimate selection for heat tolerance or potential for adaptation in a population on the GBR, but a comparison among regions can inform understanding of the evolutionary potential of the species. To understand the heritability of heat tolerance, selection for heat tolerance, and the potential for adaptation in GBR populations, more heritability estimates of heat tolerance from populations on the GBR are needed (Bairos-Novak et al., 2021).

Cross ocean basin comparisons of heritability and the genetic basis of heat tolerance are important for assessing whether results from one region can be extrapolated to another. *Platygyra daedalea* is a good candidate species for this type of study because it is widely distributed in the Indo-Pacific and also occurs in the much warmer Persian Gulf (D’Angelo et al., 2015). The Gulf regularly experiences temperatures of 33 - 34°C in the summer and the bleaching threshold for Gulf populations of *P. daedalea* is between 35 and 36°C (Howells et al., 2016; Riegl et al., 2011). Indeed, the bleaching threshold temperature for the coral assemblages is 3 - 7°C higher in the Gulf than in other regions (29 - 32°C), including the GBR (Berkelmans, 2002; Coles & Riegl, 2013; Hoegh-Guldberg, 1999). In addition, heritability estimates and preliminary data on genomic associations exist for a population of *P. daedalea* in the Persian Gulf (Kirk et al., 2018). This provides a unique opportunity to compare heritability and the genomic basis of heat tolerance between populations from these relatively warmer and cooler environments, which is important for understanding how increased thermal tolerance has evolved and for determining whether the genetic mechanisms underpinning heat tolerance in this species are conserved across ocean basins.

To understand the potential for a GBR population of *P. daedalea* to persist in a warming environment through selection and genetic adaptation we estimated narrow-sense heritability and identified genomic regions associated with heat tolerance. We also compared the genomic basis of heat tolerance in this species on the GBR to those of the Persian Gulf *P. daedalea* (Kirk et al., 2018). Estimating heritability of heat tolerance allowed us to quantify the host genetic contribution to this trait. We identified genomic markers associated with heat tolerant phenotypes by conducting a heat selection experiment and then assessing changes in allele frequency in the survivors. Sequencing our heat-selected pools of bi-parental crosses allowed us to identify markers associated with heat tolerance and examine how widespread these markers were across families in both genic and intergenic genomic regions. This study addresses fundamental questions regarding the adaptive potential of corals to heat stress and provides evidence of their ability to survive and persist in a changing environment.

## Materials and Methods

### Experimental crosses and larval culture

Six colony fragments of *Platygyra daedalea* were collected from Esk Reef, an inshore reef in the central region of the Great Barrier Reef (18° 46’21.95” S, 146° 31’ 19.33” E) under Australian Institute of Marine Science (AIMS) permit G12/35236.1. The corals were maintained in the National Sea Simulator at AIMS at Esk Reef ambient temperature (27°C) in flow-through sea water aquaria with 25% reduced ambient sunlight. In November 2016 (the 5^th^ night after the full moon, at 18:30) the six parental colonies were isolated in ∼60L bins filled with 0.04-μm filtered sea water (FSW) (Weeriyanun et al., 2021). Colonies spawned between 18:30-19:00 GMT+10. Gamete bundles were collected and washed with FSW. Sperm and eggs were then separated using 60 μm mesh sieves and eggs were thrice rinsed to eliminate residual sperm. We combined gametes from pairs of colonies according to a partial diallel design (Figure 1) to produce 10 families (hereafter termed crosses). A full diallel cross was not possible because colonies D and E did not produce enough eggs for the purposes of this experiment. Each cross was maintained at a density of 0.5 to 1.0 planula larvae per ml in separate 12L culture vessels with flow-through FSW at 27°C and gentle aeration.

**Figure 1.**
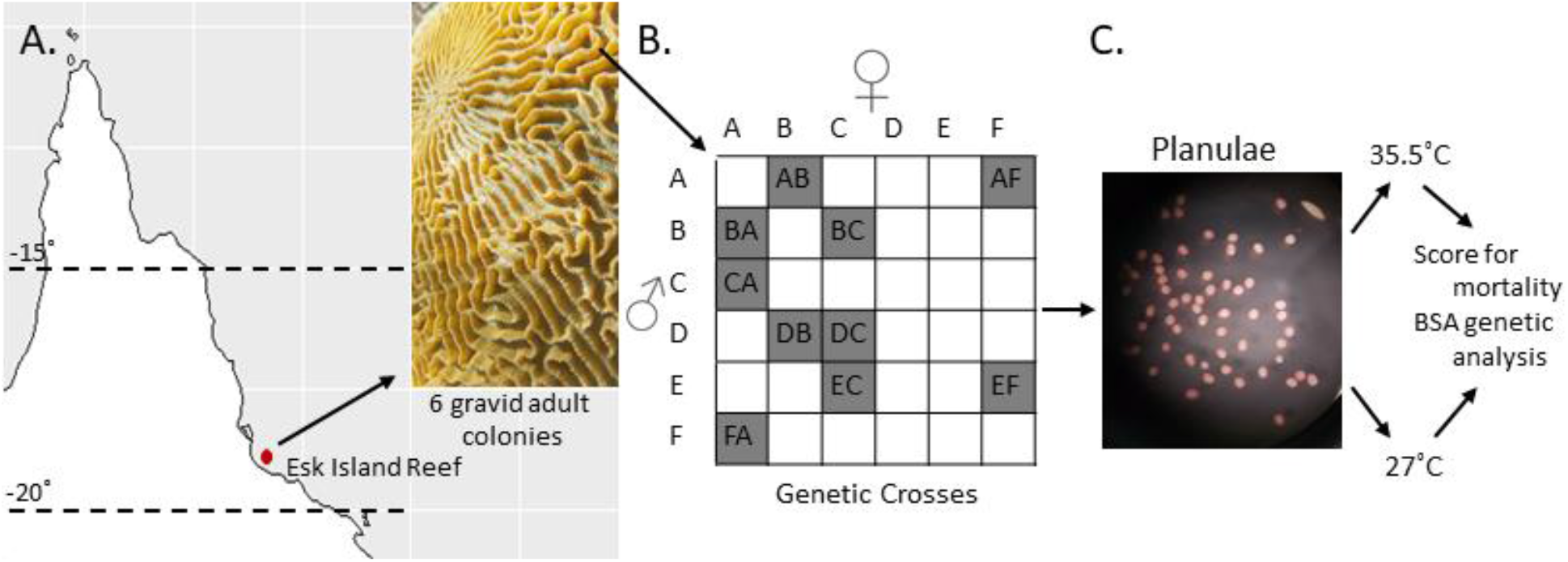
Crossing design and heat stress experiments. (A) Six *Platygyra daedalea* colonies were sampled from Esk Reef on the GBR. (B) Crossing design used to produce ten crosses among those parental colonies. (C) Representative micrograph (10X) of planula larvae used for these experiments (53 hours post-fertilization).

### Heat stress experiments

To quantify the variation in coral heat tolerance, we measured larval survival during controlled heat stress exposure following (Dixon et al., 2015). Six replicates of 20 larvae from each cross were randomly allocated to each of two temperature treatments: ambient reef temperatures from the site of collection (27°C, Supplemental Figure 1) or an acute heat stress (35.5°C). Larval replicates were contained in individual mesh bottomed 20ml wells (to allow for water circulation) within 60L flow through FSW tanks (N = 6 control and N = 6 heat per cross). A ramping rate of 1°C per hour was used to reach 35.5°C and water temperatures were maintained thereafter by computer-controlled heaters. Mortality was monitored by manually counting with a hand counter the number of surviving larvae in each well every 12 hours for the next ten days. Larval survival data can be found at https://github.com/hollandelder/Platygyradaedalea_HeritabiltyandMarkersofThermalTolerance.

### Statistical analysis of heat tolerance

Survival under heat stress was modeled using a Kaplan Meier Survival Analysis and a Cox Proportional Hazards (CPH) model that compared larval survival between crosses at 35.5°C and 27°C (Cox, 2007; Kaplan and Meier, 2017) using the Survival package in R (Therneau, 2015). Our model fit individual survival, allowing slopes to vary for temperature, cross, and the interaction of temperature and cross. We also calculated the uncensored survival probabilities as in Kirk et al., 2018. The code for the Kaplan Meier survival plots and CPH model can be referenced at https://github.com/hollandelder/Platygyradaedalea_HeritabiltyandMarkersofThermalTolerance.

CPH in the Survival package in R does not account for random effects, so we also used a generalized linear mixed model (GLMM) to determine if position contributed to variation in survival. Please see the supplemental text for complete GLMM analysis details.

We used survival fractions at the experimental endpoint and Kaplan Meier survival probabilities to estimate narrow-sense heritability of the variation in heat tolerance (Falconer & Mackay, 1996; Wilson et al., 2010) using a restricted maximum likelihood (REML) mixed-effects model implemented in the software package WOMBAT (Meyer, 2007) with the fixed effects of temperature, sire, and dam.

### Heat stress selection experiment to investigate the genomic basis for heat stress tolerance

We conducted a selection experiment to identify changes in allele frequency in response to acute heat exposure. Four replicates of 500 larvae per cross were allocated into 1L flow-through culture containers for a total of 2000 larvae in each of two temperature treatments (27°C and 35.5°C). Treatment conditions and ramping profiles were identical to those described for the survival analysis. We sampled 100 larvae from each replicate for a total of 400 larvae per treatment after 53 h, the time at which more than 50% mortality was evident in most crosses, and larvae were preserved in 100% ethanol for subsequent genetic analysis.

### Genomic library preparation

Genomic DNA from each replicate pool was extracted from preserved samples using the Omega Bio-tek E.Z.N.A. Tissue Kit (Omega Bio-tek, Norcross, GA) and genotyped using 2bRAD, a sequencing-based approach for SNP genotyping (Wang et al., 2012) that has been previously applied in corals https://github.com/z0on/2bRAD_denovo (Dixon et al., 2015; Howells et al., 2016). Libraries were sequenced on four lanes of Illumina HiSeq2500 (50bp SE) at Ramaciotti Centre for Genomics at the University of New South Wales (Sydney, Australia).

### Sequence processing

Raw reads were quality filtered to remove any that had ten or more bases with a Q-score < 30. This is a stringent filter, but 95% of reads were retained for most samples at this threshold (Supplemental Table 1). Reads exhibiting a 12bp match to the Illumina sequencing adaptors were also removed. Remaining high quality reads were mapped against a 2bRAD reference previously developed from larvae of Persian Gulf *P. daedalea* (SRA accession: SRP066627, Howells et al., 2016) to facilitate subsequent cross-region comparisons using the gmapper command of the SHRiMP software package (Rumble et al., 2009) with flags --qv-offset 33 -Q -- strata -o 3 -N 1 as in (Kirk et al., 2018). On average, 91% of reads mapped to the reference (Supplemental Table 1). After alignment to the reference, we further filtered reads with ambiguous (i.e. those mapping equally well to two or more different regions in the reference) or weak matches (those that did not span at least 33 of a 36 base sequence and matched fewer than 30 bases within that span of a sequence). SNPs were called using the CallGenotype.pl script found at https://github.com/Eli-Meyer/2bRAD_utilities as in (Dziedzic et al., 2019; Kirk et al., 2018) using observed nucleotide frequencies and a minor allele frequency of 0.25 to be considered heterozygous. Monomorphic loci were removed as well as those occurring in fewer than 25% of samples within a cross. From here on, we refer to mapped, quality filtered, and genotyped reads as tags. Samples with too many missing tags (<10,000 remaining) following read filtering were also removed. We required that each tag be genotyped in at least seven out of eight replicates for each cross to compare alleles at that locus across replicates in each treatment. Tags with more than two SNPs were also removed because they are likely to come from repetitive regions (Treangen & Salzberg, 2012). Finally, we down-sampled to one SNP per tag to control for non-independence and linkage but prioritized minimizing the number of missing SNPs. That is to say, the SNP found in most samples for a given tag was the one kept. If a sample did not have a SNP in the same base pair position in the tag compared to other copies of the tag, the SNP in the alternative position was kept. All 2bRAD genotyping and sequence filtering was conducted using scripts available at (https://github.com/Eli-Meyer/2brad_utilities).

### Allele frequency analysis

We then compared allele frequencies between control larvae and the survivors of heat stress in replicate samples from each treatment within each cross. Since these samples represented pools of large numbers of larvae (100) rather than single diploid individuals, we followed the same general approach previously described as Pool-Seq (Kofler et al., 2017). We applied a stringent coverage threshold of 80x within cross to minimize sampling error that is likely to distort estimates of allele frequencies at lower coverage considering the number of individuals in each sample (Zhu et al., 2012). To find the percentage of loci not genotyped in all crosses following filtering, we extracted and compared the number of tags to all SNPs genotyped in each cross using scripts available at https://github.com/hollandelder/Platygyradaedalea_HeritabiltyandMarkersofThermalTolerance.

We interpreted SNP frequencies at a locus as allele frequencies in the pool. To test for differences in allele frequencies between control larvae and those surviving heat stress, we used logistic regression to analyze the number of observations of each allele in each sample with generalized linear models in R (glm with argument “cross=binomial(logit)”) where treatment was a fixed effect (R Core Team, 2017). We used a Benjamini-Hochberg false discovery rate (FDR) correction to adjust p-values (Waite & Campbell, 2006). An allele was determined to be significant at an FDR-corrected p-value ≤ 0.05. We also calculated the proportion of alleles that increased or decreased in frequency as a result of heat stress treatment for each cross.

### Comparison of genotyped SNPs between crosses

All scripts and code for recreating the analyses described below can be found at https://github.com/hollandelder/Platygyradaedalea_HeritabiltyandMarkersofThermalTolerance. Pearson correlation tests as implemented in the R package ggpubr (Kassambara, 2020) tested for correlations between the number of significant markers and the number of markers genotyped in each cross, as well as the number of significant markers and the mean survival following ten days of heat stress. We calculated the genetic distance between sets of parents by conducting pairwise comparisons of the high coverage loci using a custom Perl script which tested for numbers of shared markers in each parent set. The number of shared markers determined genetic distance between parents.

We then examined the presence and absence of SNPs between all pairwise combinations of crosses. To determine if the difference in significant SNPs identified between crosses was the result of filtering and genotyping steps, we calculated the percentage of tags missing in all crosses. Tidyverse and dplyr R software packages were used to extract the number of tags before and after all the filtering steps, and at the 80x coverage threshold and to compare those to all tags genotyped in each cross (Wickham et al., 2019, 2020). To calculate the percentage of tags shared between each pair of crosses we compared the total significant tags genotyped in one cross to the sum of the significant tags genotyped in each focal pair. We also found whether significant tags were found to be insignificant or not genotyped in each cross. For tags with overlap in four or more crosses, we determined whether the allele in the cross was increasing or decreasing in frequency.

### Functional enrichment analysis

To investigate the functional implications of candidate tolerance loci, we mapped the 2bRAD reference (SRA accession: SRP066627, Howells et al., 2016) to the *P. daedalea* transcriptome (SRA accession PRJNA403854 Kirk et al., 2018) using Bowtie2 (Langmead et al., 2009) requiring a minimum MAPQ score of 23. Gene names and ontology terms were assigned to mapped 2bRAD tags based on the existing transcriptome annotations (Kirk et al., 2018). For tags that were identified in the transcriptome, we determined if the allele in the tag was increasing or decreasing in frequency. A rank-based gene ontology (GO) enrichment analysis was conducted to identify significantly enriched terms among annotated tags containing significant SNPs as identified in the allele frequency analysis using the GO_MWU suite of scripts (Wright et al., 2015), https://github.com/z0on/GO_MWU). The p-value from the allele frequency analysis was negative log-transformed and used as the continuous variable to identify GO terms enriched among the most significant SNPs using a Mann Whitney U test followed by Benjamini-Hochberg false discovery rate correction.

### Ocean Basin Comparison

As reads from our experiment were mapped against the Persian Gulf *P. daedalea* reference (SRA accession: SRP066627, Howells et al., 2016), we were able to compare tags containing significant SNPs identified in this study to the tags containing significant SNPs identified by Kirk and colleagues in their larval heat stress study (Kirk et al., 2018). This was done using dplyr and tidyverse software packages in R to extract all matching significant tags in this study and match them to the subset of tags containing significant SNPs from Kirk and colleagues’ study (Wickham et al., 2019, 2020).

## Results

### Survival in heat stress experiments

A Cox proportional hazards (CPH) model revealed significant effects of heat on larval survival (p-value < 0.0001) (Supplemental Figure. 2). Individual factors in the model, temperature and cross, as well as the interaction were significant (p-value < 0.0001). Endpoint survival probabilities ranged more among crosses at 35.5°C (0.405 to 0.750) than at 27°C (0.861 to 0.975) (Figure. 2). Cross DB had the greatest survival probability at 35.5°C (0.75+/- 0.04), while cross CA had the lowest survival probability (0.41 +/- 0.04) (Figure. 2). Coral E provided sperm in both EC and EF which both rank among the top four crosses, with (0.68+/- 0.04) and (0.64+/- 0.04) probability of survival, respectively. Coral C provided eggs in crosses DC and BC and provided sperm in cross CA, and this coral is associated with the bottom three survival probabilities 0.48 (+/- 0.04), 0.44 (+/- 0.05), and 0.40 (+/- 0.04).

**Figure 2.**
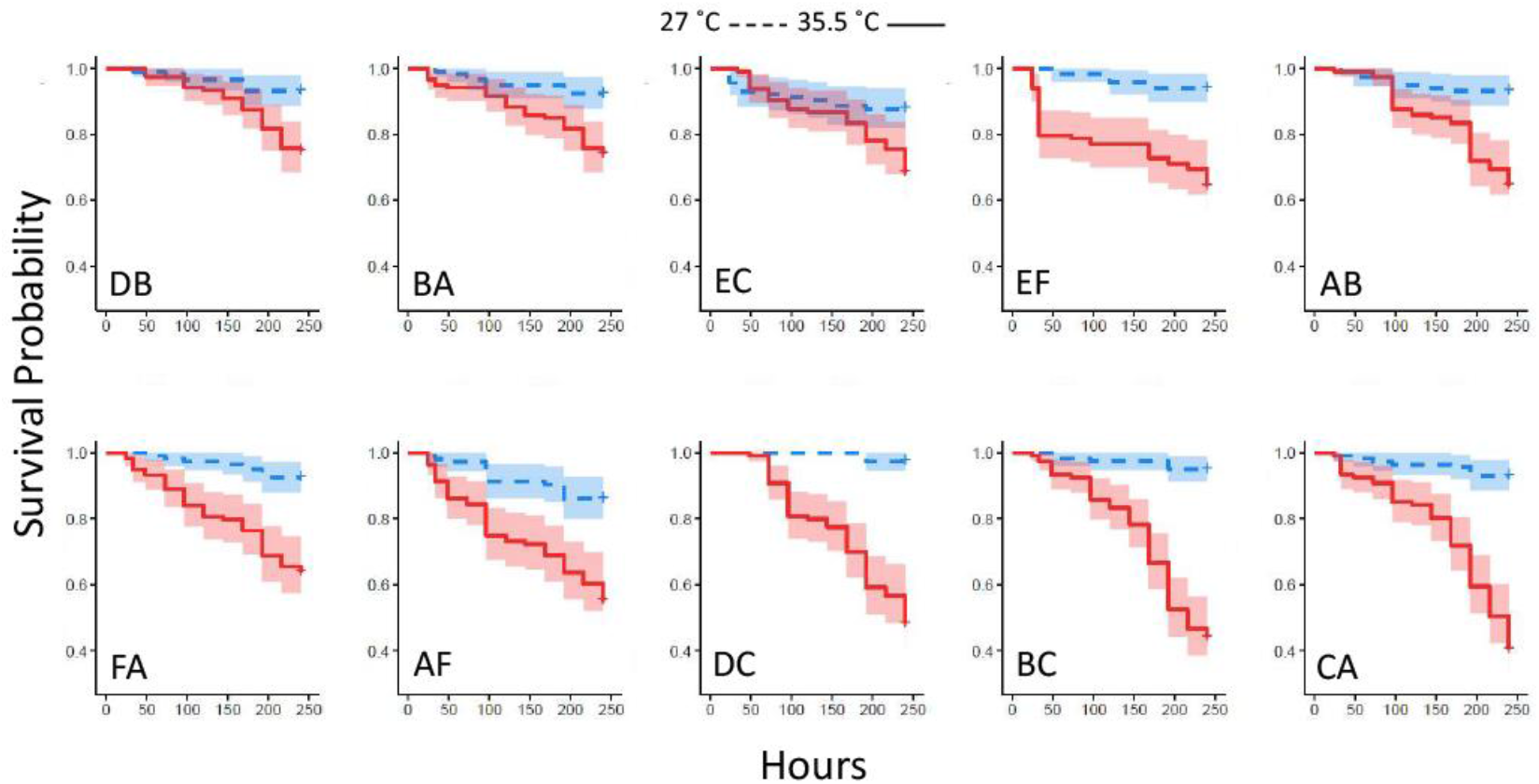
Heat tolerance differs among crosses. Kaplan Meier plots of cumulative probability of survival for each family in control 27.5°C (blue) and elevated 35.5°C (red) temperature. Panels are arranged in order of heat tolerance, based on survival probability at 240 hr. Shaded regions represent 95% confidence intervals.

CPH does not allow for testing of random effects. Since position of the net wells were randomly assigned, position could contribute to variation in thermal tolerance and thus we applied a generalized linear mixed model (GLMM) to test for position effects. We found that position did not contribute significantly to variation. Please see supplemental GLMM text for more details.

### Quantitative genetic analysis of variation in heat tolerance

Variation in survival during heat stress was highly heritable, with additive genetic variation explaining 66% of variation in survival (i.e. *h*^*2*^ = 0.66; Table 1). We also estimated *h*^*2*^ using the survival probabilities from the Kaplan Meier analysis and recovered a similar estimate (*h*^*2*^ = 0.64). Both metrics reflect the total mortality over the entire time course (Table 1).

**Table 1.** Estimates of narrow-sense heritability of thermal tolerance obtained from different survival metrics. Censored and uncensored KM estimates refer to cumulative probability of survival estimated with or without censoring individuals surviving at the experimental endpoint.

| *Data* | *Survival* | *h<sup>2</sup>* |
| --- | --- | --- |
| Survival at 240hr | 59% | 0.66 |
| KM Estimate | 59% | 0.64 |

### Genomic analysis of markers under selection during heat stress

Comparative analysis of larvae surviving an acute heat stress exposure versus paired control larval pools for each cross revealed that experimental treatment altered allele frequencies at hundreds of loci. Logistic regression of multilocus SNP genotypes (Figure. 3) uncovered tens-to-hundreds of alleles in each cross whose frequencies were significantly altered by heat stress (Table 2). Overall, there was a general pattern of increase in major allele frequency in most crosses except for crosses BC, DC, and FA (Figure. 4). This also meant that minor alleles were decreasing in frequency in most crosses. The number of alleles significantly associated with survival under heat stress differed widely among crosses; however, the number of significant alleles was not correlated with the number of tags genotyped in each sample (*P* = 0.48, *R*^*2*^ = - 0.26), genetic distance between the parents (*P* = 0.46, *R*^*2*^ = -0.27), or the heat tolerance of each cross (*P* = 0.48, *R*^*2*^ = -0.27). The variation was also not associated with a particular parent. For example, parents C and D each produced crosses with few SNPs under selection (BC and DB), but the combination of these parents in cross DC had the most, with 336 SNPs associated with survival under heat stress (Table 2).

**Table 2.**
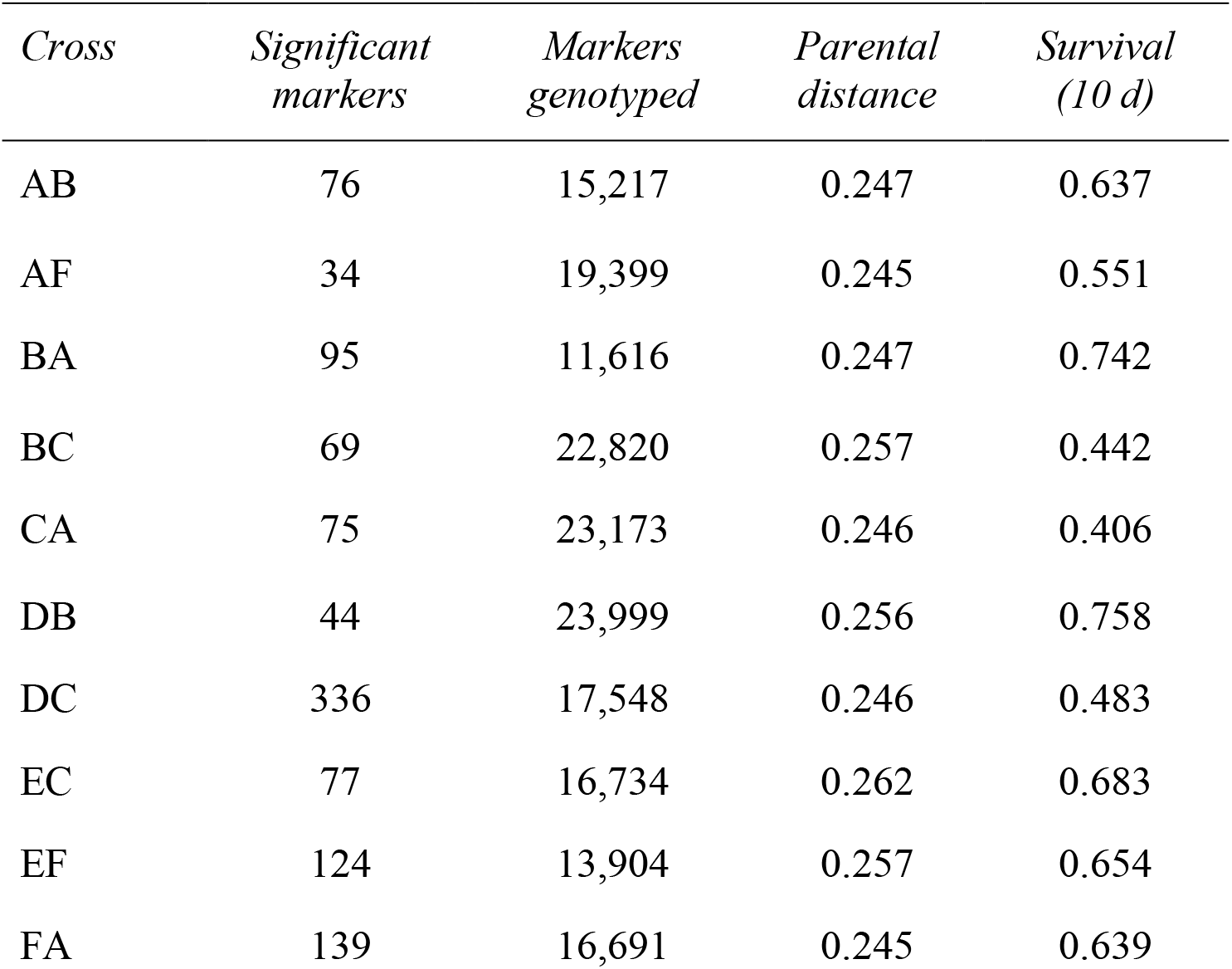
The number of loci at which allele frequencies were significantly altered in survivors of thermal stress in each cross.

To evaluate the generalizability of genetic associations we searched for overlap in significant tags among crosses. A proportion of tags were consistently associated with heat stress across multiple crosses. In total, 148 significant tags were found in more than one cross (Table 3 and Supplemental Table 2). A pairwise comparison of the percent of shared significant tags between a focal pair of crosses, revealed that 1-6% of significant tags were found to be significant in another cross (Figure. 5, Table 3, and Supplemental Table 2). The same two tags were found in seven crosses, a single tag overlapped in six crosses, and the same four tags overlapped in five crosses (Table 3). The major allele in six different tags that were found to be significantly changing in frequency in four or more crosses was found to increase in frequency (Table 3). Five tags showed mixed change in frequency for the major allele, in some crosses it increased and in others it decreased (Table 3). Only one tag showed consistent decrease in frequency of the major allele across crosses in which it was identified (Table 3). However, most tags associated with heat tolerance in one cross were not associated with heat tolerance in unrelated crosses. For example, cross DC shared no parents with crosses EF or FA and 82 - 92% of the tags identified in each cross were not associated with heat stress in the other two.

**Table 3.**
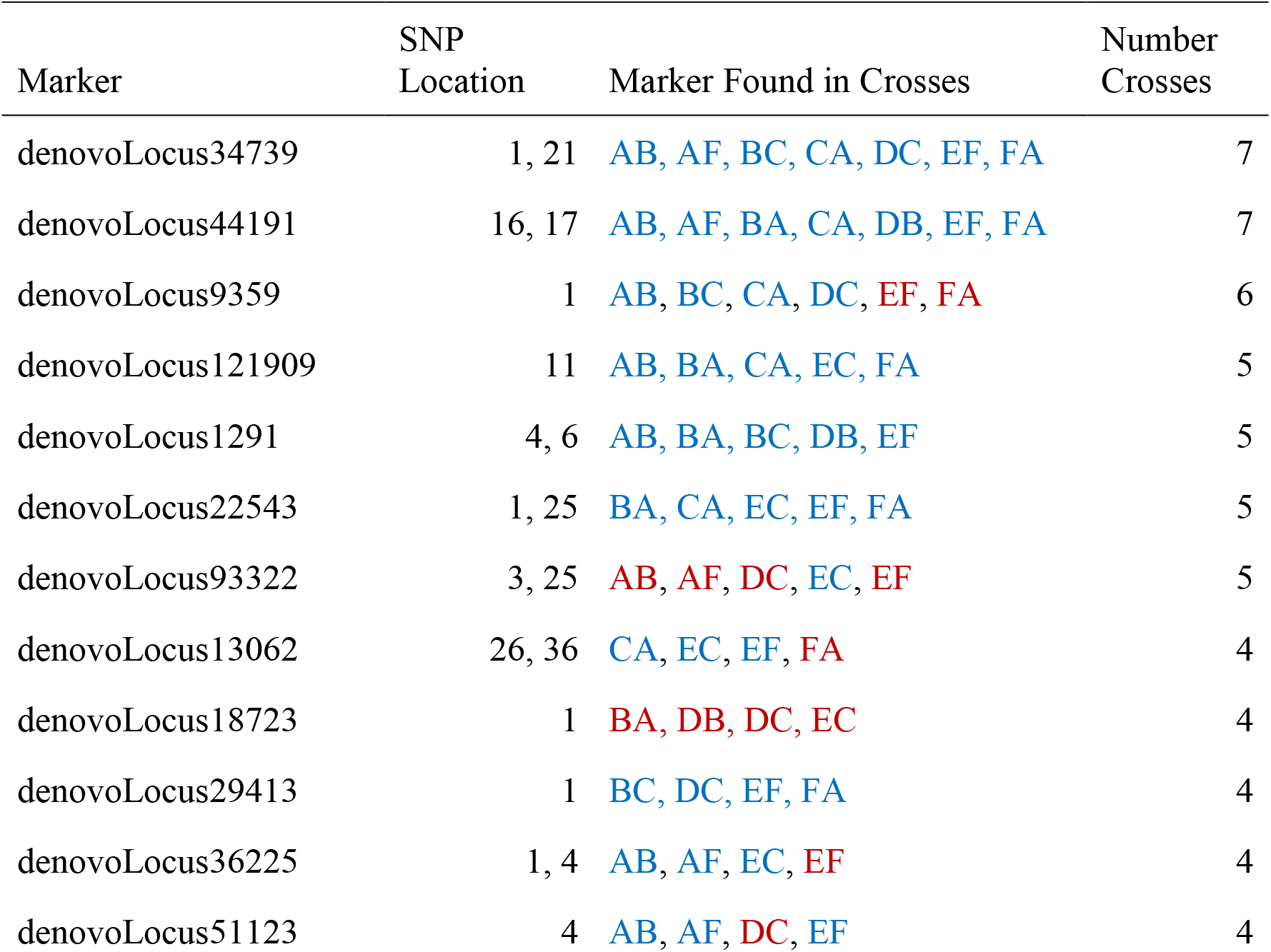
Tags identified as changing in frequency after heat stress in four or more crosses. The color of the cross indicates whether the allele in the tag was increasing in frequency (blue) or decreasing in (red).

| Marker | SNP Location | Marker Found in Crosses | Number Crosses |
| --- | --- | --- | --- |
| denovoLocus34739 | 1, 21 | AB, AF, BC, CA, DC, EF, FA | 7 |
| denovoLocus44191 | 16, 17 | AB, AF, BA, CA, DB, EF, FA | 7 |
| denovoLocus9359 | 1 | AB, BC, CA, DC, EF, FA | 6 |
| denovoLocus121909 | 11 | AB, BA, CA, EC, FA | 5 |
| denovoLocus1291 | 4, 6 | AB, BA, BC, DB, EF | 5 |
| denovoLocus22543 | 1, 25 | BA, CA, EC, EF, FA | 5 |
| denovoLocus93322 | 3, 25 | AB, AF, DC, EC, EF | 5 |
| denovoLocus13062 | 26, 36 | CA, EC, EF, FA | 4 |
| denovoLocus18723 | 1 | BA, DB, DC, EC | 4 |
| denovoLocus29413 | 1 | BC, DC, EF, FA | 4 |
| denovoLocus36225 | 1, 4 | AB, AF, EC, EF | 4 |
| denovoLocus51123 | 4 | AB, AF, DC, EF | 4 |

**Figure 3.**
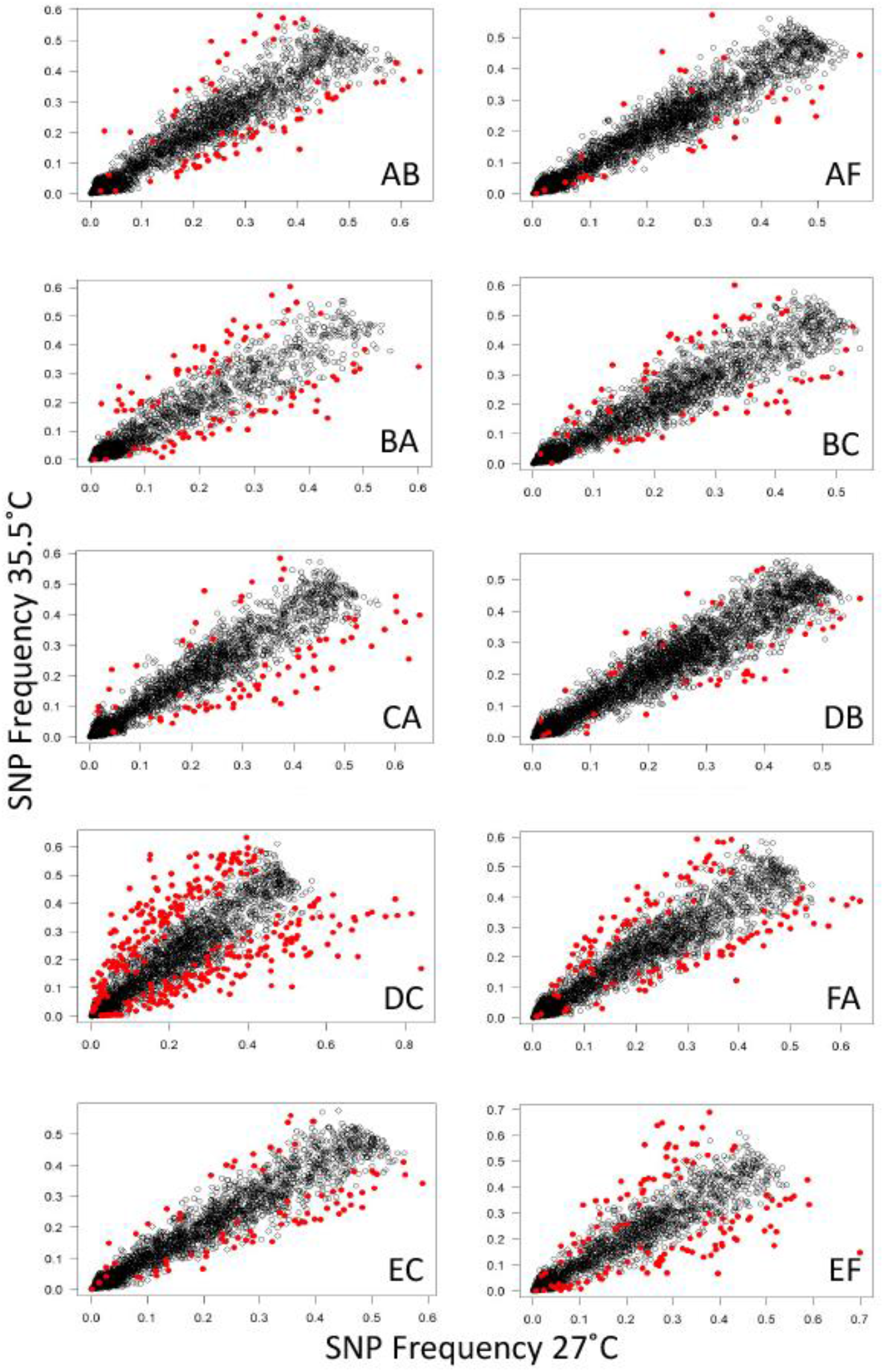
Heat stress alters allele frequencies in surviving larvae. Panels show correlations between the average allele frequencies in survivors of replicate heat stress treatments relative to the corresponding controls. Markers with significantly altered allele frequencies (logistic regression, FDR < 0.05) are highlighted in red.

**Figure 4.**
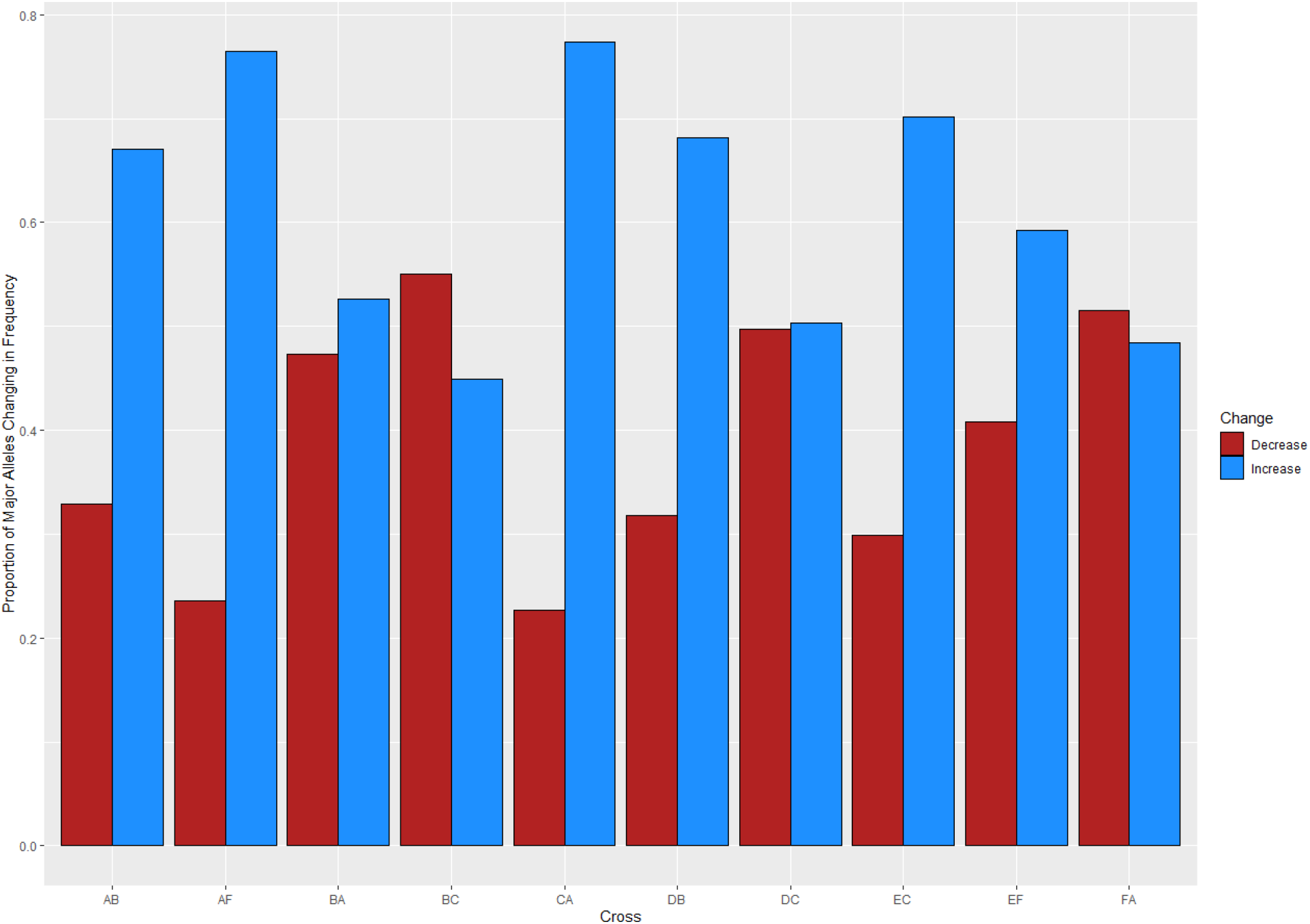
The proportion of major alleles that changed in frequency after heat stress. Crosses are indicated on the x-axis and the proportions of major alleles that increased (blue) and decreased (red) are depicted on the y-axis.

**Figure 5.**
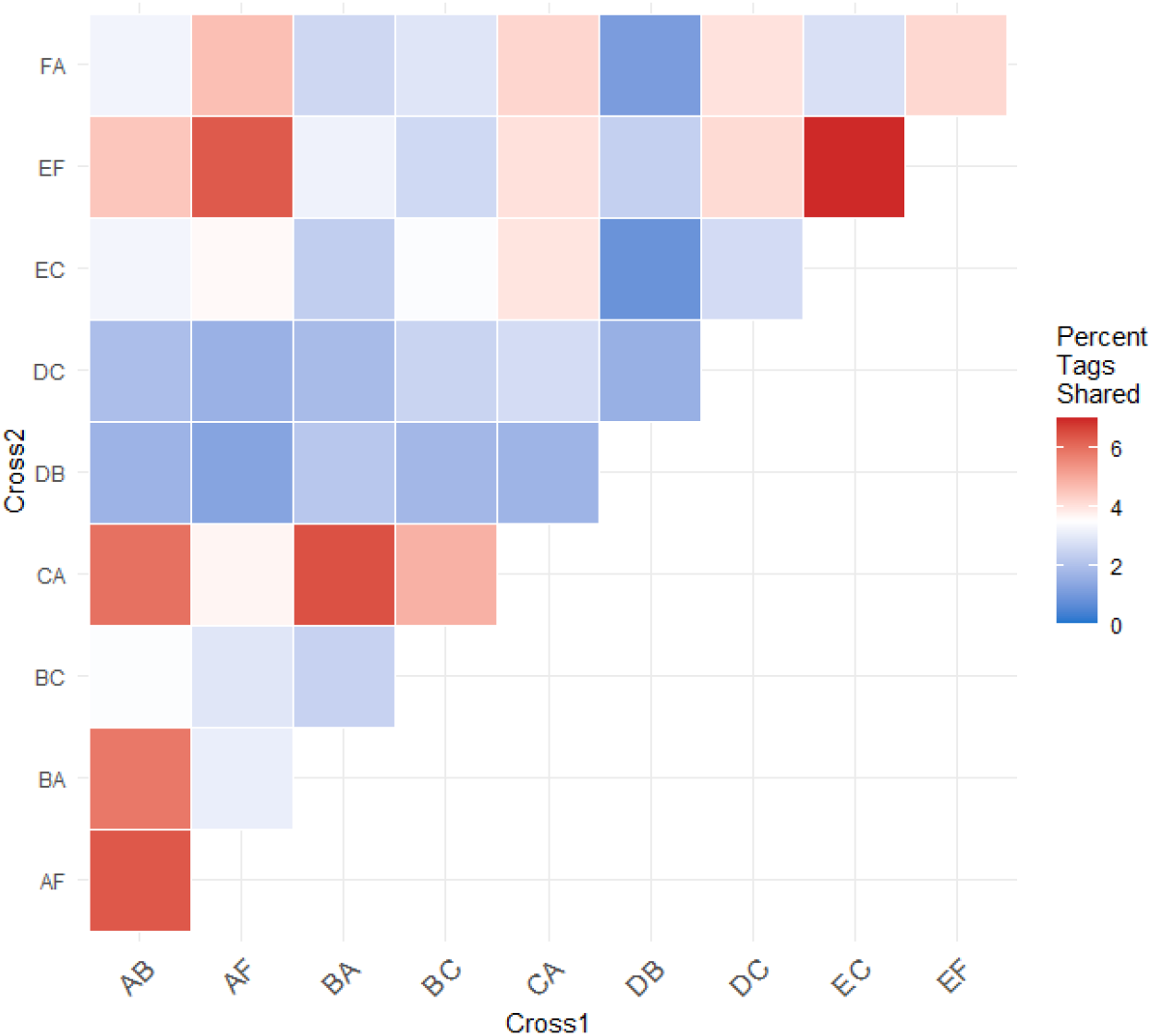
Heatmap of the percent of significant tags shared between pairs of crosses. The percent of shared significant tags for each cross was calculated from the number of significant tags shared between the pair and the total number of potential tags that could have been shared. These comparisons demonstrate independent associations between heat stress and genotype at these loci across different genetic backgrounds.

To determine whether this lack of overlap was a result of the necessarily stringent filtering required for the analysis of pool-seq data (Kofler et al., 2011; Zhu et al., 2012), we investigated overlap in tags among crosses at earlier stages of our analysis. After initial quality filtering, we found only 30 – 63% of tags were genotyped in another cross (median of 52%, Supplemental Table 3). This range did not appreciably change after the 80X coverage threshold was applied, with 21 - 66% of tags identified in other crosses (median of 49%, Supplemental Table 4), indicating that the coverage threshold was not driving this pattern. However, statistical thresholds did have a moderate impact, as the percentage of significant tags in one focal cross not identified in other pairwise comparisons ranged from 11 - 61% (median of 42%, Supplemental Table 5).

We identified whether each significant tag was either not genotyped or was genotyped and found to be insignificant. We found that a majority of significant tags (64%) were genotyped in at least five out of ten crosses (Supplemental Table 7). Of the tags that were found to be significant in at least four or more crosses, they were found to be insignificant in at least one and in most cases more than one other cross (Table 4, Supplemental Table 6).

**Table 4.** The number of crosses a tag was identified as either significant, insignificant, or not genotyped for tags that were found to be significantly changing in frequency in four or more crosses.

| *Tag* | *Number Crosses Significant* | *Number Crosses Insignificant* | *Number Crosses Not Genotyped* |
| --- | --- | --- | --- |
| denovoLocus34739 | 7 | 1 | 2 |
| denovoLocus44191 | 7 | 0 | 3 |
| denovoLocus9359 | 6 | 3 | 1 |
| denovoLocus121909 | 5 | 4 | 1 |
| denovoLocus22543 | 5 | 3 | 2 |
| denovoLocus93322 | 5 | 3 | 2 |
| denovoLocus1291 | 5 | 1 | 4 |
| denovoLocus18723 | 4 | 5 | 1 |
| denovoLocus13062 | 4 | 4 | 2 |
| denovoLocus36225 | 4 | 4 | 2 |
| denovoLocus29413 | 4 | 3 | 3 |
| denovoLocus51123 | 4 | 1 | 5 |

### Functional significance of enriched tags

Seven GO terms were significantly enriched among the nine annotated tags that changed in allele frequency in response to heat stress in any cross (Figure 6). Tags in genes regulating cell projection, receptor signaling, regulation of apoptosis, regulation of hydrolase, and negative regulation of metabolism were over-represented among the most significantly differentially represented tags under heat stress conditions (Table 5). For the subset of significant tags shared among crosses, 9 markers could be unambiguously assigned to a position in the transcriptome all of which were annotated (Table 5). Furthermore, two of these tags which were identified in three crosses were annotated as genes previously implicated in heat tolerance Monocarboxylate transporter 10 (K1PRR1) and Isocitrate dehydrogenase subunit 1 (E9C6G6) (Table 5) (Drury et al., 2021; Kenkel et al., 2013; Pathmanathan et al., 2021; Polato et al., 2010). In two tags that mapped to transcripts and overlapped in other crosses, we found consistent decrease in the major allele for the Monocarboxylate transporter and consistent increase of the major allele for the isocitrate dehydrogenase subunit (Table 5).

**Figure 6.**
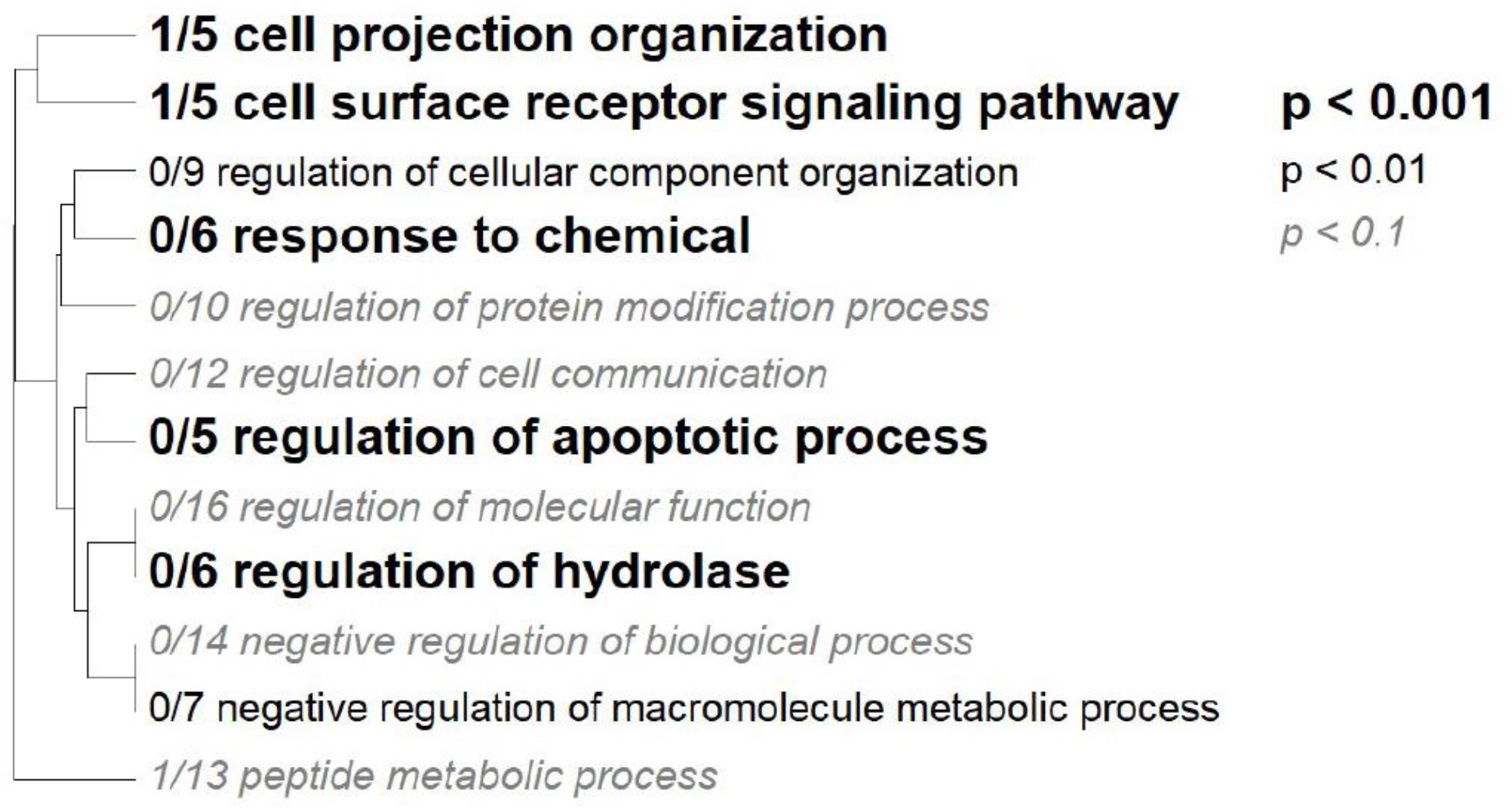
Gene Ontology Tree. Gene ontology categories enriched in significant markers. The size of the font indicates the significance of the category per the inset key. Markers were mapped to the *Platygyra daedalea* transcriptome and the matching transcripts were assigned the log adjusted p-value from the allele frequency logistic regression.

**Table 5.** Tags mapped to transcripts that changed in frequency after heat stress. Listed in order of the number of crosses in which the tag was found. Color of the cross indicates the direction in the change of frequency: red represents a decrease and blue represents an increase.

| *Crosses* | *Number of*<br>*Tags Mapped*<br>*to Transcript* | *Transcripts* | *Descriptions (Annotation)* |
| --- | --- | --- | --- |
| AB, BA, EF | 1 | c39807 | Monocarboxylate transporter 10 (K1PRR1) |
| AF, BA, CA | 1 | c15457 | Isocitrate dehydrogenase subunit 1 (E9C6G6) |
| AB | 1 | c36043 | Metalloendopeptidase (A7RVF7) |
| AB | 1 | c9426 | 60S_ribosomal_protein_L6_(Fragment) |
| BC | 1 | c10517 | 3-oxoacyl-[acyl-carrier-protein] reductase (K1QNK4) |
| DC | 1 | c7510 | Putative dna polymerase type b (A0A069DZN0) |
| DC | 1 | c11010 | Zinc finger MYM-type protein 3 (K1QD06) |
| FA | 1 | c16893 | Rab9_effector_protein_with_kelch_motifs |
| FA | 1 | c12147 | GPI mannosyltransferase 2 (K1PUQ8) |

### Ocean basin comparison

Finally, we compared significant SNPs arising from any cross with previously identified SNPs enriched in response to heat stress in Persian Gulf *P. daedalea* (Kirk et al., 2018). We found two tags significantly associated with both survival and heat tolerance in our dataset overlapped with sites identified in (Kirk et al., 2018) (Table 6). The specific SNPs identified in each tag differed among studies (Table 5) and the tags did not match any genes in the reference transcriptome, so the potential functional significance of this shared variation remains unknown.

**Table 6.** Tags shared between the Persian Gulf and GBR populations that are significant to thermal tolerance.

| Cross | Tag | SNP Base<br>Pair GBR | SNP Base<br>Pair Persian<br>Gulf | Major Allele<br>GBR | Major Allele<br>Persian Gulf |
| --- | --- | --- | --- | --- | --- |
| CA | denovoLocus12624 | 35 | 28 | T | GT |
| DC | denovoLocus60661 | 16 | 21 | T | C |

## Discussion

### Adaptive potential of Platygyra daedalea

Our analysis demonstrates substantial variation in heat tolerance is evident among larval crosses. Moreover, 66% of this variation was attributable to additive genetic effects, indicating significant adaptive potential. Coral parent E was generally associated with crosses that had high survival probabilities and Coral parent C was associated with crosses that had lower survival probabilities (Figure 2). Our results add to only four published narrow-sense heritability estimates for heat tolerance in corals, all of which find this trait is highly heritable ranging from 0.58 – 0.87 (Bairos-Novak et al., 2021; Dixon et al., 2015; Dziedzic et al., 2019; Kirk et al., 2018; Quigley et al., 2020). Heritability estimates from different populations can yield valuable information about selection in a given geographic region as estimates are dependent on the environment and the standing genetic variation in the population for which the estimate is made (Falconer & Mackay, 1996). Heritability estimates will be low if all parents express the same phenotype and genetic variation contributing to a trait is low (Falconer & Mackay, 1996). However, they will be high if there is phenotypic variation and genetic variation contributing to the trait is also high (Falconer and Mackay, 1996). A novel comparison is the contrast of the heritability estimate for the same phenotype, larval survival in acute heat stress (0.48) found by Kirk and colleagues for *P. daedalea* in the Persian Gulf (Kirk et al., 2018). Persian Gulf *P. daedalea* have survived in a very shallow and warm sea for thousands of years (Purkis & Riegl, 2012), which may mean that a degree of selection for heat tolerance has already taken place resulting in less genetic diversity in the extant population (Falconer & Mackay, 1996). The lower heritability value is evidence to support this and the comparatively higher estimate of heritability for this trait in the central GBR *P. daedalea* population suggests that until very recently, heat stress may not have been the predominant selective pressure on this population, consistent with historical bleaching datasets for this region (Hughes et al., 2018).

Responses to selection depend on genetic variation in traits (Falconer & Mackay, 1996; Lynch & Walsh, 1998). Consequently, our study contributes to understanding the quantitative genetic framework necessary to model heat tolerance under various selection regimes. In its simplest form, this can be expressed in the breeder’s equation, which describes the response to selection as the product of the selection differential (S) and heritability of the trait under selection. Selection differentials have not been widely estimated in corals (Kenkel et al., 2015; Weeriyanun et al., 2021), but considering the widespread reductions in coral populations following coral bleaching (Bellwood et al., 2004; Bruno & Selig, 2007; De’ath et al., 2012; Hoegh-Guldberg et al., 2007) they are likely high for sensitive species. For truncation selection, where all individuals with trait values above a threshold contribute offspring to the next generation, selection differentials are equal to the difference in mean trait values between the selected population and the original population (Falconer & Mackay, 1996; Lynch & Walsh, 1998) expressed as standard deviations of the population mean. If the selection differential equals one standard deviation, all parents with traits one standard deviation above the population mean contribute to phenotypic variation in the next generation. Based on the distribution of cumulative survival probabilities in heat stress from our experiment (0.57 ± 0.18) and the heritability of variation in heat tolerance we estimated from these data (0.66), the Breeder’s Equation predicts a response to selection of 0.12. We would expect that if larvae of the next generation after selection were subjected to the same heat stress treatment, the average mortality would be 0.48 (0.57 - 0.12), a 20% reduction in mortality under acute heat stress in a single generation. If instead of the high *h*^*2*^ estimated in our study (0.66), we modeled the responses to selection using a minimum *h*^*2*^ value of 0.1, we would predict only a 3% reduction in mortality in the following generation. While the estimates above are underpinned by many assumptions, the comparison highlights how empirical estimations of narrow-sense heritability can be used to predict adaptive responses. The relevance of these estimates for population responses depends on the extent to which larval survival during heat stress is directly under selection in natural populations, or alternatively the effectiveness of larval heat tolerance as a proxy for heat tolerance in adult colonies. Considering the lack of work in multi-generational comparisons, this would be a fruitful direction of future study.

### Allele frequency analysis supports multilocus adaptation

Physiological and fitness traits can be determined by single or few loci of large effect, but more commonly they are underpinned by many loci of small effect (Sella & Barton, 2019). Surveying a larger portion of the population and limiting our detection of allele frequency to within the offspring of two known parents has helped to detect tens-to-hundreds of alleles supporting survival during heat stress. Comparing between crosses allowed us to investigate the generality of these patterns among different genetic backgrounds. This comparison revealed a number of markers with consistent changes in allele frequencies in multiple crosses (Table 3 and Supplemental Table 2). The tags that show allele frequency change in a consistent direction are important targets for future studies of markers of thermal tolerance. Those that show inconsistent direction of frequency change could be further evidence of the polygenicity of this trait. One allele might be important for thermal tolerance response, but if the right combination of alleles are not found at other sites in the genome thermal tolerance will not be as high.

However, many of the associations detected in each cross were not shared in comparisons of multiple crosses. Our investigation into the lack of overlap among markers revealed that 49% of markers in a given cross on average were not genotyped in other crosses even before the application of the 80X within cross coverage filter (Supplementary tables 3-5), suggesting that this loss is not a result of bioinformatic filtering applied but instead may result from aspects of the sequencing methodology and the biology of the animals involved. Null alleles are one common explanation for missing markers during sequencing (e.g. Crooks et al., 2013). Alternatively, this bias could also result from the pooled sequencing approach. There were 100 individuals in each of eight replicate pools. Although amplification of libraries was kept to a minimum (no more than 17 cycles of PCR) a certain amount of PCR bias is unavoidable and in a given cross certain regions may have been amplified more frequently (Dohm et al., 2008). A small number of additional sites were lost through the filtering process, but stringent thresholds are required to reduce false positive allele calls given the small size of our RAD tags (Kofler et al., 2011). In this study, a pooled sequencing approach was chosen to maximize the resources available to survey as much of the population as possible. However, this is a clear tradeoff of pooled sequencing with this number of individuals per pool and sequencing relatively short fragments of the genome. Sequencing individual larvae may help alleviate this issue (e.g. Dixon et al. 2015).

Statistical thresholds also had an impact and the observation that certain SNPs were found to be significant in some crosses but not others could also result from the underlying distribution of those alleles in our parental population. In isolated populations the loci contributing to a trait are predicted to be fewer and of larger effect size (Savolainen et al., 2013). However, in populations experiencing immigration, the locus effect size and the number of loci that contribute to a trait become more variable (Savolainen et al., 2013). Corals can disperse in their larval phase before metamorphosis into a sedentary juvenile, so it is entirely possible that individuals originating from disparate reefs were among those chosen for our parental population (Davies et al., 2015). The observation that many markers associated with heat stress were specific to individual crosses suggests that many alleles of small effect contribute to variation in heat tolerance and that many different combinations of these small effect alleles can affect heat tolerance as is the growing consensus for thermal tolerance in coral (Bay & Palumbi, 2014; Fuller et al., 2020; Kirk et al., 2018; Thomas & Palumbi, 2017). The fact that many of our significant tags were found to be insignificant in other crosses also supports this (Table 4, Supplemental Table 7). These differences may also result in part from epistatic interactions where the effects of variation at these loci depend on the interaction of two or more genes and composition of alleles at these genes which is dependent on genetic background (Day et al., 2008). The large number of alleles changing in frequency in cross DC could result from an unfortunate combination of parental alleles at many loci producing larvae with multiple deleterious alleles. This could be the result of additive genetic effects, but it could also be an indicator of epistasis, which is harder to detect but found frequently in polygenic traits in model organisms (Soyk et al., 2020). Nevertheless, repeated observation of some of these associations across different genetic backgrounds provides strong evidence that these markers are linked to genetic factors contributing to variation in heat tolerance.

Future studies can use the alleles that increased in frequency and the genomic regions identified in this and other studies to develop biomarkers to identify resilient coral populations for assisted gene flow or use in selective breeding programs. Starting with genetic markers associated with habitat variation in previous studies (Capper et al., 2015), we can test these associations through a combination of laboratory experiments and extensive population surveys of allele frequencies (Jin et al., 2016). These studies revealed that the frequencies of alleles associated with enhanced tolerance for environmental stress range from 0.48 in some populations to 0.92 in others (Jin et al., 2016). Further studies are needed to extend these surveys across species, markers, and geographic regions to develop an ecosystem-scale perspective on the potential for coral adaptive responses to ocean warming.

### SNPs associated with heat tolerance occur in genes related to extracellular matrix and metabolism

Among the annotated SNPs exhibiting significant changes in allele frequency in response to experimental selection, a number of functional enrichments were identified in genes associated with the organization of the cellular matrix and cellular components, the regulation of apoptosis, regulation of hydrolase, and metabolism. In addition, we found a subset of tags mapped to genes previously implicated in heat tolerance, two of which were observed in multiple crosses (Table 5). These genes are prime candidates for further functional studies aimed at delineating the potential mechanistic role of these polymorphisms (Cleves et al., n.d., 2018, 2020).

Four markers were associated with genes that play roles in the extracellular matrix, with enrichments in cell projection organization, surface receptor signaling, and cell component organization (Table 5, Figure. 6). GPI Mannosyltransferase 2 functions in the assembly of glycosylphosphatidylinositol (GPI) anchors that attach proteins to cell membranes (Ji et al., 2005). In corals this protein is in the pathway that changes the mannose content of the extracellular matrix (Lee et al., 2016). The abundance of Gammaproteobacteria is reduced with the change in mannose in coral mucus (Lee et al., 2016) Metalloendopeptidase, a metal-dependent protease, and Zinc finger MYM-type protein both restructure and organize the extracellular matrix (Lodish et al., 2020). The Rab9 effector with kelch motifs controls docking of endosomes in the process of exocytosis (Díaz et al., 1997). While metalloendopeptidase and this zinc finger protein were not found in other coral studies, they are found in heat stress in other invertebrates (Juárez et al., 2021; Prieto et al., 2019; Zhang et al., 2015). On the other hand, Rab proteins have been identified as important for signaling the start of phagocytosis mechanisms and cell destruction in anemones during oxidative stress and bleaching (Chen et al., 2005; Downs et al., 2009). In this case it is likely the Rab 9 effector is involved in signaling the start of the cell death during heat stress. SNPs in these genes indicate that control of cell reorganization and structure is important during heat stress.

Negative regulation of macromolecule metabolic process was also identified as a significant GO category (Figure 6). Three coral crosses had a SNP in the Krebs cycle enzyme isocitrate dehydrogenase (IDH) and a monocarboxylate transporter both of which are associated with this annotation. These proteins have been identified in gene expression studies as important to the coral thermal stress response (Drury et al., 2021; Kenkel et al., 2013; Pathmanathan et al., 2021; Polato et al., 2010). IDH participates in reducing reactive oxygen species and is one of the 44 proteins identified as the minimal stress proteome (Kultz, 2005). The monocarboxylate transporter moves lactate, pyruvate, and ketone bodies across plasma membranes, thus it is associated with the regulation of metabolism (Halestrap, 2012). This transporter has been identified as important in recent coral heat stress studies and is upregulated under heat stress (Drury et al., 2021; Pathmanathan et al., 2021). The SNPs in IDH and the monocarboxylate transporter associated with heat tolerance in this study further strengthens their proposed role in the thermal stress response in corals.

Additional genes annotated with GO categories like metabolism, regulation of hydrolase, and regulation of apoptosis, may also be important for handling reactive oxygen species. While the 3oxoacyl (acyl-carrier-protein) reductase has a larger role in fatty acid biosynthesis, it also catalyzes a reduction reaction (Chan & Vogel, 2010). Similarly, IDH is responsible for regulation of metabolism, but the reaction it catalyzes is also a reduction. These genes have been identified as important for controlling reactive oxygen species levels resulting from increased metabolism during heat stress (Dixon et al., 2015) and our results add further evidence of the possible importance of these SNPs to corals’ heat stress response.

The consistent change in the frequency of the major alleles in the monocarboxylate transporter and isocitrate dehydrogenase across multiple crosses also suggests that these alleles are important for heat stress survival. They are prime targets for future studies assessing markers of thermal tolerance.

### Ocean Basin Comparison

Comparisons of populations of *P. daedalea* in more typical reef temperature conditions (this study) and unusually warm conditions (Kirk et al., 2018) will be important to understand the genomic basis of heat tolerance. Two markers from the present study were significantly associated with heat tolerance in the Persian Gulf population (Table 6). Neither of these markers matched genes in the transcriptome and without a genomic resource it is not possible to identify their location in the genome or proximal genes. However, their repeated identification strongly supports an important role in coral thermal tolerance. Further work could include screening for differential abundance of these alleles among natural thermal gradients and potential validation for gene function through knock-down from CRISPR-Cas9 induced mutagenesis (e.g., Cleves et al., 2018). The lack of shared markers between these cross-ocean basin populations may be further evidence that many alleles of small effect contribute to heat tolerance and that they vary in their distribution across populations.

### Conclusions

Our findings demonstrate substantial genetic variation and high heritability of heat tolerance (i.e., > 60%) in the coral *Platygyra daedalea* from the GBR supporting the potential for adaptation. Our analysis identified hundreds of markers associated with heat tolerance, including markers that were repeatedly associated with heat tolerance across different genetic backgrounds. This supports the conclusion that heat tolerance is a complex polygenic trait determined by many alleles of small effect that are likely to be species and population specific. Despite this complexity, we nevertheless found markers shared in multiple crosses that may be important for heat tolerance in *P. daedalea*. Two markers were also identified in Persian Gulf populations, and a number occurred in genes previously implicated in coral thermal tolerance. Our findings build on the growing body of evidence that natural populations of corals harbor substantial genetic variation in heat tolerance that can support adaptive responses to selection and identify genetic markers for heat tolerance in a globally distributed coral species. These heat tolerance markers are essential to any type of reseeding or assisted gene flow and assisted migration programs, which may be necessary to maintaining reefs in the future as we work to reduce CO2 emissions and mitigate the effects of climate change.

## Acknowledgements

We thank the staff of the National Sea Simulator at the Australian Institute of Marine Science for their assistance in the collection and husbandry of the corals used in these experiments. We also thank Ida Bjornsbo for her tireless assistance in counting coral larvae at all hours of the day and night. This research was conducted under Great Barrier Reef Marine Park Authority permit (G11/34671.1) and supported with funding from the National Science Foundation Graduate Research Fellowship to Holland Elder, National Science Foundation Graduate Research Opportunities Worldwide to Holland Elder and internal grants from AIMS to Line K. Bay.

## Code and Data Availability

Sequence Access: NCBI, Accession: PRJNA542930 https://github.com/hollandelder/Platygyradaedalea

_HeritabiltyandMarkersofThermalTolerance https://github.com/Eli-Meyer/sequence_processing https://github.com/z0on/GO_MWU)

## Authors’ contributions

E.M., H.E., and L.B. conceived the experiments. H.E., V.M., J.M., and L.B. conducted experiments, L.B. and A.B. provided advice and logistical support through the project, H.E. conducted genetic and statistical analysis and wrote the manuscript. L.B. and E.M. helped with genetic and statistical analysis and L.B., V.W. and A. B. helped write the manuscript.

## References

Bairos-Novak, K. R., Hoogenboom, M. O., van Oppen, M. J. H., & Connolly, S. R. (2021). Coral adaptation to climate change: Meta-analysis reveals high heritability across multiple traits. In Global Change Biology (Vol. 27, Issue 22, pp. 5694–5710). John Wiley and Sons Inc. https://doi.org/10.1111/gcb.15829

Barshis, D. J., Ladner, J. T., Oliver, T. a, Seneca, F. O., Traylor-Knowles, N., & Palumbi, S. R. (2013). Genomic basis for coral resilience to climate change. Proceedings of the National Academy of Sciences of the United States of America, 110(4), 1387–1392. https://doi.org/10.1073/pnas.1210224110

Baums, I. B., Baker, A. C., Davies, S. W., Grottoli, A. G., Kenkel, C. D., Kitchen, S. A., Kuffner, I. B., LaJeunesse, T. C., Matz, M. v., Miller, M. W., Parkinson, J. E., & Shantz, A. A. (2019). Considerations for maximizing the adaptive potential of restored coral populations in the western Atlantic. Ecological Applications, 29(8), 1–23. https://doi.org/10.1002/eap.1978

Bay, L. K., Doyle, J., Logan, M., & Berkelmans, R. (2016). Recovery from bleaching is mediated by threshold densities of background thermo-tolerant symbiont types in a reef-building coral. Royal Society Open Science, 3(160322).

Bay, R. a, & Palumbi, S. R. (2014). Report Multilocus Adaptation Associated with Heat Resistance in Reef-Building Corals. Current Biology, 1–5. https://doi.org/10.1016/j.cub.2014.10.044

Bay, R., Rose, N., Logan, C., & Palumbi, S. (2017). Genomic models predict successful coral adaptation if future ocean warming rates are reduced. Science Advances, 1–10. https://doi.org/10.1126/sciadv.1701413

Bellwood, D. R., Hughes, T. P., Folke, C., & Nystro, M. (2004). Confronting the coral reef crisis. Nature, 429(June), 827–833.

Berkelmans, R. (2002). Time-integrated thermal bleaching thresholds of reefs and their variation on the Great Barrier Reef. Marine Ecology Progress Series, 229, 73–82.

Berkelmans, R., & van Oppen, M. J. H. (2006). The role of zooxanthellae in the thermal tolerance of corals: a “nugget of hope” for coral reefs in an era of climate change. Proceedings. Biological Sciences / The Royal Society, 273(1599), 2305–2312. https://doi.org/10.1098/rspb.2006.3567

Bruno, J. F., & Selig, E. R. (2007). Regional Decline of Coral Cover in the Indo-Pacific : Timing, Extent, and Subregional Comparisons. PloS One, 8. https://doi.org/10.1371/journal.pone.0000711

Capper, R. L., Jin, Y. K., Lundgren, P. B., Peplow, L. M., Matz, M. v, & Oppen, M. J. H. van. (2015). Quantitative high resolution melting : two methods to determine SNP allele frequencies from pooled samples. BMC Genetics, 16(62), 1–13. https://doi.org/10.1186/s12863-015-0222-z

Chan, D. I., & Vogel, H. J. (2010). Current understanding of fatty acid biosynthesis and the acyl carrier protein. In Biochemical Journal (Vol. 430, Issue 1, pp. 1–19). https://doi.org/10.1042/BJ20100462

Charmantier, A., & Garant, D. (2005). Environmental quality and evolutionary potential : lessons from wild populations. Proceedings of the Royal Society, 272, 1415–1425. https://doi.org/10.1098/rspb.2005.3117

Chen, M. C., Hong, M. C., Huang, Y. sen, Liu, M. C., Cheng, Y. M., & Fang, L. S. (2005). ApRab11, a cnidarian homologue of the recycling regulatory protein Rab11, is involved in the establishment and maintenance of the Aiptasia-Symbiodinium endosymbiosis. Biochemical and Biophysical Research Communications, 338(3), 1607–1616. https://doi.org/10.1016/j.bbrc.2005.10.133

Cleves, P. A., Shumaker, A., Lee, J. M., Putnam, H. M., & Bhattacharya, D. (2020). Unknown to Known: Advancing Knowledge of Coral Gene Function. In Trends in Genetics (Vol. 36, Issue 2, pp. 93–104). Elsevier Ltd. https://doi.org/10.1016/j.tig.2019.11.001

Cleves, P. A., Strader, M. E., Bay, L. K., Pringle, J. R., & Matz, M. v. (2018). CRISPR/Cas9-mediated genome editing in a reef building coral. PNAS, 115(20), 5235–5240. https://doi.org/10.1073/pnas.1722151115

Cleves, P. A., Tinoco, A. I., Bradford, J., Perrin, D., Bay, L. K., & Pringle, J.R. (n.d.). Reduced thermal tolerance in a coral carrying CRISPR-induced mutations in the gene for a heat-shock transcription factor. https://doi.org/10.1073/pnas.1920779117/-/DCSupplemental

Coles, S. L., & Riegl, B. M. (2013). Thermal tolerances of reef corals in the Gulf : A review of the potential for increasing coral survival and adaptation to climate change through assisted translocation. Marine Pollution Bulletin, 72(2), 323–332. https://doi.org/10.1016/j.marpolbul.2012.09.006

Core Team, R. (2017). R: A Language and Environment for Statistical Computing. https://www.r-project.org/

Cox, D. R. (2007). Regression Models and Life-Tables. Journal of the Royal StatisticalSociety. Series B (Methodological), 34(2), 187–220.

D’Angelo, C., Hume, B. C. C., Burt, J., Smith, E. G., Achterberg, E. P., & Wiedenmann, J. (2015). Local adaptation constrains the distribution potential of heat-tolerant Symbiodinium from the Persian/Arabian Gulf. ISME Journal, 9(12), 2551–2560. https://doi.org/10.1038/ismej.2015.80

Davies, S. W., Treml, E. A., Kenkel, C. D., & Matz, M. v. (2015). Exploring the role of Micronesian islands in the maintenance of coral genetic diversity in the Pacific Ocean. Molecular Ecology, 24(1), 70–82. https://doi.org/10.1111/mec.13005

Day, T., Nagel, L., van Oppen, M. J. H., & Caley, M. J. (2008). Factors affecting the evolution of bleaching resistance in corals. American Naturalist, 171(2). https://doi.org/10.1086/524956

De’ath, G., Fabricius, K. E., Sweatman, H., & Puotinen, M. (2012). The 27 – year decline of coral cover on the Great Barrier Reef and its causes. PNAS, 109(44), 17995–17999. https://doi.org/10.1073/pnas.1208909109

Díaz, E., Schimmöller, F., & Pfeffer, S. R. (1997). A Novel Rab9 Effector Required for Endosome-to-TGN Transport. In The Journal of Cell Biology (Vol. 138, Issue 2).

Dixon, G., Davies, S., Aglyamova, G., Meyer, E., Bay, L. K., & Matz, M. v. (2015). Genomic Determinants of coral heat tolerance across latitudes. Science, 348(6242), 1460–1462.

Dohm, J. C., Lottaz, C., Borodina, T., & Himmelbauer, H. (2008). Substantial biases in ultra-short read data sets from high-throughput DNA sequencing. Nucleic Acids Research, 36(16). https://doi.org/10.1093/nar/gkn425

Downs, C. A., Kramarsky-Winter, E., Martinez, J., Kushmaro, A., Woodley, C. M., Loya, Y., & Ostrander, G. K. (2009). Symbiophagy as a cellular mechanism for coral bleaching. Autophagy, 5(2), 211–216. https://doi.org/10.4161/auto.5.2.7405

Drury, C., Bean, N., Harris, C., Hancock, J., Hucekba, J., Martin, C., Roach, T., Quinn, R., & Gates, R. D. (2021). Intrapopulation adaptive varience supports selective breeding in a reef building coral. BioArXIV, 1–19.

Dziedzic, K. E., Elder, H., & Meyer, E. (2019). Heritable variation in bleaching responses and its functional genomic basis in reef-building corals (Orbicella faveolata). Molecular Ecology, 1–16.

Falconer, D. S., & Mackay, T. F. C. (1996). Introduction to Quantitative Genetics.

Fuller, Z. L., Mocellin, V. J. L., Morris, L. A., Cantin, N., Shepherd, J., Sarre, L., Peng, J., Liao, Y., Pickrell, J., Andolfatto, P., Matz, M., Bay, L. K., & Przeworski, M. (2020). Population genetics of the coroal Acropora millepora: Toward genomic prediction of bleaching. Science, 369(6501). https://doi.org/10.1126/science.aba4674

Gienapp, P., Lof, M., Reed, T. E., McNamara, J., Verhulst, S., & Visser, M. E. (2013). Predicting demographically sustainable rates of adaptation: Can great tit breeding time keep pace with climate change? Philosophical Transactions of the Royal Society B: Biological Sciences, 368(1610). https://doi.org/10.1098/rstb.2012.0289

Glynn, P. W. (1993). Coral reef bleaching: ecological perspectives. Coral Reefs, 12, 1–17. https://doi.org/10.1007/BF00303779

Halestrap, A. P. (2012). The monocarboxylate transporter family-Structure and functional characterization. In IUBMB Life (Vol. 64, Issue 1, pp. 1–9). https://doi.org/10.1002/iub.573

Hoegh-Guldberg, O. (1999). Climate change, coral bleaching and the future of the world’s coral reefs. Marine and Freshwater Research, 50(8), 839–866. https://doi.org/10.1071/MF99078

Hoegh-Guldberg, O., Harvell, C. D., Sale, P. F., Edwards, A. J., Caldeira, K., Knowlton, N., Eakin, C. M., Muthiga, N., Bradbury, R. H., Dubi, A., Hatziolos, M. E., Mumby, P. J., Hooten, A., Steneck, R. S., Greenfield, P., Gomez, E., & Iglesias-Prieto, R. (2007). Coral Reefs Under Rapid Climate Change and Ocean Acidification. Science, 318(5857), 1737– 1742.

Howells, E. J., Abrego, D., Meyer, E., Kirk, N., & Burt, J. A. (2016). Host adaptation and unexpected symbiont partners enable reef-building corals to tolerate extreme temperatures. Global Change Biology, 1–13. https://doi.org/10.1111/gcb.13250

Hughes, T., Kerry, J., Baird, A., Connolly, S., Chase, T., Dietzel, A., Hill, T., Hoey, A., Hoogenboom, M., Jacobson, M., Kerswell, A., Madin, J., Mieog, A., Paley, A., Pratchett, M., Torda, G., & Woods, R. (2019). Global warming impairs stock -recruitment dynamics of corals. Nature, 568, 387–390. https://doi.org/10.1038/s41586-019-1081-y

Hughes, T. P., Kerry, J. T., Álvarez-noriega, M., Álvarez-romero, J. G., Anderson, K. D., Baird, A. H., Babcock, R. C., Beger, M., Bellwood, D. R., Berkelmans, R., Bridge, T. C., Butler, I. R., Byrne, M., Cantin, N. E., Comeau, S., Connolly, S. R., Cumming, G. S., Dalton, S. J., Diaz-Pulido, G., … Wilson, S. K. (2017). Global warming and recurrent mass bleaching of corals. Nature, 543, 373–377. https://doi.org/10.1038/nature21707

Hughes, T. P., Kerry, J. T., Baird, A. H., Connolly, S. R., Dietzel, A., Eakin, C. M., Heron, S. F., Hoey, A. S., Hoogenboom, M. O., Liu, G., Mcwilliam, M. J., Pears, R. J., Pratchett, M. S., Skirving, W. J., Stella, J. S., & Torda, G. (2018). Global warming transforms coral reef assemblages. Nature, 556, 492–496.

Hutchings, P., Kingsford, M., & Hoegh-Guldberg, O. (2019). The Great Barrier Reef: Biology, Environment, and Management. Csiro Publishing.

Ji, Y. K., Hong, Y., Ashida, H., Shishioh, N., Murakami, Y., Morita, Y. S., Maeda, Y., & Kinoshita, T. (2005). PIG-V involved in transferring the second mannose in glycosylphosphatidylinositol. Journal of Biological Chemistry, 280(10), 9489–9497. https://doi.org/10.1074/jbc.M413867200

Jin, Y. K., Lundgren, P., Lutz, A., Raina, J., Howells, E. J., Paley, A. S., Willis, B. L., & Oppen, M. J. H. van. (2016). Genetic markers for antioxidant capacity in a reef-building coral. May.

Juárez, O. E., Escobedo-Fregoso, C., Arredondo-Espinoza, R., & Ibarra, A. M. (2021). Development of SNP markers for identification of thermo-resistant families of the Pacific oyster Crassostrea gigas based on RNA-seq. Aquaculture, 539. https://doi.org/10.1016/j.aquaculture.2021.736618

Kaplan, E. L., & Meier, P. (2017). Nonparametric Estimation from Incomplete Observations. Journal of the American Statistical Association, 53(282), 457–481.

Kassambara, A. (2020). ggpubr R package: Create Publication Ready Plots. https://rpkgs.datanovia.com/ggpubr/

Kenkel, C. D., Matz, M. v, & Tx, A. (2016). Enhanced gene expression plasticity as a mechanism of adaptation to a variable environment in a reef-building coral. 3, 1–21.

Kenkel, C. D., Meyer, E., & Matz, M. v. (2013). Gene expression under chronic heat stress in populations of the mustard hill coral (Porites astreoides) from different thermal environments. Molecular Ecology, 22(16), 4322–4334. https://doi.org/10.1111/mec.12390

Kenkel, C. D., Setta, S. P., & Matz, M. v. (2015). Heritable differences in fitness-related traits among populations of the mustard hill coral, Porites astreoides. Heredity, 115(6), 509–516. https://doi.org/10.1038/hdy.2015.52

Kirk, N., Howells, E., Abrego, D., Burt, J., & Meyer, E. (2018). Genomic and transcriptomic signals of thermal tolerance in heat-tolerant corals (Platygyra daedalea) of the Arabian/Persian Gulf. Molecular Ecology, 27, 5180–5194.

Kofler, R., Orozco-terWengel, P., de Maio, N., Pandey, R. V., Nolte, V., Futschik, A., Kosiol, C., & Schlötterer, C. (2011). Popoolation: A toolbox for population genetic analysis of next generation sequencing data from pooled individuals. PLoS ONE, 6(1). https://doi.org/10.1371/journal.pone.0015925

Kofler, R., Pandey, R. V., Schlötterer, C., Populationsgenetik, I., Vienna, V., & Wien, A.-. (2017). PoPoolation2 : identifying differentiation between populations using sequencing of pooled DNA samples (Pool-Seq). Bioinformatics, 27(24), 3435–3436. https://doi.org/10.1093/bioinformatics/btr589

Kultz, D. (2005). Molecular And Evolutionary Basis of the Cellular Stress Response. Annual Review Physiology, 67, 225–257. https://doi.org/10.1146/annurev.physiol.67.040403.103635

Lajeunesse, T. C., Parkinson, J. E., Gabrielson, P. W., Jeong, H. J., Reimer, J. D., Voolstra, C. R., Santos, S. R., Lajeunesse, T. C., Parkinson, J. E., Gabrielson, P. W., Jeong, H. J., & Reimer, J. D. (2018). Systematic Revision of Symbiodiniaceae Highlights the Antiquity and Diversity of Coral Endosymbionts Article Systematic Revision of Symbiodiniaceae Highlights the Antiquity and Diversity of Coral Endosymbionts. Current Biology, 28(16), 2570-2580.e6. https://doi.org/10.1016/j.cub.2018.07.008

Langmead, B., Trapnell, C., Pop, M., & Salzberg, S. L. (2009). Ultrafast and memory-efficient alignment of short DNA sequences to the human genome. Genome Biology, 10(R25).

Lee, S. T. M., Davy, S. K., Tang, S. L., & Kench, P. S. (2016). Mucus sugar content shapes the bacterial community structure in thermally stressed Acropora muricata. Frontiers in Microbiology, 7(MAR). https://doi.org/10.3389/fmicb.2016.00371

Lodish, H., Berk, A., Kaiser, C. A., Krieger, M., Bretscher, A., Ploegh, H., Martin, K. C., Yaffe, M. B., & Amon, A. (2020). Molecular Cell Biology (9th ed.). Freeman.

Long Term Reef Monitoring Program. (2019). Annual Report on Reef Condition: Mixed bill of health for the Great Barrier Reef. https://www.aims.gov.au/reef-monitoring/gbr-condition-summary-2018-2019

Lynch, M., & Walsh, B. (1998). Genetics and Analysis of Quantitative Traits.

Manzello, D. P., Jankulak, M., Matz, M. v, Enochs, I. C., Valentino, L., Carlton, R. D., Kolodziej, G., Serrano, X., & Towle, E. K. (2019). Role of host genetics and heat - tolerant algal symbionts in sustaining populations of the endangered coral Orbicella faveolata in the Florida Keys with ocean warming. Global Change Biology, 25, 1016–1031. https://doi.org/10.1111/gcb.14545

Matz, M. v., Treml, E. A., & Haller, B. C. (2020). Estimating the potential for coral adaptation to global warming across the Indo-West Pacific. Global Change Biology, 26(6), 3473–3481. https://doi.org/10.1111/gcb.15060

Meyer, E., Davies, S., Wang, S., Willis, B., Abrego, D., Juenger, T., & Matz, M. (2009). Genetic variation in responses to a settlement cue and elevated temperature in the reef-building coral Acropora millepora. Marine Ecology Progress Series, 392, 81–92. https://doi.org/10.3354/meps08208

Meyer, K. (2007). WOMBAT — A tool for mixed model analyses in quantitative genetics by restricted maximum likelihood (REML). Journal of Zhejiang University Science B, 8(11), 815–821. https://doi.org/10.1631/jzus.2007.B0815

Oliver, T. A., & Palumbi, S. R. (2011). Many corals host thermally resistant symbionts in high-temperature habitat. Coral Reefs, 30, 241–250. https://doi.org/10.1007/s00338-010-0696-0

Pathmanathan, J. S., Williams, A., Stephens, T. G., Su, X., Eric, N., Conetta, D., Putnam, H. M., & Bhattacharya, D. (2021). Multi-omic characterization of the thermal stress phenome in the stony coral Montipora capitata. BioArXIV, 1–25.

Polato, N. R., Voolstra, C. R., Schnetzer, J., DeSalvo, M. K., Randall, C. J., Szmant, A. M., Medina, M., & Baums, I. B. (2010). Location-specific responses to thermal stress in larvae of the reef-building coral Montastraea faveolata. PloS One, 5(6), e11221. https://doi.org/10.1371/journal.pone.0011221

Prieto, D., Markaide, P., Urrutxurtu, I., Navarro, E., Artigaud, S., Fleury, E., Ibarrola, I., & Urrutia, M. B. (2019). Gill transcriptomic analysis in fast- and slow-growing individuals of Mytilus galloprovincialis. Aquaculture, 511. https://doi.org/10.1016/j.aquaculture.2019.734242

Purkis, S. J., & Riegl, B. M. (2012). Geomorphology and Reef Building in the SE Gulf. In Coral Reefs of the Gulf: Adaptation to Climatic Extremes in the World’s Hottest Sea (pp. 33–50). https://doi.org/10.1007/978-94-007-3008-3

Quigley, K. M., Bay, L. K., & van Oppen, M. J. H. (2020). Genome-wide SNP analysis reveals an increase in adaptive genetic variation through selective breeding of coral. Molecular Ecology, May. https://doi.org/10.1111/mec.15482

Riegl, B., Purkis, S., Al-cibahy, A., Abdel-moati, M., & Hoegh-Guldberg, O. (2011). Present Limits to Heat-Adaptability in Corals and Population-Level Responses to Climate Extremes. PloS One, 6(9). https://doi.org/10.1371/journal.pone.0024802

Ruiz-jones, L. J., & Palumbi, S. R. (2017). Tidal heat pulses on a reef trigger a fine-tuned transcriptional response in corals to maintain homeostasis. Science Advances, 3(e1601298), 1–11.

Rumble, S. M., Lacroute, P., Dalca, A. v., Fiume, M., Sidow, A., & Brudno, M. (2009). SHRiMP: Accurate mapping of short color-space reads. PLoS Computational Biology, 5(5), 1–11. https://doi.org/10.1371/journal.pcbi.1000386

Savolainen, O., Lascoux, M., & Merilä, J. (2013). Ecological genomics of local adaptation. Nature Reviews Genetics, 14(November), 807–820. https://doi.org/10.1038/nrg3522

Sella, G., & Barton, N. H. (2019). Thinking about the Evolution of Complex Traits in the Era of Genome-Wide Association Studies. Annual Review of Genomics and Human Genetics, 20, 461–493. https://doi.org/10.1146/annurev-genom-083115-022316

Silverstein, R. N., Cunning, R., & Baker, A. C. (2014). Change in algal symbiont communities after bleaching, not prior heat exposure, increases heat tolerance of reef corals. Global Change Biology, 236–249. https://doi.org/10.1111/gcb.12706

Singh, D., Singh, C. K., Sewak, R., Tomar, S., & Pal, M. (2017). Genetics and Molecular Mapping of Heat Tolerance for Seedling Survival and Pod Set in Lentil. Crop Science, 57, 3059–3067. https://doi.org/10.2135/cropsci2017.05.0284

Souter, D., Planes, S., Wicquart, J., Logan, M., Obura, D., & Staub, F. (2020). Status of Coral Reefs of the World: 2020 Chapter 2. Status of Coral Reefs of the World.

Soyk, S., Benoit, M., & Lippman, Z. B. (2020). New Horizons for Dissecting Epistasis in Crop Quantitative Trait Variation. https://doi.org/10.1146/annurev-genet-050720

Therneau, T. (2015). A Package for Survival Analysis in S. verson 2.38 (p. https://CRAN.R-project.org/package=survival).

Thomas, L., & Palumbi, S. R. (2017). The genomics of recovery from coral bleaching. Proceedings of the Royal Society B: Biological Sciences, 284(1865). https://doi.org/10.1098/rspb.2017.1790

Traylor-knowles, N., Rose, N. H., Sheets, E. A., & Palumbi, S. R. (2017). Early Transcriptional Responses during Heat Stress in the Coral Acropora hyacinthus. Biological Bulletin, 232, 91–100.

Treangen, T. J., & Salzberg, S. L. (2012). Repetitive DNA and next-generation sequencing: Computational challenges and solutions. Nature Reviews Genetics, 13(1), 36–46. https://doi.org/10.1038/nrg3117

van Oppen, M. J. H., Oliver, J. K., Putnam, H. M., & Gates, R. D. (2015). Building coral reef resilience through assisted evolution. Proceedings of the National Academy of Sciences, 112(8), 2307–2313. https://doi.org/10.1073/pnas.1422301112

Venuprasad, R., Dalid, C. O., del Valle, M., Zhoa, D., Espiritu, M., Sta Cruz, M. T., Amante, M., Kumar, A., & Atlin, G. N. (2009). Identification and characterization of large-effect quantitative trait loci for grain yield under lowland drought stress in rice using bulk-segregant analysis. Theoretical Applied Genetics, 120, 177–190. https://doi.org/10.1007/s00122-009-1168-1

Vikram, P., Swamy, B. P. M., Dixit, S., Ahmed, H., Cruz, M. T. S., Singh, A. K., Ye, G., & Kumar, A. (2012). Field Crops Research Bulk segregant analysis : “An effective approach for mapping consistent-effect drought grain yield QTLs in rice.” Field Crops Research, 134, 185–192. https://doi.org/10.1016/j.fcr.2012.05.012

Waite, T. A., & Campbell, L. G. (2006). Controlling the false discovery rate and increasing statistical power in ecological studies. Ecoscience, 13(4), 439–442. https://doi.org/10.2980/1195-6860(2006)13[439:CTFDRA]2.0.CO;2

Wang, M., Liu, W., Jiang, B., & Peng, Q. (2019). Genetic Analysis and Related Gene Primary Mapping of Heat Stress Tolerance in Cucumber Using Bulked Segregant Analysis. Horticulture Science, 54(3), 423–428. https://doi.org/10.21273/HORTSCI13734-18

Wang, S., Meyer, E., McKay, J., & Matz, M. (2012). 2b-RAD: a simple and flexible method for genome-wide genotyping. Nature Methods, 9(August), 808–810. http://www.nature.com/nmeth/journal/v9/n8/abs/nmeth.2023.html

Weeriyanun, P., Collins, R. B., Macadam, A., Kiff, H., Randle, J. L., & Quigley, K. M. (2021). Predicting selection-response gradients of heat tolerance in a wide-ranging reef-building coral. BioRxiv. https://doi.org/10.1101/2021.10.06.463349

Wellenreuther, M., & Hansson, B. (2016). Detecting Polygenic Evolution: Problems, Pitfalls, and Promises. In Trends in Genetics (Vol. 32, Issue 3, pp. 155–164). Elsevier Ltd. https://doi.org/10.1016/j.tig.2015.12.004

Wickham, H., Averick, M., Byan, J., Chang, W., McGowan, L. D., Francois, R., Grolemund, G., Hayes, A., Henry, L., Hester, J., Kuhn, M., Pedersen, T. L., Miller, E., Bache, S. M., Muller, K., Ooms, J., Robinson, D., Seidel, D. P., Spinu, V., … Yutani, H. (2019). Welcome to the Tidyverse. The Journal of Open Source Software, 4(43), 1–6.

Wickham, H., Francois, R., Henry, L., & Muller, K. (2020). dplyr: A Grammar of Data Manipulation (R package version 0.8.4). https://cran.r-project.org/package=dplyr

Wilson, A. J., Réale, D., Clements, M. N., Morrissey, M. M., Postma, E., Walling, C. A., Kruuk, L. E. B., & Nussey, D. H. (2010). An ecologist’s guide to the animal model. Journal of Animal Ecology, 79(1), 13–26. https://doi.org/10.1111/j.1365-2656.2009.01639.x

Woolsey, E. S., Keith, S. a., Byrne, M., Schmidt-Roach, S., & Baird, a. H. (2014). Latitudinal variation in thermal tolerance thresholds of early life stages of corals. Coral Reefs, 34, 471– 478. https://doi.org/10.1007/s00338-014-1253-z

Wright, R. M., Aglyamova, G. v., Meyer, E., & Matz, M. v. (2015). Gene expression associated with white syndromes in a reef building coral, Acropora hyacinthus. BMC Genomics, 16(1), 1–12. https://doi.org/10.1186/s12864-015-1540-2

Zhang, Y., Sun, J., Mu, H., Li, J., Zhang, Y., Xu, F., Xiang, Z., Qian, P. Y., Qiu, J. W., & Yu, Z. (2015). Proteomic basis of stress responses in the gills of the pacific oyster Crassostrea gigas. Journal of Proteome Research, 14(1), 304–317. https://doi.org/10.1021/pr500940s

Zhu, Y., Bergland, A. O., González, J., & Petrov, D. A. (2012). Empirical validation of pooled whole genome population re-sequencing in Drosophila melanogaster. PLoS ONE, 7(7), 1–7. https://doi.org/10.1371/journal.pone.0041901

