## SupplementalFigure2 for "Genetic variation in heat tolerance of the coral *Platygyra daedalea* indicates potential for adaptation to ocean warming"

**Supplementary Material: Cox Proportional Hazards Model Two Way Interaction Treatment by Time**

There was also a significant effect heat (p-value < 0.0001). At 240 hours, the survival probability for an individual at 35.5˚C was 0.598 (± 0.014), but 0.982 (± 0.008) at 27˚C (Supp. Fig. 2). The average survival curves estimated from all crosses in each temperature treatment show a significant difference in survival probability between the 35.5˚C treatment and 27˚C control (Supp. Fig. 2). We observed little mortality in control conditions throughout the 10-day experiment, whereas the heat stress treatment resulted in substantial mortality (Supp. Fig. 2).

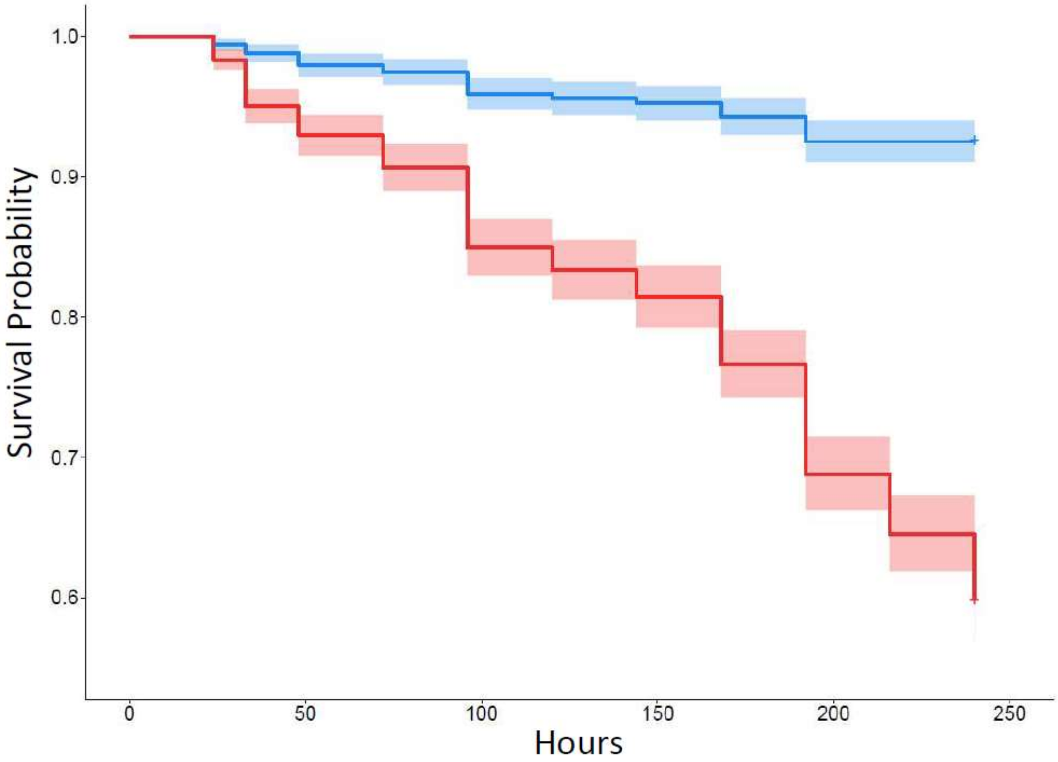


**Supplemental Fig. 2. Larval survival is reduced at elevated temperature.** Kaplan Meier plot shows cumulative probability of survival across all families in control 27˚C (blue) and elevated 35.5˚C (red) temperature. The shaded region represents 95% confidence intervals.
