## SupplementalGLMM for "Genetic variation in heat tolerance of the coral *Platygyra daedalea* indicates potential for adaptation to ocean warming"

**Supplemental Generalized Linear Mixed Model**

*Methods*

To evaluate the significance of potential variation introduced by position assignment, we fit a model to the percent larvae surviving using the nlme R package (Pinheiro et al. 2017). We modeled temperature, time, and cross as fixed effects and tank and plate positions as nested random effects. The fixed effect of time accounts for the repeated measures inherent to counting individual survival. Tank and plate assignments for each sample were fitted as nested random effects. We used penalized quasi-likelihood (PQL) estimation to account for the over dispersion of the survival data (Dean et al. 2004). Code available at <https://github.com/hollandelder/Platygyradaedalea_HeritabiltyandMarkersofThermalTolerance>.

*Results*

The full GLMM indicated that there was a significant interaction of cross x temperature x time (p-value < 0.0001). The random factors of position did not significantly contribute to variation in survival.
