## SupplementalTable1 for "Genetic variation in heat tolerance of the coral *Platygyra daedalea* indicates potential for adaptation to ocean warming"

**Supplemental Table 1**. The number and percent of reads after quality filtering steps and mapping to the de novo reference.

| Sample | Number Raw Reads | Number of Clean Reads | | Percent Reads Retained Q=30 | Reads Mapped One Or More Times | Percent Reads Mapped | Unique Mappings | Percent Unique Mappings |
| --- | --- | --- | --- | --- | --- | --- | --- | --- |
| AB2C_Pdae | 7962616 | 7580433 | 95.20028342 | | 6720183 | 88.65170367 | 4952737 | 65.33580602 |
| AB3C_Pdae | 4247054 | 4013046 | 94.49011009 | | 3612840 | 90.02737571 | 2348346 | 58.51779421 |
| AB4H_Pdae | 8576400 | 8181948 | 95.40072758 | | 7363968 | 90.00262529 | 5154778 | 63.00184259 |
| AC1C_Pdae | 6710966 | 6362051 | 94.80082301 | | 5780058 | 90.85211672 | 4375883 | 68.78101103 |
| AF1C_Pdae | 5972695 | 5712781 | 95.64829612 | | 5201386 | 91.04823028 | 3939184 | 68.95387728 |
| AC3C_Pdae | 11749736 | 11117592 | 94.61993018 | | 10091314 | 90.76888233 | 7558474 | 67.98661077 |
| AF3C_Pdae | 18276809 | 17303556 | 94.67492931 | | 15573100 | 89.99941977 | 10745439 | 62.09959964 |
| AC3H_Pdae | 10355095 | 9821248 | 94.84459582 | | 8970866 | 91.3414059 | 6447834 | 65.65188049 |
| AC4C_Pdae | 7007372 | 6589246 | 94.03305547 | | 5906425 | 89.63734242 | 4416176 | 67.02096112 |
| AF1H_Pdae | 6485287 | 6156738 | 94.93393276 | | 5594433 | 90.86683565 | 4228873 | 68.68690855 |
| AF2H_Pdae | 11392634 | 10789598 | 94.70679037 | | 9849844 | 91.29018523 | 7225102 | 66.96358845 |
| AF3H_Pdae | 7775745 | 7461527 | 95.95899814 | | 6850581 | 91.81205134 | 4603509 | 61.69660714 |
| BA1C_Pdae | 8511015 | 7962093 | 93.55045197 | | 7137078 | 89.63821447 | 5298402 | 66.54534178 |
| BA2C_Pdae | 10049968 | 9512117 | 94.64823172 | | 8635045 | 90.77942376 | 6410094 | 67.38872114 |
| BA2H_Pdae | 12871061 | 12195236 | 94.74926737 | | 11117743 | 91.16464003 | 8063741 | 66.12205783 |
| BA3C_Pdae | 13634624 | 12793118 | 93.82816864 | | 11336537 | 88.61433937 | 8689974 | 67.92694322 |
| BA3H_Pdae | 9098676 | 8567200 | 94.15875453 | | 7810547 | 91.16802456 | 5913766 | 69.02799048 |
| BA4C_Pdae | 1827765 | 1739958 | 95.19593602 | | 1608011 | 92.41665603 | 1092241 | 62.7739865 |
| BD1H_Pdae | 13107119 | 12317422 | 93.97505279 | | 11199956 | 90.9277607 | 8302413 | 67.40382038 |
| BD2C_Pdae | 10687185 | 10138688 | 94.86771306 | | 9267463 | 91.40692563 | 6920971 | 68.26298432 |
| BD4H_Pdae | 4270963 | 4065829 | 95.19700826 | | 3788154 | 93.17051947 | 2578460 | 63.41781713 |
| CB1H_Pdae | 7063261 | 6475916 | 91.68450663 | | 5789465 | 89.39993972 | 4365763 | 67.41537413 |
| CB2C_Pdae | 7473241 | 6903238 | 92.37274698 | | 6244011 | 90.45046687 | 4701957 | 68.1123409 |
| CB2H_Pdae | 11385563 | 10447162 | 91.75797455 | | 9431254 | 90.27575144 | 6818160 | 65.26327437 |
| CB3C_Pdae | 7406900 | 6963245 | 94.0102472 | | 6281953 | 90.21588354 | 4723857 | 67.83987925 |
| CB3H_Pdae | 3271715 | 3108744 | 95.01878984 | | 2860779 | 92.02362755 | 1949577 | 62.7126904 |
| CE1C_Pdae | 7062468 | 6671160 | 94.45933065 | | 6020310 | 90.24382566 | 4614903 | 69.17691976 |
| CE2C_Pdae | 10984549 | 10429576 | 94.94769426 | | 9653670 | 92.56052212 | 6647971 | 63.74152698 |
| CE3H_Pdae | 5113005 | 4833714 | 94.53763491 | | 4422646 | 91.49581461 | 3163964 | 65.4561689 |
| FA1C_Pdae | 10397740 | 9920958 | 95.41456124 | | 9123113 | 91.9579843 | 6912554 | 69.67627521 |
| FA1H_Pdae | 10015235 | 9491478 | 94.7703973 | | 8725463 | 91.9294445 | 6336518 | 66.76007678 |
| FA2C_Pdae | 10435895 | 9863922 | 94.51917636 | | 8891981 | 90.14650562 | 6686345 | 67.78586651 |
| FA4C_Pdae | 5184640 | 4894219 | 94.39843461 | | 4513437 | 92.21975968 | 3220163 | 65.7952372 |
| FE1C_Pdae | 12013236 | 11377599 | 94.70886113 | | 10139558 | 89.1186093 | 7681722 | 67.51619564 |
| FE1H_Pdae | 8299377 | 7785293 | 93.80575193 | | 7131035 | 91.59623151 | 5035758 | 64.68296055 |
| FE2C_Pdae | 7683276 | 7224953 | 94.03479714 | | 6587671 | 91.17943051 | 4909684 | 67.95454586 |
| FE2H_Pdae | 6219234 | 5898104 | 94.83650237 | | 5451828 | 92.43356848 | 3756115 | 63.68343115 |
| FE4H_Pdae | 8658862 | 8257634 | 95.3662733 | | 7703052 | 93.28400847 | 5145377 | 62.31054803 |
| AB1C_Pdae | 6249043 | 5686906 | 91.0044306 | | 5119516 | 90.02287008 | 3696489 | 65.00000176 |
| AB1H_Pdae | 9014751 | 8615047 | 95.56611159 | | 7842094 | 91.02787251 | 5773081 | 67.01160191 |
| AB2H_Pdae | 5150229 | 4907369 | 95.28448152 | | 4171364 | 85.00204488 | 3043668 | 62.02239938 |
| AB3H_Pdae | 5209236 | 5003730 | 96.05496852 | | 4756343 | 95.05594826 | 3256242 | 65.07629309 |
| AB4C_Pdae | 5188565 | 4958593 | 95.56771477 | | 4463533 | 90.01611949 | 3460109 | 69.78005656 |
| AC1H_Pdae | 10400198 | 9879516 | 94.99353762 | | 8948293 | 90.57420424 | 6364412 | 64.42028132 |
| AC2C_Pdae | 9108736 | 8642314 | 94.87939929 | | 7813897 | 90.41440753 | 5876815 | 68.00047996 |
| AF2C_Pdae | 11031208 | 10553843 | 95.67259542 | | 9471491 | 89.74447507 | 6697541 | 63.46068442 |
| AF4C_Pdae | 8351410 | 8056311 | 96.46647692 | | 7341353 | 91.12549156 | 5455724 | 67.71987824 |
| AC4H_Pdae | 6106115 | 5764439 | 94.40436349 | | 5269830 | 91.41965072 | 3585695 | 62.20371141 |
| AF4H_Pdae | 7926939 | 7609184 | 95.99145395 | | 7056740 | 92.73977341 | 4782404 | 62.85041865 |
| BA1H_Pdae | 7528955 | 7238785 | 96.14594588 | | 6723638 | 92.88351567 | 4576916 | 63.22768255 |
| BA4H_Pdae | 10068173 | 9649253 | 95.83916566 | | 8975951 | 93.02223706 | 6118550 | 63.4095717 |
| BD1C_Pdae | 13561965 | 12809712 | 94.4532153 | | 11424483 | 89.18610348 | 8597239 | 67.11500618 |
| BD2H_Pdae | 10290812 | 9799810 | 95.22873414 | | 8985258 | 91.68808375 | 6630785 | 67.66238325 |
| BD3H_Pdae | 9553327 | 9083525 | 95.08232054 | | 8265804 | 90.99775693 | 6183599 | 68.07488282 |
| BD4C_Pdae | 7854994 | 7507059 | 95.57052494 | | 6682582 | 89.01731024 | 5180870 | 69.01331134 |
| CB1C_Pdae | 11322811 | 10767050 | 95.09166937 | | 9851640 | 91.49804264 | 7320015 | 67.98533489 |
| CB4C_Pdae | 7779142 | 7302747 | 93.87599558 | | 6433019 | 88.09039941 | 4839656 | 66.2717194 |
| CB4H_Pdae | 8194257 | 7638048 | 93.21220948 | | 6849758 | 89.67943118 | 5123498 | 67.07863056 |
| CD1C_Pdae | 9298995 | 8613415 | 92.62737532 | | 7656079 | 88.88552334 | 5747416 | 66.72633328 |
| CD1H_Pdae | 12839583 | 12286396 | 95.69155011 | | 11080504 | 90.18514461 | 7784114 | 63.35555195 |
| CD2C_Pdae | 6491417 | 6005888 | 92.52044661 | | 5371532 | 89.43776507 | 4055259 | 67.52138901 |
| CD3C_Pdae | 10079153 | 9538011 | 94.63107664 | | 8600511 | 90.1709067 | 6423114 | 67.34227922 |
| CD3H_Pdae | 6601452 | 6280422 | 95.13697896 | | 5715771 | 91.00934619 | 4260586 | 67.8391675 |
| CD4C_Pdae | 9361859 | 8862510 | 94.66613415 | | 7980673 | 90.0498053 | 5982866 | 67.50757968 |
| CD4H_Pdae | 9907614 | 9517241 | 96.0598687 | | 8427618 | 88.55106222 | 5791946 | 60.85740605 |
| CE1H_Pdae | 10137793 | 9685186 | 95.5354484 | | 8992526 | 92.84825299 | 6274165 | 64.78104809 |
| CE2H_Pdae | 8854224 | 8361071 | 94.43030807 | | 7583115 | 90.69549822 | 5640451 | 67.46086715 |
| CE3C_Pdae | 6943560 | 6432239 | 92.63603973 | | 5723352 | 88.9791564 | 4297764 | 66.8159874 |
| CE4C_Pdae | 11814426 | 11191813 | 94.73006137 | | 10038238 | 89.69268875 | 7512182 | 67.12211864 |
| FA2H_Pdae | 11383324 | 10889559 | 95.66238297 | | 9906318 | 90.97079138 | 7344960 | 67.44956338 |
| FA3C_Pdae | 6513515 | 6239013 | 95.7856549 | | 5697396 | 91.31886726 | 4280201 | 68.60381602 |
| FA3H_Pdae | 8490101 | 8039965 | 94.69810783 | | 7358499 | 91.52401783 | 5381245 | 66.93119933 |
| FE3C_Pdae | 7633368 | 7246301 | 94.9292763 | | 6598722 | 91.0633163 | 4765460 | 65.76403602 |
| FE3H_Pdae | 3936600 | 3761057 | 95.54074582 | | 3504956 | 93.1907174 | 2385360 | 63.42259636 |
| FE4C_Pdae | 9529754 | 9106377 | 95.55731449 | | 7974124 | 87.56637244 | 5930189 | 65.1212771 |
| BD3C_Pdae | 9312969 | 8808966 | 94.5881598 | | 8014164 | 90.97735194 | 6032207 | 68.47803704 |
| CE4H_Pdae | 9553437 | 9128542 | 95.55243835 | | 8451834 | 92.58689942 | 5806765 | 63.61108926 |
| FA4H_Pdae | 5515231 | 5283326 | 95.79518972 | | 4923554 | 93.19042588 | 3249682 | 61.50826203 |
|  |  | Median | 95 | | Median | 91 | Median | 67 |
|  |  | Mean | 95 | | Mean | 91 | Mean | 66 |
