## SupplementalTable2 for "Genetic variation in heat tolerance of the coral *Platygyra daedalea* indicates potential for adaptation to ocean warming"

**Supplemental Table 2**. Markers identified in three or fewer crosses.

| Marker | SNP Location | Marker Found in Crosses | Number Crosses |
| --- | --- | --- | --- |
| denovoLocus104310 | 4 | BC, DC, EF | 3 |
| denovoLocus10613 | 11, 29 | CA, DC, EC | 3 |
| denovoLocus13278 | 1 | DC, EC, EF | 3 |
| denovoLocus13680 | 25, 34 | CA, DB, EF | 3 |
| denovoLocus14265 | 18 | AB, EC, EF | 3 |
| denovoLocus16627 | 3 | BC, DB, DC | 3 |
| denovoLocus1685 | 33 | BA, BC, CA | 3 |
| denovoLocus18867 | 6 | BC, DC, FA | 3 |
| denovoLocus22915 | 34 | AF, BA, CA | 3 |
| denovoLocus24499 | 21, 27 | AF, DC, EF | 3 |
| denovoLocus2572 | 34 | CA, DC, FA | 3 |
| denovoLocus30715 | 2, 29 | DC, EC, FA | 3 |
| denovoLocus32940 | 8 | AF, DC, EF | 3 |
| denovoLocus38500 | 8 | AB, BA, EF | 3 |
| denovoLocus49341 | 32 | AF, CA, FA | 3 |
| denovoLocus52938 | 27 | AF, EF, FA | 3 |
| denovoLocus58853 | 6, 30 | AF, BA, FA | 3 |
| denovoLocus6036 | 6 | AB, BA, BC | 3 |
| denovoLocus6154 | 7 | BA, DC, FA | 3 |
| denovoLocus6164 | 36 | AB, BA, CA | 3 |
| denovoLocus75321 | 19 | BC, CA, DC | 3 |
| denovoLocus85411 | 7 | AB, EC, FA | 3 |
| denovoLocus8753 | 32 | BC, CA, EC | 3 |
| denovoLocus10129 | 1, 34 | EC, EF | 2 |
| denovoLocus10133 | 33 | DC, EC | 2 |
| denovoLocus105160 | 32 | DC, FA | 2 |
| denovoLocus105818 | 36 | EF, FA | 2 |
| denovoLocus10860 | 6, 36 | DC, EF | 2 |
| denovoLocus1110 | 8 | DC, FA | 2 |
| denovoLocus111812 | 11 | DB, EF | 2 |
| denovoLocus111991 | 34 | DC, EF | 2 |
| denovoLocus11443 | 32 | AB, BA | 2 |
| denovoLocus11595 | 25 | DC, FA | 2 |
| denovoLocus11764 | 19 | BA, DC | 2 |
| denovoLocus12018 | 36 | AB, DC | 2 |
| denovoLocus12258 | 27 | AB, CA | 2 |
| denovoLocus13663 | 30 | DC, EF | 2 |
| denovoLocus13732 | 3 | DC, EC | 2 |
| denovoLocus13835 | 5 | EC, EF | 2 |
| denovoLocus1416 | 16, 36 | AF, BA | 2 |
| denovoLocus14360 | 30 | EC, FA | 2 |
| Marker | SNP Location | Marker Found in Crosses | Number Crosses |
| denovoLocus14433 | 20, 34 | BA, DC | 2 |
| denovoLocus1480 | 5 | EF, FA | 2 |
| denovoLocus14833 | 33 | AB, BA | 2 |
| denovoLocus15476 | 27 | BC, DC | 2 |
| denovoLocus15899 | 34 | BA, CA | 2 |
| denovoLocus16027 | 34 | DB, DC | 2 |
| denovoLocus16414 | 6 | DC, EC | 2 |
| denovoLocus1668 | 26 | AB, CA | 2 |
| denovoLocus17167 | 1 | BA, CA | 2 |
| denovoLocus17994 | 31 | AB, DC | 2 |
| denovoLocus19024 | 3 | EF, FA | 2 |
| denovoLocus19325 | 32 | AB, BA | 2 |
| denovoLocus19951 | 32 | CA, DC | 2 |
| denovoLocus20019 | 34 | DC, EF | 2 |
| denovoLocus20580 | 36 | BA, BC | 2 |
| denovoLocus21802 | 11, 32 | AF, DC | 2 |
| denovoLocus22104 | 17 | BC, FA | 2 |
| denovoLocus22267 | 4 | AB, BC | 2 |
| denovoLocus23162 | 32, 36 | DC, EF | 2 |
| denovoLocus23425 | 4 | BA, CA | 2 |
| denovoLocus23712 | 10 | DC, FA | 2 |
| denovoLocus24254 | 21 | BA, CA | 2 |
| denovoLocus24558 | 3 | CA, DC | 2 |
| denovoLocus24976 | 19, 28 | DC, EF | 2 |
| denovoLocus25267 | 26, 29 | AF, BC | 2 |
| denovoLocus25366 | 34 | DC, FA | 2 |
| denovoLocus25423 | 33 | DC, FA | 2 |
| denovoLocus26881 | 36 | AB, FA | 2 |
| denovoLocus27225 | 11, 25 | BA, EF | 2 |
| denovoLocus27975 | 25 | BA, FA | 2 |
| denovoLocus2899 | 36 | BC, DC | 2 |
| denovoLocus29455 | 18 | CA, FA | 2 |
| denovoLocus3033 | 25 | BC, DC | 2 |
| denovoLocus30716 | 1, 36 | CA, EF | 2 |
| denovoLocus31095 | 34 | EC, EF | 2 |
| denovoLocus311 | 28 | EC, EF | 2 |
| denovoLocus31194 | 28 | CA, EC | 2 |
| denovoLocus31423 | 7 | DC, FA | 2 |
| denovoLocus31648 | 32 | AB, BA | 2 |
| denovoLocus32401 | 21, 33 | BC, EC | 2 |
| denovoLocus32448 | 33 | BA, EF | 2 |
| denovoLocus32998 | 3 | DB, FA | 2 |
| Marker | SNP Location | Marker Found in Crosses | Number Crosses |
| denovoLocus33479 | 34 | AF, EF | 2 |
| denovoLocus36935 | 1 | DC, FA | 2 |
| denovoLocus3805 | 3 | EC, EF | 2 |
| denovoLocus38089 | 31 | BC, EC | 2 |
| denovoLocus38700 | 18 | AB, CA | 2 |
| denovoLocus39998 | 34 | DC, EC | 2 |
| denovoLocus41168 | 20 | EC, EF | 2 |
| denovoLocus41205 | 28 | DC, FA | 2 |
| denovoLocus44024 | 16 | AF, FA | 2 |
| denovoLocus44176 | 3 | DB, DC | 2 |
| denovoLocus44384 | 1 | EC, EF | 2 |
| denovoLocus44692 | 6 | EC, EF | 2 |
| denovoLocus45468 | 31 | DC, EC | 2 |
| denovoLocus46004 | 32 | CA, DC | 2 |
| denovoLocus47367 | 11 | AF, EF | 2 |
| denovoLocus48424 | 17, 35 | AB, FA | 2 |
| denovoLocus48746 | 9 | BA, DC | 2 |
| denovoLocus49247 | 25 | DC, FA | 2 |
| denovoLocus49498 | 21 | BC, EC | 2 |
| denovoLocus49606 | 33 | DC, EF | 2 |
| denovoLocus49865 | 32 | BA, DC | 2 |
| denovoLocus52651 | 9, 17 | DC, EF | 2 |
| denovoLocus53380 | 27 | AB, AF | 2 |
| denovoLocus55443 | 21 | DB, DC | 2 |
| denovoLocus57700 | 6, 28 | BA, DC | 2 |
| denovoLocus5881 | 4 | BC, EC | 2 |
| denovoLocus6000 | 27 | AF, BC | 2 |
| denovoLocus60119 | 3 | BC, CA | 2 |
| denovoLocus60238 | 27 | AF, FA | 2 |
| denovoLocus6041 | 1, 18 | BA, EC | 2 |
| denovoLocus62710 | 17 | CA, DC | 2 |
| denovoLocus64963 | 27 | DC, FA | 2 |
| denovoLocus6620 | 36 | DC, EF | 2 |
| denovoLocus67017 | 16 | EF, FA | 2 |
| denovoLocus68 | 32 | DC, EC | 2 |
| denovoLocus6969 | 1 | BA, DC | 2 |
| denovoLocus71438 | 6 | AB, DC | 2 |
| denovoLocus73508 | 36 | AB, DC | 2 |
| denovoLocus7920 | 11, 25 | BC, FA | 2 |
| denovoLocus79497 | 1 | AF, EC | 2 |
| denovoLocus80641 | 3 | CA, DC | 2 |
| denovoLocus81929 | 31 | BC, CA | 2 |
| Marker | SNP Location | Marker Found in Crosses | Number Crosses |
| denovoLocus83186 | 1 | AB, AF | 2 |
| denovoLocus84456 | 33 | DC, FA | 2 |
| denovoLocus8450 | 28 | DC, EF | 2 |
| denovoLocus85726 | 19 | DB, DC | 2 |
| denovoLocus8626 | 27 | AB, CA | 2 |
| denovoLocus8751 | 3 | CA, EF | 2 |
| denovoLocus89683 | 16 | BA, CA | 2 |
| denovoLocus9237 | 18 | AF, FA | 2 |
| denovoLocus94104 | 34 | BA, EF | 2 |
| denovoLocus960 | 4 | AF, EC | 2 |
| denovoLocus9987 | 29 | CA, DC | 2 |
