## SupplementalTable3 for "Genetic variation in heat tolerance of the coral *Platygyra daedalea* indicates potential for adaptation to ocean warming"

**Supplemental Table 3**. A comparison of the number and percent of tags genotyped in the first cross (C1) versus the set tags in the second cross (C2) post quality filtering.

| C1toC2 | Cross1 | Cross2 | NumTagsInBoth | TotalTagsC2 | NumReadsC1NotGeno | PercentGenotyped | PercentNotGenotyped |
| --- | --- | --- | --- | --- | --- | --- | --- |
| DBtoBA | DB | BA | 7254 | 23999 | 16745 | 30.22625943 | 69.77374057 |
| BCtoBA | BC | BA | 7109 | 22820 | 15711 | 31.15249781 | 68.84750219 |
| CAtoBA | CA | BA | 7301 | 23173 | 15872 | 31.50649463 | 68.49350537 |
| DBtoEF | DB | EF | 7920 | 23999 | 16079 | 33.00137506 | 66.99862494 |
| CAtoEF | CA | EF | 7845 | 23173 | 15328 | 33.85405429 | 66.14594571 |
| BCtoEF | BC | EF | 7766 | 22820 | 15054 | 34.03155127 | 65.96844873 |
| AFtoBA | AF | BA | 7194 | 19399 | 12205 | 37.08438579 | 62.91561421 |
| DCtoBA | DC | BA | 6582 | 17548 | 10966 | 37.50854798 | 62.49145202 |
| DBtoAB | DB | AB | 9085 | 23999 | 14914 | 37.85574399 | 62.14425601 |
| DBtoFA | DB | FA | 9200 | 23999 | 14799 | 38.33493062 | 61.66506938 |
| DBtoEC | DB | EC | 9288 | 23999 | 14711 | 38.70161257 | 61.29838743 |
| BCtoAB | BC | AB | 9011 | 22820 | 13809 | 39.48729185 | 60.51270815 |
| ECtoBA | EC | BA | 6615 | 16734 | 10119 | 39.5302976 | 60.4697024 |
| CAtoAB | CA | AB | 9193 | 23173 | 13980 | 39.67116903 | 60.32883097 |
| BCtoFA | BC | FA | 9141 | 22820 | 13679 | 40.05696757 | 59.94303243 |
| AFtoEF | AF | EF | 8159 | 19399 | 11240 | 42.05886901 | 57.94113099 |
| AFtoDC | AF | DC | 8196 | 19399 | 11203 | 42.24960049 | 57.75039951 |
| DBtoDC | DB | DC | 10181 | 23999 | 13818 | 42.42260094 | 57.57739906 |
| AFtoEC | AF | EC | 8232 | 19399 | 11167 | 42.43517707 | 57.56482293 |
| AFtoAB | AF | AB | 8240 | 19399 | 11159 | 42.47641631 | 57.52358369 |
| CAtoEC | CA | EC | 10052 | 23173 | 13121 | 43.3780693 | 56.6219307 |
| CAtoFA | CA | FA | 10089 | 23173 | 13084 | 43.53773788 | 56.46226212 |
| BCtoEC | BC | EC | 9950 | 22820 | 12870 | 43.60210342 | 56.39789658 |
| DCtoEF | DC | EF | 7699 | 17548 | 9849 | 43.87394575 | 56.12605425 |
| BCtoAF | BC | AF | 10124 | 22820 | 12696 | 44.36459246 | 55.63540754 |
| CAtoDC | CA | DC | 10283 | 23173 | 12890 | 44.37491909 | 55.62508091 |
| BCtoDC | BC | DC | 10216 | 22820 | 12604 | 44.76774759 | 55.23225241 |
| EFtoBA | EF | BA | 6278 | 13904 | 7626 | 45.15247411 | 54.84752589 |
| DBtoAF | DB | AF | 10882 | 23999 | 13117 | 45.34355598 | 54.65644402 |
| DCtoAF | DC | AF | 8139 | 17548 | 9409 | 46.381354 | 53.618646 |
| DCtoAB | DC | AB | 8337 | 17548 | 9211 | 47.50968771 | 52.49031229 |
| CAtoAF | CA | AF | 11100 | 23173 | 12073 | 47.90057394 | 52.09942606 |
| ECtoAB | EC | AB | 8106 | 16734 | 8628 | 48.44030118 | 51.55969882 |
| FAtoEF | FA | EF | 8089 | 16691 | 8602 | 48.46324366 | 51.53675634 |
| ECtoEF | EC | EF | 8124 | 16734 | 8610 | 48.54786662 | 51.45213338 |
| ABtoBA | AB | BA | 7419 | 15217 | 7798 | 48.75468226 | 51.24531774 |
| ABtoEF | AB | EF | 7445 | 15217 | 7772 | 48.9255438 | 51.0744562 |
| ECtoAF | EC | AF | 8232 | 16734 | 8502 | 49.19325923 | 50.80674077 |
| AFtoFA | AF | FA | 9590 | 19399 | 9809 | 49.43553791 | 50.56446209 |
| DCtoFA | DC | FA | 8846 | 17548 | 8702 | 50.41030317 | 49.58969683 |
| DBtoCA | DB | CA | 12345 | 23999 | 11654 | 51.43964332 | 48.56035668 |
| ECtoFA | EC | FA | 8632 | 16734 | 8102 | 51.58360225 | 48.41639775 |
| FAtoEC | FA | EC | 8637 | 16691 | 8054 | 51.74645018 | 48.25354982 |
| AFtoBC | AF | BC | 10124 | 19399 | 9275 | 52.18825713 | 47.81174287 |
| FAtoAB | FA | AB | 8814 | 16691 | 7877 | 52.80690192 | 47.19309808 |
| FAtoBA | FA | BA | 8814 | 16691 | 7877 | 52.80690192 | 47.19309808 |
| FAtoDC | FA | DC | 8846 | 16691 | 7845 | 52.99862201 | 47.00137799 |
| ABtoEC | AB | EC | 8106 | 15217 | 7111 | 53.26936978 | 46.73063022 |
| CAtoDB | CA | DB | 12345 | 23173 | 10828 | 53.27320589 | 46.72679411 |
| EFtoAB | EF | AB | 7445 | 13904 | 6459 | 53.54574223 | 46.45425777 |
| BAtoEF | BA | EF | 6278 | 11616 | 5338 | 54.04614325 | 45.95385675 |
| ABtoAF | AB | AF | 8240 | 15217 | 6977 | 54.14996386 | 45.85003614 |
| DBtoBC | DB | BC | 13098 | 23999 | 10901 | 54.57727405 | 45.42272595 |
| FAtoBC | FA | BC | 9141 | 16691 | 7550 | 54.76604158 | 45.23395842 |
| DCtoEC | DC | EC | 9613 | 17548 | 7935 | 54.78117164 | 45.21882836 |
| ABtoDC | AB | DC | 8337 | 15217 | 6880 | 54.78740882 | 45.21259118 |
| FAtoDB | FA | DB | 9200 | 16691 | 7491 | 55.11952549 | 44.88047451 |
| EFtoDC | EF | DC | 7699 | 13904 | 6205 | 55.37255466 | 44.62744534 |
| ECtoDB | EC | DB | 9288 | 16734 | 7446 | 55.50376479 | 44.49623521 |
| CAtoBC | CA | BC | 12939 | 23173 | 10234 | 55.8365339 | 44.1634661 |
| EFtoBC | EF | BC | 7766 | 13904 | 6138 | 55.85443038 | 44.14556962 |
| AFtoDB | AF | DB | 10882 | 19399 | 8517 | 56.09567503 | 43.90432497 |
| EFtoCA | EF | CA | 7845 | 13904 | 6059 | 56.4226122 | 43.5773878 |
| BAtoDC | BA | DC | 6582 | 11616 | 5034 | 56.66322314 | 43.33677686 |
| BCtoCA | BC | CA | 12939 | 22820 | 9881 | 56.70026293 | 43.29973707 |
| BAtoEC | BA | EC | 6615 | 11616 | 5001 | 56.94731405 | 43.05268595 |
| EFtoDB | EF | DB | 7920 | 13904 | 5984 | 56.96202532 | 43.03797468 |
| AFtoCA | AF | CA | 11100 | 19399 | 8299 | 57.2194443 | 42.7805557 |
| BCtoDB | BC | DB | 13098 | 22820 | 9722 | 57.39702016 | 42.60297984 |
| ECtoDC | EC | DC | 9613 | 16734 | 7121 | 57.44591849 | 42.55408151 |
| FAtoAF | FA | AF | 9590 | 16691 | 7101 | 57.45611407 | 42.54388593 |
| ABtoFA | AB | FA | 8814 | 15217 | 6403 | 57.92206085 | 42.07793915 |
| DCtoDB | DC | DB | 10181 | 17548 | 7367 | 58.01800775 | 41.98199225 |
| EFtoFA | EF | FA | 8089 | 13904 | 5815 | 58.17750288 | 41.82249712 |
| DCtoBC | DC | BC | 10216 | 17548 | 7332 | 58.21746068 | 41.78253932 |
| EFtoEC | EF | EC | 8124 | 13904 | 5780 | 58.429229 | 41.570771 |
| DCtoCA | DC | CA | 10283 | 17548 | 7265 | 58.59927057 | 41.40072943 |
| EFtoAF | EF | AF | 8159 | 13904 | 5745 | 58.68095512 | 41.31904488 |
| ABtoBC | AB | BC | 9011 | 15217 | 6206 | 59.21666557 | 40.78333443 |
| ECtoBC | EC | BC | 9950 | 16734 | 6784 | 59.45978248 | 40.54021752 |
| ABtoDB | AB | DB | 9085 | 15217 | 6132 | 59.70296379 | 40.29703621 |
| ECtoCA | EC | CA | 10052 | 16734 | 6682 | 60.06931995 | 39.93068005 |
| BAtoFA | BA | FA | 7003 | 11616 | 4613 | 60.28753444 | 39.71246556 |
| ABtoCA | AB | CA | 9193 | 15217 | 6024 | 60.41269633 | 39.58730367 |
| FAtoCA | FA | CA | 10089 | 16691 | 6602 | 60.44574921 | 39.55425079 |
| BAtoBC | BA | BC | 7109 | 11616 | 4507 | 61.20006887 | 38.79993113 |
| BAtoAF | BA | AF | 7194 | 11616 | 4422 | 61.93181818 | 38.06818182 |
| BAtoDB | BA | DB | 7254 | 11616 | 4362 | 62.44834711 | 37.55165289 |
| BAtoCA | BA | CA | 7301 | 11616 | 4315 | 62.85296143 | 37.14703857 |
| BAtoAB | BA | AB | 7419 | 11616 | 4197 | 63.86880165 | 36.13119835 |
|  |  |  |  |  | Median | 52.80690192 | 47.19309808 |
|  |  |  |  |  | Mean | 50.28564063 | 49.71435937 |
