## SupplementalTable4 for "Genetic variation in heat tolerance of the coral *Platygyra daedalea* indicates potential for adaptation to ocean warming"

**Supplemental Table 4**. A comparison of the number and percent of tags genotyped in the first cross (80.1) versus the set tags in the second cross (80.2) post the 80X filter.

| 80to80 | 80.1 | 80.2 | Num80.1in80.2 | Total80.1 | NumNotGenoIn80.2 | PercentGenotyped | PercentNotGenotyped |
| --- | --- | --- | --- | --- | --- | --- | --- |
| DBtoBA | DB | BA | 2250 | 10699 | 8449 | 21.0300028 | 78.9699972 |
| AFtoBA | AF | BA | 2268 | 8375 | 6107 | 27.08059701 | 72.91940299 |
| DBtoEF | DB | EF | 3060 | 10699 | 7639 | 28.60080381 | 71.39919619 |
| FAtoBA | FA | BA | 2277 | 7828 | 5551 | 29.08788963 | 70.91211037 |
| DCtoBA | DC | BA | 2149 | 7305 | 5156 | 29.41820671 | 70.58179329 |
| DBtoEC | DB | EC | 3211 | 10699 | 7488 | 30.01215067 | 69.98784933 |
| BCtoBA | BC | BA | 2077 | 6911 | 4834 | 30.05353784 | 69.94646216 |
| CAtoBA | CA | BA | 2290 | 7076 | 4786 | 32.3629169 | 67.6370831 |
| ECtoBA | EC | BA | 2084 | 6371 | 4287 | 32.71072045 | 67.28927955 |
| ABtoBA | AB | BA | 2515 | 7497 | 4982 | 33.54675203 | 66.45324797 |
| EFtoBA | EF | BA | 1997 | 5668 | 3671 | 35.23288638 | 64.76711362 |
| DBtoCA | DB | CA | 3826 | 10699 | 6873 | 35.76035143 | 64.23964857 |
| AFtoEC | AF | EC | 3046 | 8375 | 5329 | 36.37014925 | 63.62985075 |
| DBtoDC | DB | DC | 3985 | 10699 | 6714 | 37.24647163 | 62.75352837 |
| DBtoFA | DB | FA | 3995 | 10699 | 6704 | 37.33993831 | 62.66006169 |
| BCtoEF | BC | EF | 2612 | 6911 | 4299 | 37.79481985 | 62.20518015 |
| DBtoBC | DB | BC | 4071 | 10699 | 6628 | 38.05028507 | 61.94971493 |
| ABtoEF | AB | EF | 2954 | 7497 | 4543 | 39.40242764 | 60.59757236 |
| DBtoAB | DB | AB | 4222 | 10699 | 6477 | 39.46163193 | 60.53836807 |
| DCtoEF | DC | EF | 2891 | 7305 | 4414 | 39.57563313 | 60.42436687 |
| AFtoEF | AF | EF | 3318 | 8375 | 5057 | 39.61791045 | 60.38208955 |
| AFtoBC | AF | BC | 3367 | 8375 | 5008 | 40.20298507 | 59.79701493 |
| AFtoDC | AF | DC | 3421 | 8375 | 4954 | 40.84776119 | 59.15223881 |
| FAtoEF | FA | EF | 3203 | 7828 | 4625 | 40.91722024 | 59.08277976 |
| CAtoEF | CA | EF | 2898 | 7076 | 4178 | 40.955342 | 59.044658 |
| FAtoEC | FA | EC | 3232 | 7828 | 4596 | 41.28768523 | 58.71231477 |
| FAtoBC | FA | BC | 3242 | 7828 | 4586 | 41.41543178 | 58.58456822 |
| DBtoAF | DB | AF | 4532 | 10699 | 6167 | 42.35909898 | 57.64090102 |
| ABtoEC | AB | EC | 3206 | 7497 | 4291 | 42.76377218 | 57.23622782 |
| FAtoDC | FA | DC | 3580 | 7828 | 4248 | 45.7332652 | 54.2667348 |
| EFtoBC | EF | BC | 2612 | 5668 | 3056 | 46.08327452 | 53.91672548 |
| ECtoEF | EC | EF | 2941 | 6371 | 3430 | 46.16229791 | 53.83770209 |
| AFtoCA | AF | CA | 3872 | 8375 | 4503 | 46.23283582 | 53.76716418 |
| BCtoEC | BC | EC | 3215 | 6911 | 3696 | 46.52004052 | 53.47995948 |
| AFtoAB | AF | AB | 3905 | 8375 | 4470 | 46.62686567 | 53.37313433 |
| DCtoAF | DC | AF | 3421 | 7305 | 3884 | 46.83093771 | 53.16906229 |
| BCtoFA | BC | FA | 3242 | 6911 | 3669 | 46.91072204 | 53.08927796 |
| ABtoBC | AB | BC | 3536 | 7497 | 3961 | 47.16553288 | 52.83446712 |
| ECtoAF | EC | AF | 3046 | 6371 | 3325 | 47.81039083 | 52.18960917 |
| DCtoBC | DC | BC | 3500 | 7305 | 3805 | 47.91238877 | 52.08761123 |
| ABtoDC | AB | DC | 3596 | 7497 | 3901 | 47.96585301 | 52.03414699 |
| CAtoEC | CA | EC | 3425 | 7076 | 3651 | 48.40305257 | 51.59694743 |
| BCtoAF | BC | AF | 3367 | 6911 | 3544 | 48.71943279 | 51.28056721 |
| DCtoFA | DC | FA | 3580 | 7305 | 3725 | 49.00752909 | 50.99247091 |
| FAtoCA | FA | CA | 3853 | 7828 | 3975 | 49.22074604 | 50.77925396 |
| DCtoAB | DC | AB | 3596 | 7305 | 3709 | 49.22655715 | 50.77344285 |
| DCtoEC | DC | EC | 3659 | 7305 | 3646 | 50.08898015 | 49.91101985 |
| ECtoAB | EC | AB | 3206 | 6371 | 3165 | 50.32177052 | 49.67822948 |
| ECtoDB | EC | DB | 3211 | 6371 | 3160 | 50.40025114 | 49.59974886 |
| ECtoBC | EC | BC | 3215 | 6371 | 3156 | 50.46303563 | 49.53696437 |
| BCtoDC | BC | DC | 3500 | 6911 | 3411 | 50.64390103 | 49.35609897 |
| ECtoFA | EC | FA | 3232 | 6371 | 3139 | 50.72986972 | 49.27013028 |
| ABtoCA | AB | CA | 3813 | 7497 | 3684 | 50.86034414 | 49.13965586 |
| DCtoCA | DC | CA | 3720 | 7305 | 3585 | 50.92402464 | 49.07597536 |
| EFtoDC | EF | DC | 2891 | 5668 | 2777 | 51.00564573 | 48.99435427 |
| FAtoDB | FA | DB | 3995 | 7828 | 3833 | 51.03474706 | 48.96525294 |
| EFtoCA | EF | CA | 2898 | 5668 | 2770 | 51.12914608 | 48.87085392 |
| CAtoBC | CA | BC | 3625 | 7076 | 3451 | 51.2295082 | 48.7704918 |
| BCtoAB | BC | AB | 3565 | 6911 | 3346 | 51.58443062 | 48.41556938 |
| FAtoAB | FA | AB | 4043 | 7828 | 3785 | 51.64793051 | 48.35206949 |
| EFtoEC | EF | EC | 2941 | 5668 | 2727 | 51.88779111 | 48.11220889 |
| ABtoAF | AB | AF | 3905 | 7497 | 3592 | 52.08750167 | 47.91249833 |
| EFtoAB | EF | AB | 2954 | 5668 | 2714 | 52.11714891 | 47.88285109 |
| BCtoCA | BC | CA | 3625 | 6911 | 3286 | 52.45261178 | 47.54738822 |
| BAtoEF | BA | EF | 1997 | 3804 | 1807 | 52.49737119 | 47.50262881 |
| CAtoDC | CA | DC | 3720 | 7076 | 3356 | 52.57207462 | 47.42792538 |
| ECtoCA | EC | CA | 3425 | 6371 | 2946 | 53.75922147 | 46.24077853 |
| CAtoAB | CA | AB | 3813 | 7076 | 3263 | 53.88637648 | 46.11362352 |
| ABtoFA | AB | FA | 4043 | 7497 | 3454 | 53.92823796 | 46.07176204 |
| EFtoDB | EF | DB | 3060 | 5668 | 2608 | 53.98729711 | 46.01270289 |
| AFtoFA | AF | FA | 4525 | 8375 | 3850 | 54.02985075 | 45.97014925 |
| CAtoDB | CA | DB | 3826 | 7076 | 3250 | 54.0700961 | 45.9299039 |
| AFtoDB | AF | DB | 4532 | 8375 | 3843 | 54.11343284 | 45.88656716 |
| CAtoFA | CA | FA | 3853 | 7076 | 3223 | 54.45166761 | 45.54833239 |
| DCtoDB | DC | DB | 3985 | 7305 | 3320 | 54.55167693 | 45.44832307 |
| BAtoBC | BA | BC | 2077 | 3804 | 1727 | 54.60042061 | 45.39957939 |
| CAtoAF | CA | AF | 3872 | 7076 | 3204 | 54.72018089 | 45.27981911 |
| BAtoEC | BA | EC | 2084 | 3804 | 1720 | 54.78443743 | 45.21556257 |
| ABtoDB | AB | DB | 4222 | 7497 | 3275 | 56.31585968 | 43.68414032 |
| BAtoDC | BA | DC | 2149 | 3804 | 1655 | 56.49316509 | 43.50683491 |
| EFtoFA | EF | FA | 3203 | 5668 | 2465 | 56.51023289 | 43.48976711 |
| ECtoDC | EC | DC | 3659 | 6371 | 2712 | 57.43211427 | 42.56788573 |
| FAtoAF | FA | AF | 4525 | 7828 | 3303 | 57.80531426 | 42.19468574 |
| EFtoAF | EF | AF | 3318 | 5668 | 2350 | 58.53916725 | 41.46083275 |
| BCtoDB | BC | DB | 4071 | 6911 | 2840 | 58.90609174 | 41.09390826 |
| BAtoDB | BA | DB | 2250 | 3804 | 1554 | 59.14826498 | 40.85173502 |
| BAtoAF | BA | AF | 2268 | 3804 | 1536 | 59.6214511 | 40.3785489 |
| BAtoFA | BA | FA | 2277 | 3804 | 1527 | 59.85804416 | 40.14195584 |
| BAtoCA | BA | CA | 2290 | 3804 | 1514 | 60.1997897 | 39.8002103 |
| BAtoAB | BA | AB | 2515 | 3804 | 1289 | 66.11461619 | 33.88538381 |
|  |  |  |  |  | Median | 49.2236516 | 50.7763484 |
|  |  |  |  |  | Mean | 47.00602438 | 52.99397562 |
