## SupplementalTable5 for "Genetic variation in heat tolerance of the coral *Platygyra daedalea* indicates potential for adaptation to ocean warming"

**Supplemental Table 5**. A comparison of the number and percent of significant tags genotyped in the first cross (C1) versus the whole set of tags in the second cross (C2) post analysis.

| C1toC2 | Sig | Insig | NumTagsBothSig | NumSigC1InInsigC2 | TotalSigInsigC1inC2 | TotalSigC1 | PercentGenotyped | PercentNotGenotyped |
| --- | --- | --- | --- | --- | --- | --- | --- | --- |
| AFtoDB | AF | DB | 1 | 12 | 13 | 34 | 38.23529412 | 61.76470588 |
| BCtoBA | BC | BA | 4 | 24 | 28 | 69 | 40.57971014 | 59.42028986 |
| DCtoBA | DC | BA | 8 | 134 | 142 | 336 | 42.26190476 | 57.73809524 |
| DBtoBC | DB | BC | 2 | 17 | 19 | 44 | 43.18181818 | 56.81818182 |
| DBtoCA | DB | CA | 2 | 17 | 19 | 44 | 43.18181818 | 56.81818182 |
| DBtoEC | DB | EC | 1 | 18 | 19 | 44 | 43.18181818 | 56.81818182 |
| EFtoBC | EF | BC | 5 | 51 | 56 | 124 | 45.16129032 | 54.83870968 |
| DBtoFA | DB | FA | 2 | 18 | 20 | 44 | 45.45454545 | 54.54545455 |
| BCtoAF | BC | AF | 3 | 30 | 33 | 69 | 47.82608696 | 52.17391304 |
| BCtoEF | BC | EF | 5 | 28 | 33 | 69 | 47.82608696 | 52.17391304 |
| FAtoBA | FA | BA | 6 | 61 | 67 | 139 | 48.20143885 | 51.79856115 |
| ABtoBC | AB | BC | 5 | 32 | 37 | 76 | 48.68421053 | 51.31578947 |
| DCtoAF | DC | AF | 6 | 158 | 164 | 336 | 48.80952381 | 51.19047619 |
| BCtoDB | BC | DB | 2 | 32 | 34 | 69 | 49.27536232 | 50.72463768 |
| BCtoFA | BC | FA | 6 | 28 | 34 | 69 | 49.27536232 | 50.72463768 |
| FAtoDB | FA | DB | 2 | 67 | 69 | 139 | 49.64028777 | 50.35971223 |
| DBtoAB | DB | AB | 2 | 20 | 22 | 44 | 50 | 50 |
| DBtoAF | DB | AF | 1 | 21 | 22 | 44 | 50 | 50 |
| EFtoBA | EF | BA | 7 | 56 | 63 | 124 | 50.80645161 | 49.19354839 |
| EFtoDB | EF | DB | 4 | 59 | 63 | 124 | 50.80645161 | 49.19354839 |
| FAtoEC | FA | EC | 6 | 66 | 72 | 139 | 51.79856115 | 48.20143885 |
| BCtoDC | BC | DC | 10 | 26 | 36 | 69 | 52.17391304 | 47.82608696 |
| DBtoBA | DB | BA | 3 | 20 | 23 | 44 | 52.27272727 | 47.72727273 |
| FAtoEF | FA | EF | 11 | 62 | 73 | 139 | 52.51798561 | 47.48201439 |
| DCtoAB | DC | AB | 8 | 169 | 177 | 336 | 52.67857143 | 47.32142857 |
| DCtoBC | DC | BC | 10 | 168 | 178 | 336 | 52.97619048 | 47.02380952 |
| FAtoBC | FA | BC | 6 | 68 | 74 | 139 | 53.23741007 | 46.76258993 |
| DCtoFA | DC | FA | 19 | 161 | 180 | 336 | 53.57142857 | 46.42857143 |
| DCtoCA | DC | CA | 11 | 171 | 182 | 336 | 54.16666667 | 45.83333333 |
| DCtoEC | DC | EC | 11 | 172 | 183 | 336 | 54.46428571 | 45.53571429 |
| DBtoEF | DB | EF | 4 | 20 | 24 | 44 | 54.54545455 | 45.45454545 |
| CAtoDB | CA | DB | 2 | 39 | 41 | 75 | 54.66666667 | 45.33333333 |
| CAtoEF | CA | EF | 8 | 33 | 41 | 75 | 54.66666667 | 45.33333333 |
| BAtoDC | BA | DC | 8 | 45 | 53 | 95 | 55.78947368 | 44.21052632 |
| DCtoEF | DC | EF | 19 | 169 | 188 | 336 | 55.95238095 | 44.04761905 |
| FAtoAB | FA | AB | 7 | 71 | 78 | 139 | 56.11510791 | 43.88489209 |
| BCtoCA | BC | CA | 7 | 32 | 39 | 69 | 56.52173913 | 43.47826087 |
| BCtoEC | BC | EC | 5 | 34 | 39 | 69 | 56.52173913 | 43.47826087 |
| DBtoDC | DB | DC | 6 | 19 | 25 | 44 | 56.81818182 | 43.18181818 |
| FAtoCA | FA | CA | 9 | 70 | 79 | 139 | 56.83453237 | 43.16546763 |
| DCtoDB | DC | DB | 6 | 186 | 192 | 336 | 57.14285714 | 42.85714286 |
| ECtoBA | EC | BA | 4 | 40 | 44 | 77 | 57.14285714 | 42.85714286 |
| EFtoFA | EF | FA | 11 | 60 | 71 | 124 | 57.25806452 | 42.74193548 |
| ABtoDC | AB | DC | 8 | 36 | 44 | 76 | 57.89473684 | 42.10526316 |
| ABtoFA | AB | FA | 7 | 37 | 44 | 76 | 57.89473684 | 42.10526316 |
| BCtoAB | BC | AB | 5 | 35 | 40 | 69 | 57.97101449 | 42.02898551 |
| EFtoAB | EF | AB | 9 | 63 | 72 | 124 | 58.06451613 | 41.93548387 |
| CAtoBC | CA | BC | 7 | 37 | 44 | 75 | 58.66666667 | 41.33333333 |
| EFtoCA | EF | CA | 8 | 65 | 73 | 124 | 58.87096774 | 41.12903226 |
| BAtoBC | BA | BC | 4 | 52 | 56 | 95 | 58.94736842 | 41.05263158 |
| ECtoDB | EC | DB | 1 | 45 | 46 | 77 | 59.74025974 | 40.25974026 |
| ECtoFA | EC | FA | 6 | 40 | 46 | 77 | 59.74025974 | 40.25974026 |
| CAtoEC | CA | EC | 6 | 39 | 45 | 75 | 60 | 40 |
| ECtoAF | EC | AF | 4 | 43 | 47 | 77 | 61.03896104 | 38.96103896 |
| CAtoAF | CA | AF | 4 | 42 | 46 | 75 | 61.33333333 | 38.66666667 |
| AFtoBA | AF | BA | 4 | 17 | 21 | 34 | 61.76470588 | 38.23529412 |
| AFtoBC | AF | BC | 3 | 18 | 21 | 34 | 61.76470588 | 38.23529412 |
| AFtoCA | AF | CA | 4 | 17 | 21 | 34 | 61.76470588 | 38.23529412 |
| AFtoDC | AF | DC | 6 | 15 | 21 | 34 | 61.76470588 | 38.23529412 |
| AFtoEC | AF | EC | 4 | 17 | 21 | 34 | 61.76470588 | 38.23529412 |
| ABtoCA | AB | CA | 9 | 38 | 47 | 76 | 61.84210526 | 38.15789474 |
| ABtoDB | AB | DB | 2 | 45 | 47 | 76 | 61.84210526 | 38.15789474 |
| ABtoEF | AB | EF | 9 | 38 | 47 | 76 | 61.84210526 | 38.15789474 |
| BAtoEF | BA | EF | 7 | 52 | 59 | 95 | 62.10526316 | 37.89473684 |
| BAtoFA | BA | FA | 6 | 53 | 59 | 95 | 62.10526316 | 37.89473684 |
| ECtoAB | EC | AB | 5 | 43 | 48 | 77 | 62.33766234 | 37.66233766 |
| ABtoEC | AB | EC | 5 | 43 | 48 | 76 | 63.15789474 | 36.84210526 |
| BAtoEC | BA | EC | 4 | 56 | 60 | 95 | 63.15789474 | 36.84210526 |
| BAtoDB | BA | DB | 3 | 58 | 61 | 95 | 64.21052632 | 35.78947368 |
| ABtoAF | AB | AF | 7 | 42 | 49 | 76 | 64.47368421 | 35.52631579 |
| ABtoBA | AB | BA | 10 | 39 | 49 | 76 | 64.47368421 | 35.52631579 |
| ECtoBC | EC | BC | 5 | 45 | 50 | 77 | 64.93506494 | 35.06493506 |
| EFtoDC | EF | DC | 19 | 62 | 81 | 124 | 65.32258065 | 34.67741935 |
| CAtoBA | CA | BA | 11 | 38 | 49 | 75 | 65.33333333 | 34.66666667 |
| EFtoEC | EF | EC | 14 | 68 | 82 | 124 | 66.12903226 | 33.87096774 |
| FAtoAF | FA | AF | 8 | 84 | 92 | 139 | 66.18705036 | 33.81294964 |
| FAtoDC | FA | DC | 19 | 73 | 92 | 139 | 66.18705036 | 33.81294964 |
| BAtoAF | BA | AF | 4 | 59 | 63 | 95 | 66.31578947 | 33.68421053 |
| EFtoAF | EF | AF | 10 | 73 | 83 | 124 | 66.93548387 | 33.06451613 |
| ECtoCA | EC | CA | 6 | 46 | 52 | 77 | 67.53246753 | 32.46753247 |
| CAtoDC | CA | DC | 11 | 40 | 51 | 75 | 68 | 32 |
| CAtoFA | CA | FA | 9 | 42 | 51 | 75 | 68 | 32 |
| BAtoAB | BA | AB | 10 | 55 | 65 | 95 | 68.42105263 | 31.57894737 |
| AFtoEF | AF | EF | 10 | 14 | 24 | 34 | 70.58823529 | 29.41176471 |
| CAtoAB | CA | AB | 9 | 45 | 54 | 75 | 72 | 28 |
| ECtoDC | EC | DC | 11 | 45 | 56 | 77 | 72.72727273 | 27.27272727 |
| ECtoEF | EC | EF | 14 | 42 | 56 | 77 | 72.72727273 | 27.27272727 |
| BAtoCA | BA | CA | 11 | 62 | 73 | 95 | 76.84210526 | 23.15789474 |
| AFtoAB | AF | AB | 7 | 22 | 29 | 34 | 85.29411765 | 14.70588235 |
| AFtoFA | AF | FA | 8 | 22 | 30 | 34 | 88.23529412 | 11.76470588 |
|  |  |  |  |  |  | Median | 57.93287567 | 42.06712433 |
|  |  |  |  |  |  | Mean | 58.22745184 | 41.77254816 |
