## SupplementalTable6 for "Genetic variation in heat tolerance of the coral *Platygyra daedalea* indicates potential for adaptation to ocean warming"

**Supplemental Table 6**. For each tag, it was determined to be insignificant or significant in each cross. There is a count of the number of crosses the tag is found to be significant, the number of crosses in which it was found to be insignificant, and the number of crosses the tag was not genotyped.

| Tag | AB | AF | BA | BC | CA | DB | DC | EC | EF | FA | Num  Crosses Sig | Num Crosses  Insig | Not Geno-typed |
| --- | --- | --- | --- | --- | --- | --- | --- | --- | --- | --- | --- | --- | --- |
| denovoLocus34739 | significant | significant | not genotyped | significant | significant | insignificant | significant | not genotyped | significant | significant | 7 | 1 | 2 |
| denovoLocus44191 | significant | significant | significant | not genotyped | significant | significant | not genotyped | not genotyped | significant | significant | 7 | 0 | 3 |
| denovoLocus9359 | significant | insignificant | insignificant | significant | significant | insignificant | significant | not genotyped | significant | significant | 6 | 3 | 1 |
| denovoLocus121909 | significant | insignificant | significant | insignificant | significant | insignificant | insignificant | significant | not genotyped | significant | 5 | 4 | 1 |
| denovoLocus22543 | not genotyped | not genotyped | significant | insignificant | significant | insignificant | insignificant | significant | significant | significant | 5 | 3 | 2 |
| denovoLocus93322 | significant | significant | not genotyped | insignificant | insignificant | insignificant | significant | significant | significant | not genotyped | 5 | 3 | 2 |
| denovoLocus1291 | significant | insignificant | significant | significant | not genotyped | significant | not genotyped | not genotyped | significant | not genotyped | 5 | 1 | 4 |
| denovoLocus18723 | insignificant | insignificant | significant | not genotyped | insignificant | significant | significant | significant | insignificant | insignificant | 4 | 5 | 1 |
| denovoLocus13062 | insignificant | insignificant | not genotyped | not genotyped | significant | insignificant | insignificant | significant | significant | significant | 4 | 4 | 2 |
| denovoLocus36225 | significant | significant | not genotyped | insignificant | insignificant | insignificant | not genotyped | significant | significant | insignificant | 4 | 4 | 2 |
| denovoLocus29413 | not genotyped | insignificant | not genotyped | significant | not genotyped | insignificant | significant | insignificant | significant | significant | 4 | 3 | 3 |
| denovoLocus51123 | significant | significant | not genotyped | not genotyped | not genotyped | not genotyped | significant | not genotyped | significant | insignificant | 4 | 1 | 5 |
| denovoLocus18867 | insignificant | insignificant | insignificant | significant | insignificant | insignificant | significant | insignificant | insignificant | significant | 3 | 7 | 0 |
| denovoLocus10613 | insignificant | insignificant | insignificant | insignificant | significant | insignificant | significant | significant | not genotyped | insignificant | 3 | 6 | 1 |
| denovoLocus13278 | insignificant | insignificant | insignificant | not genotyped | insignificant | insignificant | significant | significant | significant | insignificant | 3 | 6 | 1 |
| denovoLocus6154 | insignificant | insignificant | significant | insignificant | insignificant | insignificant | significant | insignificant | not genotyped | significant | 3 | 6 | 1 |
| denovoLocus30715 | not genotyped | insignificant | insignificant | insignificant | insignificant | insignificant | significant | significant | not genotyped | significant | 3 | 5 | 2 |
| denovoLocus104310 | not genotyped | not genotyped | insignificant | significant | insignificant | insignificant | significant | not genotyped | significant | insignificant | 3 | 4 | 3 |
| denovoLocus13680 | insignificant | insignificant | insignificant | not genotyped | significant | significant | insignificant | not genotyped | significant | not genotyped | 3 | 4 | 3 |
| denovoLocus49341 | insignificant | significant | insignificant | insignificant | significant | not genotyped | not genotyped | insignificant | not genotyped | significant | 3 | 4 | 3 |
| denovoLocus75321 | insignificant | insignificant | not genotyped | significant | significant | insignificant | significant | insignificant | not genotyped | not genotyped | 3 | 4 | 3 |
| denovoLocus8753 | insignificant | insignificant | insignificant | significant | significant | not genotyped | not genotyped | significant | not genotyped | insignificant | 3 | 4 | 3 |
| denovoLocus1685 | not genotyped | insignificant | significant | significant | significant | insignificant | not genotyped | not genotyped | not genotyped | insignificant | 3 | 3 | 4 |
| denovoLocus24499 | insignificant | significant | insignificant | insignificant | not genotyped | not genotyped | significant | not genotyped | significant | not genotyped | 3 | 3 | 4 |
| denovoLocus32940 | insignificant | significant | insignificant | not genotyped | not genotyped | not genotyped | significant | insignificant | significant | not genotyped | 3 | 3 | 4 |
| denovoLocus52938 | not genotyped | significant | not genotyped | insignificant | not genotyped | insignificant | insignificant | not genotyped | significant | significant | 3 | 3 | 4 |
| denovoLocus6164 | significant | insignificant | significant | not genotyped | significant | insignificant | insignificant | not genotyped | not genotyped | not genotyped | 3 | 3 | 4 |
| denovoLocus14265 | significant | not genotyped | not genotyped | insignificant | not genotyped | insignificant | not genotyped | significant | significant | not genotyped | 3 | 2 | 5 |
| denovoLocus2572 | not genotyped | insignificant | not genotyped | not genotyped | significant | not genotyped | significant | insignificant | not genotyped | significant | 3 | 2 | 5 |
| denovoLocus6036 | significant | not genotyped | significant | significant | insignificant | not genotyped | not genotyped | insignificant | not genotyped | not genotyped | 3 | 2 | 5 |
| denovoLocus16627 | not genotyped | not genotyped | not genotyped | significant | not genotyped | significant | significant | not genotyped | insignificant | not genotyped | 3 | 1 | 6 |
| denovoLocus22915 | not genotyped | significant | significant | not genotyped | significant | not genotyped | not genotyped | not genotyped | not genotyped | insignificant | 3 | 1 | 6 |
| denovoLocus38500 | significant | not genotyped | significant | not genotyped | not genotyped | not genotyped | not genotyped | insignificant | significant | not genotyped | 3 | 1 | 6 |
| denovoLocus58853 | not genotyped | significant | significant | not genotyped | not genotyped | not genotyped | not genotyped | not genotyped | not genotyped | significant | 3 | 0 | 7 |
| denovoLocus85411 | significant | not genotyped | not genotyped | not genotyped | not genotyped | not genotyped | not genotyped | significant | not genotyped | significant | 3 | 0 | 7 |
| denovoLocus14433 | insignificant | insignificant | significant | insignificant | insignificant | insignificant | significant | insignificant | insignificant | insignificant | 2 | 8 | 0 |
| denovoLocus26881 | significant | insignificant | insignificant | insignificant | insignificant | insignificant | insignificant | insignificant | insignificant | significant | 2 | 8 | 0 |
| denovoLocus64963 | insignificant | insignificant | insignificant | insignificant | insignificant | insignificant | significant | insignificant | insignificant | significant | 2 | 8 | 0 |
| denovoLocus16027 | insignificant | insignificant | insignificant | insignificant | insignificant | significant | significant | insignificant | insignificant | not genotyped | 2 | 7 | 1 |
| denovoLocus25366 | insignificant | insignificant | insignificant | insignificant | insignificant | insignificant | significant | insignificant | not genotyped | significant | 2 | 7 | 1 |
| denovoLocus49865 | insignificant | insignificant | significant | not genotyped | insignificant | insignificant | significant | insignificant | insignificant | insignificant | 2 | 7 | 1 |
| denovoLocus6620 | insignificant | insignificant | not genotyped | insignificant | insignificant | insignificant | significant | insignificant | significant | insignificant | 2 | 7 | 1 |
| denovoLocus68 | insignificant | not genotyped | insignificant | insignificant | insignificant | insignificant | significant | significant | insignificant | insignificant | 2 | 7 | 1 |
| denovoLocus6969 | insignificant | not genotyped | significant | insignificant | insignificant | insignificant | significant | insignificant | insignificant | insignificant | 2 | 7 | 1 |
| denovoLocus89683 | insignificant | insignificant | significant | not genotyped | significant | insignificant | insignificant | insignificant | insignificant | insignificant | 2 | 7 | 1 |
| denovoLocus10129 | insignificant | insignificant | insignificant | insignificant | not genotyped | not genotyped | insignificant | significant | significant | insignificant | 2 | 6 | 2 |
| denovoLocus105818 | insignificant | insignificant | insignificant | insignificant | insignificant | insignificant | not genotyped | not genotyped | significant | significant | 2 | 6 | 2 |
| denovoLocus10860 | insignificant | not genotyped | insignificant | insignificant | insignificant | insignificant | significant | not genotyped | significant | insignificant | 2 | 6 | 2 |
| denovoLocus11595 | insignificant | insignificant | insignificant | not genotyped | not genotyped | insignificant | significant | insignificant | insignificant | significant | 2 | 6 | 2 |
| denovoLocus14833 | significant | not genotyped | significant | insignificant | insignificant | insignificant | insignificant | not genotyped | insignificant | insignificant | 2 | 6 | 2 |
| denovoLocus15476 | insignificant | insignificant | insignificant | significant | not genotyped | insignificant | significant | not genotyped | insignificant | insignificant | 2 | 6 | 2 |
| denovoLocus17167 | insignificant | insignificant | significant | insignificant | significant | insignificant | not genotyped | insignificant | not genotyped | insignificant | 2 | 6 | 2 |
| denovoLocus20019 | insignificant | insignificant | insignificant | insignificant | insignificant | not genotyped | significant | insignificant | significant | not genotyped | 2 | 6 | 2 |
| denovoLocus23162 | insignificant | insignificant | not genotyped | insignificant | not genotyped | insignificant | significant | insignificant | significant | insignificant | 2 | 6 | 2 |
| denovoLocus311 | insignificant | insignificant | insignificant | insignificant | insignificant | not genotyped | not genotyped | significant | significant | insignificant | 2 | 6 | 2 |
| denovoLocus31423 | insignificant | insignificant | not genotyped | insignificant | insignificant | not genotyped | significant | insignificant | insignificant | significant | 2 | 6 | 2 |
| denovoLocus44692 | insignificant | insignificant | insignificant | not genotyped | insignificant | insignificant | not genotyped | significant | significant | insignificant | 2 | 6 | 2 |
| denovoLocus48424 | significant | insignificant | insignificant | insignificant | insignificant | not genotyped | not genotyped | insignificant | insignificant | significant | 2 | 6 | 2 |
| denovoLocus8450 | insignificant | insignificant | insignificant | not genotyped | insignificant | not genotyped | significant | insignificant | significant | insignificant | 2 | 6 | 2 |
| denovoLocus111812 | insignificant | not genotyped | not genotyped | insignificant | insignificant | significant | insignificant | not genotyped | significant | insignificant | 2 | 5 | 3 |
| denovoLocus13663 | insignificant | insignificant | insignificant | insignificant | not genotyped | insignificant | significant | not genotyped | significant | not genotyped | 2 | 5 | 3 |
| denovoLocus17994 | significant | not genotyped | insignificant | not genotyped | insignificant | insignificant | significant | insignificant | insignificant | not genotyped | 2 | 5 | 3 |
| denovoLocus19325 | significant | not genotyped | significant | insignificant | insignificant | insignificant | insignificant | not genotyped | not genotyped | insignificant | 2 | 5 | 3 |
| denovoLocus25423 | not genotyped | insignificant | insignificant | not genotyped | insignificant | insignificant | significant | insignificant | not genotyped | significant | 2 | 5 | 3 |
| denovoLocus27975 | insignificant | insignificant | significant | insignificant | insignificant | not genotyped | not genotyped | insignificant | not genotyped | significant | 2 | 5 | 3 |
| denovoLocus31095 | insignificant | insignificant | not genotyped | not genotyped | insignificant | not genotyped | insignificant | significant | significant | insignificant | 2 | 5 | 3 |
| denovoLocus31194 | insignificant | insignificant | not genotyped | insignificant | significant | not genotyped | insignificant | significant | not genotyped | insignificant | 2 | 5 | 3 |
| denovoLocus31648 | significant | insignificant | significant | insignificant | not genotyped | insignificant | insignificant | insignificant | not genotyped | not genotyped | 2 | 5 | 3 |
| denovoLocus32448 | insignificant | not genotyped | significant | insignificant | insignificant | insignificant | insignificant | not genotyped | significant | not genotyped | 2 | 5 | 3 |
| denovoLocus38089 | not genotyped | insignificant | not genotyped | significant | insignificant | insignificant | insignificant | significant | not genotyped | insignificant | 2 | 5 | 3 |
| denovoLocus44384 | not genotyped | insignificant | not genotyped | not genotyped | insignificant | insignificant | insignificant | significant | significant | insignificant | 2 | 5 | 3 |
| denovoLocus49247 | insignificant | not genotyped | not genotyped | insignificant | insignificant | not genotyped | significant | insignificant | insignificant | significant | 2 | 5 | 3 |
| denovoLocus49606 | insignificant | not genotyped | insignificant | not genotyped | insignificant | not genotyped | significant | insignificant | significant | insignificant | 2 | 5 | 3 |
| denovoLocus57700 | insignificant | not genotyped | significant | not genotyped | insignificant | insignificant | significant | not genotyped | insignificant | insignificant | 2 | 5 | 3 |
| denovoLocus6000 | not genotyped | significant | insignificant | significant | not genotyped | insignificant | not genotyped | insignificant | insignificant | insignificant | 2 | 5 | 3 |
| denovoLocus7920 | insignificant | not genotyped | not genotyped | significant | insignificant | not genotyped | insignificant | insignificant | insignificant | significant | 2 | 5 | 3 |
| denovoLocus79497 | insignificant | significant | insignificant | not genotyped | not genotyped | insignificant | insignificant | significant | not genotyped | insignificant | 2 | 5 | 3 |
| denovoLocus81929 | insignificant | insignificant | insignificant | significant | significant | not genotyped | not genotyped | insignificant | insignificant | not genotyped | 2 | 5 | 3 |
| denovoLocus84456 | insignificant | insignificant | not genotyped | insignificant | not genotyped | not genotyped | significant | insignificant | insignificant | significant | 2 | 5 | 3 |
| denovoLocus85726 | not genotyped | insignificant | not genotyped | insignificant | insignificant | significant | significant | insignificant | not genotyped | insignificant | 2 | 5 | 3 |
| denovoLocus8626 | significant | not genotyped | insignificant | insignificant | significant | not genotyped | insignificant | insignificant | insignificant | not genotyped | 2 | 5 | 3 |
| denovoLocus960 | insignificant | significant | insignificant | insignificant | not genotyped | not genotyped | insignificant | significant | not genotyped | insignificant | 2 | 5 | 3 |
| denovoLocus10133 | insignificant | insignificant | not genotyped | not genotyped | insignificant | not genotyped | significant | significant | not genotyped | insignificant | 2 | 4 | 4 |
| denovoLocus11443 | significant | insignificant | significant | not genotyped | insignificant | insignificant | not genotyped | insignificant | not genotyped | not genotyped | 2 | 4 | 4 |
| denovoLocus12018 | significant | insignificant | insignificant | not genotyped | not genotyped | not genotyped | significant | insignificant | not genotyped | insignificant | 2 | 4 | 4 |
| denovoLocus13835 | insignificant | not genotyped | insignificant | not genotyped | not genotyped | insignificant | insignificant | significant | significant | not genotyped | 2 | 4 | 4 |
| denovoLocus1416 | insignificant | significant | significant | insignificant | insignificant | not genotyped | not genotyped | insignificant | not genotyped | not genotyped | 2 | 4 | 4 |
| denovoLocus14360 | not genotyped | insignificant | not genotyped | not genotyped | not genotyped | insignificant | insignificant | significant | insignificant | significant | 2 | 4 | 4 |
| denovoLocus1480 | insignificant | insignificant | insignificant | not genotyped | insignificant | not genotyped | not genotyped | not genotyped | significant | significant | 2 | 4 | 4 |
| denovoLocus22267 | significant | not genotyped | insignificant | significant | not genotyped | insignificant | insignificant | insignificant | not genotyped | not genotyped | 2 | 4 | 4 |
| denovoLocus24558 | insignificant | insignificant | not genotyped | not genotyped | significant | insignificant | significant | not genotyped | insignificant | not genotyped | 2 | 4 | 4 |
| denovoLocus3033 | insignificant | not genotyped | not genotyped | significant | not genotyped | insignificant | significant | insignificant | insignificant | not genotyped | 2 | 4 | 4 |
| denovoLocus30716 | insignificant | not genotyped | insignificant | not genotyped | significant | insignificant | not genotyped | insignificant | significant | not genotyped | 2 | 4 | 4 |
| denovoLocus3805 | insignificant | not genotyped | insignificant | not genotyped | insignificant | insignificant | not genotyped | significant | significant | not genotyped | 2 | 4 | 4 |
| denovoLocus38700 | significant | insignificant | not genotyped | not genotyped | significant | not genotyped | insignificant | insignificant | not genotyped | insignificant | 2 | 4 | 4 |
| denovoLocus39998 | insignificant | insignificant | insignificant | not genotyped | not genotyped | insignificant | significant | significant | not genotyped | not genotyped | 2 | 4 | 4 |
| denovoLocus41205 | not genotyped | insignificant | not genotyped | not genotyped | insignificant | not genotyped | significant | insignificant | insignificant | significant | 2 | 4 | 4 |
| denovoLocus45468 | not genotyped | insignificant | insignificant | not genotyped | not genotyped | insignificant | significant | significant | insignificant | not genotyped | 2 | 4 | 4 |
| denovoLocus48746 | insignificant | insignificant | significant | not genotyped | not genotyped | not genotyped | significant | insignificant | insignificant | not genotyped | 2 | 4 | 4 |
| denovoLocus60119 | insignificant | not genotyped | not genotyped | significant | significant | insignificant | insignificant | not genotyped | not genotyped | insignificant | 2 | 4 | 4 |
| denovoLocus60238 | not genotyped | significant | insignificant | insignificant | not genotyped | not genotyped | insignificant | insignificant | not genotyped | significant | 2 | 4 | 4 |
| denovoLocus6041 | not genotyped | not genotyped | significant | insignificant | insignificant | insignificant | not genotyped | significant | not genotyped | insignificant | 2 | 4 | 4 |
| denovoLocus62710 | not genotyped | insignificant | insignificant | not genotyped | significant | not genotyped | significant | insignificant | insignificant | not genotyped | 2 | 4 | 4 |
| denovoLocus67017 | not genotyped | not genotyped | not genotyped | insignificant | insignificant | not genotyped | insignificant | insignificant | significant | significant | 2 | 4 | 4 |
| denovoLocus80641 | not genotyped | insignificant | not genotyped | not genotyped | significant | insignificant | significant | insignificant | not genotyped | insignificant | 2 | 4 | 4 |
| denovoLocus94104 | insignificant | insignificant | significant | insignificant | not genotyped | not genotyped | not genotyped | insignificant | significant | not genotyped | 2 | 4 | 4 |
| denovoLocus1110 | not genotyped | insignificant | not genotyped | not genotyped | insignificant | not genotyped | significant | insignificant | not genotyped | significant | 2 | 3 | 5 |
| denovoLocus111991 | not genotyped | not genotyped | not genotyped | not genotyped | insignificant | insignificant | significant | insignificant | significant | not genotyped | 2 | 3 | 5 |
| denovoLocus13732 | not genotyped | not genotyped | insignificant | not genotyped | insignificant | not genotyped | significant | significant | not genotyped | insignificant | 2 | 3 | 5 |
| denovoLocus15899 | not genotyped | insignificant | significant | insignificant | significant | not genotyped | not genotyped | not genotyped | not genotyped | insignificant | 2 | 3 | 5 |
| denovoLocus1668 | significant | not genotyped | not genotyped | not genotyped | significant | insignificant | insignificant | insignificant | not genotyped | not genotyped | 2 | 3 | 5 |
| denovoLocus19024 | not genotyped | insignificant | not genotyped | not genotyped | not genotyped | not genotyped | insignificant | insignificant | significant | significant | 2 | 3 | 5 |
| denovoLocus21802 | not genotyped | significant | not genotyped | insignificant | insignificant | not genotyped | significant | insignificant | not genotyped | not genotyped | 2 | 3 | 5 |
| denovoLocus23425 | not genotyped | not genotyped | significant | insignificant | significant | not genotyped | insignificant | not genotyped | insignificant | not genotyped | 2 | 3 | 5 |
| denovoLocus24254 | not genotyped | not genotyped | significant | insignificant | significant | insignificant | insignificant | not genotyped | not genotyped | not genotyped | 2 | 3 | 5 |
| denovoLocus25267 | insignificant | significant | not genotyped | significant | insignificant | not genotyped | not genotyped | insignificant | not genotyped | not genotyped | 2 | 3 | 5 |
| denovoLocus2899 | insignificant | not genotyped | not genotyped | significant | not genotyped | not genotyped | significant | insignificant | not genotyped | insignificant | 2 | 3 | 5 |
| denovoLocus32401 | insignificant | not genotyped | not genotyped | significant | insignificant | not genotyped | not genotyped | significant | insignificant | not genotyped | 2 | 3 | 5 |
| denovoLocus32998 | not genotyped | insignificant | insignificant | not genotyped | not genotyped | significant | not genotyped | insignificant | not genotyped | significant | 2 | 3 | 5 |
| denovoLocus36935 | not genotyped | not genotyped | insignificant | insignificant | not genotyped | insignificant | significant | not genotyped | not genotyped | significant | 2 | 3 | 5 |
| denovoLocus41168 | not genotyped | not genotyped | not genotyped | not genotyped | not genotyped | insignificant | insignificant | significant | significant | insignificant | 2 | 3 | 5 |
| denovoLocus44176 | insignificant | not genotyped | insignificant | not genotyped | insignificant | significant | significant | not genotyped | not genotyped | not genotyped | 2 | 3 | 5 |
| denovoLocus5881 | not genotyped | not genotyped | insignificant | significant | not genotyped | insignificant | not genotyped | significant | insignificant | not genotyped | 2 | 3 | 5 |
| denovoLocus73508 | significant | not genotyped | not genotyped | insignificant | not genotyped | insignificant | significant | insignificant | not genotyped | not genotyped | 2 | 3 | 5 |
| denovoLocus83186 | significant | significant | insignificant | not genotyped | not genotyped | insignificant | not genotyped | insignificant | not genotyped | not genotyped | 2 | 3 | 5 |
| denovoLocus8751 | not genotyped | not genotyped | not genotyped | not genotyped | significant | insignificant | insignificant | insignificant | significant | not genotyped | 2 | 3 | 5 |
| denovoLocus9237 | not genotyped | significant | insignificant | insignificant | insignificant | not genotyped | not genotyped | not genotyped | not genotyped | significant | 2 | 3 | 5 |
| denovoLocus9987 | insignificant | not genotyped | not genotyped | not genotyped | significant | insignificant | significant | not genotyped | not genotyped | insignificant | 2 | 3 | 5 |
| denovoLocus11764 | not genotyped | not genotyped | significant | insignificant | not genotyped | not genotyped | significant | not genotyped | insignificant | not genotyped | 2 | 2 | 6 |
| denovoLocus12258 | significant | not genotyped | not genotyped | not genotyped | significant | insignificant | not genotyped | not genotyped | insignificant | not genotyped | 2 | 2 | 6 |
| denovoLocus16414 | not genotyped | not genotyped | not genotyped | not genotyped | insignificant | not genotyped | significant | significant | not genotyped | insignificant | 2 | 2 | 6 |
| denovoLocus19951 | not genotyped | not genotyped | not genotyped | not genotyped | significant | not genotyped | significant | insignificant | insignificant | not genotyped | 2 | 2 | 6 |
| denovoLocus20580 | not genotyped | insignificant | significant | significant | not genotyped | not genotyped | not genotyped | not genotyped | not genotyped | insignificant | 2 | 2 | 6 |
| denovoLocus22104 | insignificant | not genotyped | not genotyped | significant | not genotyped | insignificant | not genotyped | not genotyped | not genotyped | significant | 2 | 2 | 6 |
| denovoLocus24976 | insignificant | insignificant | not genotyped | not genotyped | not genotyped | not genotyped | significant | not genotyped | significant | not genotyped | 2 | 2 | 6 |
| denovoLocus29455 | insignificant | not genotyped | not genotyped | insignificant | significant | not genotyped | not genotyped | not genotyped | not genotyped | significant | 2 | 2 | 6 |
| denovoLocus33479 | insignificant | significant | not genotyped | not genotyped | not genotyped | insignificant | not genotyped | not genotyped | significant | not genotyped | 2 | 2 | 6 |
| denovoLocus44024 | insignificant | significant | not genotyped | not genotyped | not genotyped | not genotyped | insignificant | not genotyped | not genotyped | significant | 2 | 2 | 6 |
| denovoLocus52651 | not genotyped | insignificant | not genotyped | insignificant | not genotyped | not genotyped | significant | not genotyped | significant | not genotyped | 2 | 2 | 6 |
| denovoLocus53380 | significant | significant | not genotyped | not genotyped | insignificant | not genotyped | not genotyped | insignificant | not genotyped | not genotyped | 2 | 2 | 6 |
| denovoLocus55443 | not genotyped | not genotyped | not genotyped | insignificant | insignificant | significant | significant | not genotyped | not genotyped | not genotyped | 2 | 2 | 6 |
| denovoLocus105160 | not genotyped | not genotyped | not genotyped | not genotyped | not genotyped | insignificant | significant | not genotyped | not genotyped | significant | 2 | 1 | 7 |
| denovoLocus23712 | not genotyped | not genotyped | not genotyped | not genotyped | not genotyped | not genotyped | significant | insignificant | not genotyped | significant | 2 | 1 | 7 |
| denovoLocus47367 | not genotyped | significant | not genotyped | not genotyped | not genotyped | not genotyped | not genotyped | not genotyped | significant | insignificant | 2 | 1 | 7 |
| denovoLocus71438 | significant | not genotyped | not genotyped | not genotyped | insignificant | not genotyped | significant | not genotyped | not genotyped | not genotyped | 2 | 1 | 7 |
| denovoLocus27225 | not genotyped | not genotyped | significant | not genotyped | not genotyped | not genotyped | not genotyped | not genotyped | significant | not genotyped | 2 | 0 | 8 |
| denovoLocus46004 | not genotyped | not genotyped | not genotyped | not genotyped | significant | not genotyped | significant | not genotyped | not genotyped | not genotyped | 2 | 0 | 8 |
| denovoLocus49498 | not genotyped | not genotyped | not genotyped | significant | not genotyped | not genotyped | not genotyped | significant | not genotyped | not genotyped | 2 | 0 | 8 |
| denovoLocus21677 | insignificant | insignificant | insignificant | insignificant | insignificant | insignificant | insignificant | insignificant | significant | insignificant | 1 | 9 | 0 |
| denovoLocus22190 | insignificant | insignificant | insignificant | insignificant | insignificant | insignificant | insignificant | significant | insignificant | insignificant | 1 | 9 | 0 |
| denovoLocus28019 | insignificant | insignificant | insignificant | insignificant | insignificant | insignificant | significant | insignificant | insignificant | insignificant | 1 | 9 | 0 |
| denovoLocus28223 | insignificant | insignificant | insignificant | insignificant | insignificant | insignificant | insignificant | insignificant | insignificant | significant | 1 | 9 | 0 |
| denovoLocus4978 | insignificant | significant | insignificant | insignificant | insignificant | insignificant | insignificant | insignificant | insignificant | insignificant | 1 | 9 | 0 |
| denovoLocus5212 | insignificant | insignificant | insignificant | insignificant | insignificant | insignificant | insignificant | insignificant | insignificant | significant | 1 | 9 | 0 |
| denovoLocus55411 | insignificant | insignificant | significant | insignificant | insignificant | insignificant | insignificant | insignificant | insignificant | insignificant | 1 | 9 | 0 |
| denovoLocus72790 | insignificant | insignificant | significant | insignificant | insignificant | insignificant | insignificant | insignificant | insignificant | insignificant | 1 | 9 | 0 |
| denovoLocus92 | insignificant | insignificant | insignificant | insignificant | insignificant | insignificant | insignificant | significant | insignificant | insignificant | 1 | 9 | 0 |
| denovoLocus10104 | insignificant | insignificant | insignificant | insignificant | insignificant | insignificant | significant | not genotyped | insignificant | insignificant | 1 | 8 | 1 |
| denovoLocus10598 | insignificant | insignificant | not genotyped | insignificant | insignificant | insignificant | significant | insignificant | insignificant | insignificant | 1 | 8 | 1 |
| denovoLocus10931 | not genotyped | insignificant | insignificant | insignificant | insignificant | insignificant | insignificant | insignificant | significant | insignificant | 1 | 8 | 1 |
| denovoLocus10935 | insignificant | insignificant | insignificant | insignificant | insignificant | insignificant | not genotyped | significant | insignificant | insignificant | 1 | 8 | 1 |
| denovoLocus11201 | significant | insignificant | insignificant | insignificant | insignificant | not genotyped | insignificant | insignificant | insignificant | insignificant | 1 | 8 | 1 |
| denovoLocus12913 | insignificant | not genotyped | insignificant | insignificant | insignificant | significant | insignificant | insignificant | insignificant | insignificant | 1 | 8 | 1 |
| denovoLocus17755 | insignificant | insignificant | insignificant | not genotyped | insignificant | insignificant | insignificant | insignificant | significant | insignificant | 1 | 8 | 1 |
| denovoLocus1895 | insignificant | insignificant | insignificant | insignificant | insignificant | insignificant | insignificant | significant | insignificant | not genotyped | 1 | 8 | 1 |
| denovoLocus19679 | insignificant | insignificant | insignificant | insignificant | insignificant | not genotyped | significant | insignificant | insignificant | insignificant | 1 | 8 | 1 |
| denovoLocus19925 | insignificant | insignificant | significant | insignificant | insignificant | insignificant | insignificant | insignificant | not genotyped | insignificant | 1 | 8 | 1 |
| denovoLocus19961 | not genotyped | insignificant | insignificant | insignificant | insignificant | insignificant | significant | insignificant | insignificant | insignificant | 1 | 8 | 1 |
| denovoLocus2068 | insignificant | insignificant | insignificant | insignificant | insignificant | insignificant | insignificant | insignificant | significant | not genotyped | 1 | 8 | 1 |
| denovoLocus23972 | not genotyped | insignificant | insignificant | insignificant | insignificant | insignificant | significant | insignificant | insignificant | insignificant | 1 | 8 | 1 |
| denovoLocus24589 | insignificant | insignificant | not genotyped | insignificant | insignificant | insignificant | significant | insignificant | insignificant | insignificant | 1 | 8 | 1 |
| denovoLocus25904 | insignificant | insignificant | significant | insignificant | not genotyped | insignificant | insignificant | insignificant | insignificant | insignificant | 1 | 8 | 1 |
| denovoLocus2678 | insignificant | insignificant | significant | insignificant | insignificant | insignificant | insignificant | insignificant | not genotyped | insignificant | 1 | 8 | 1 |
| denovoLocus27065 | insignificant | insignificant | insignificant | insignificant | insignificant | insignificant | insignificant | insignificant | not genotyped | significant | 1 | 8 | 1 |
| denovoLocus27356 | insignificant | not genotyped | significant | insignificant | insignificant | insignificant | insignificant | insignificant | insignificant | insignificant | 1 | 8 | 1 |
| denovoLocus2736 | insignificant | not genotyped | insignificant | insignificant | insignificant | insignificant | significant | insignificant | insignificant | insignificant | 1 | 8 | 1 |
| denovoLocus28616 | insignificant | insignificant | insignificant | insignificant | insignificant | significant | not genotyped | insignificant | insignificant | insignificant | 1 | 8 | 1 |
| denovoLocus31843 | not genotyped | insignificant | insignificant | insignificant | insignificant | insignificant | insignificant | insignificant | insignificant | significant | 1 | 8 | 1 |
| denovoLocus32126 | insignificant | insignificant | insignificant | insignificant | not genotyped | insignificant | significant | insignificant | insignificant | insignificant | 1 | 8 | 1 |
| denovoLocus32151 | insignificant | insignificant | insignificant | insignificant | not genotyped | insignificant | insignificant | insignificant | insignificant | significant | 1 | 8 | 1 |
| denovoLocus33180 | not genotyped | insignificant | insignificant | insignificant | insignificant | insignificant | significant | insignificant | insignificant | insignificant | 1 | 8 | 1 |
| denovoLocus33777 | insignificant | insignificant | insignificant | significant | insignificant | insignificant | insignificant | insignificant | not genotyped | insignificant | 1 | 8 | 1 |
| denovoLocus3449 | insignificant | insignificant | insignificant | insignificant | insignificant | insignificant | significant | insignificant | insignificant | not genotyped | 1 | 8 | 1 |
| denovoLocus42495 | insignificant | insignificant | insignificant | insignificant | insignificant | insignificant | insignificant | not genotyped | insignificant | significant | 1 | 8 | 1 |
| denovoLocus47489 | insignificant | insignificant | insignificant | insignificant | insignificant | insignificant | not genotyped | significant | insignificant | insignificant | 1 | 8 | 1 |
| denovoLocus4855 | insignificant | insignificant | insignificant | insignificant | insignificant | insignificant | significant | insignificant | insignificant | not genotyped | 1 | 8 | 1 |
| denovoLocus49486 | insignificant | insignificant | insignificant | insignificant | insignificant | not genotyped | insignificant | insignificant | insignificant | significant | 1 | 8 | 1 |
| denovoLocus51485 | not genotyped | insignificant | insignificant | insignificant | insignificant | insignificant | insignificant | insignificant | insignificant | significant | 1 | 8 | 1 |
| denovoLocus51748 | insignificant | insignificant | insignificant | insignificant | significant | insignificant | not genotyped | insignificant | insignificant | insignificant | 1 | 8 | 1 |
| denovoLocus53513 | significant | insignificant | insignificant | insignificant | insignificant | insignificant | insignificant | insignificant | insignificant | not genotyped | 1 | 8 | 1 |
| denovoLocus54540 | insignificant | insignificant | insignificant | insignificant | not genotyped | insignificant | significant | insignificant | insignificant | insignificant | 1 | 8 | 1 |
| denovoLocus57433 | insignificant | insignificant | insignificant | not genotyped | insignificant | insignificant | insignificant | insignificant | significant | insignificant | 1 | 8 | 1 |
| denovoLocus6384 | insignificant | insignificant | insignificant | insignificant | insignificant | not genotyped | significant | insignificant | insignificant | insignificant | 1 | 8 | 1 |
| denovoLocus6614 | insignificant | insignificant | insignificant | significant | insignificant | insignificant | insignificant | not genotyped | insignificant | insignificant | 1 | 8 | 1 |
| denovoLocus730 | significant | insignificant | insignificant | insignificant | insignificant | insignificant | not genotyped | insignificant | insignificant | insignificant | 1 | 8 | 1 |
| denovoLocus73994 | not genotyped | insignificant | insignificant | insignificant | insignificant | insignificant | significant | insignificant | insignificant | insignificant | 1 | 8 | 1 |
| denovoLocus76105 | insignificant | insignificant | not genotyped | insignificant | insignificant | insignificant | significant | insignificant | insignificant | insignificant | 1 | 8 | 1 |
| denovoLocus7905 | insignificant | insignificant | insignificant | insignificant | insignificant | insignificant | not genotyped | insignificant | insignificant | significant | 1 | 8 | 1 |
| denovoLocus8141 | insignificant | insignificant | significant | insignificant | insignificant | not genotyped | insignificant | insignificant | insignificant | insignificant | 1 | 8 | 1 |
| denovoLocus8617 | insignificant | insignificant | significant | insignificant | not genotyped | insignificant | insignificant | insignificant | insignificant | insignificant | 1 | 8 | 1 |
| denovoLocus95 | insignificant | significant | insignificant | insignificant | insignificant | not genotyped | insignificant | insignificant | insignificant | insignificant | 1 | 8 | 1 |
| denovoLocus10177 | not genotyped | insignificant | significant | insignificant | insignificant | insignificant | insignificant | insignificant | not genotyped | insignificant | 1 | 7 | 2 |
| denovoLocus10477 | insignificant | not genotyped | insignificant | significant | insignificant | insignificant | insignificant | insignificant | not genotyped | insignificant | 1 | 7 | 2 |
| denovoLocus105867 | not genotyped | insignificant | insignificant | not genotyped | insignificant | insignificant | significant | insignificant | insignificant | insignificant | 1 | 7 | 2 |
| denovoLocus11170 | insignificant | not genotyped | insignificant | insignificant | insignificant | not genotyped | significant | insignificant | insignificant | insignificant | 1 | 7 | 2 |
| denovoLocus11998 | insignificant | insignificant | insignificant | insignificant | not genotyped | insignificant | significant | insignificant | insignificant | not genotyped | 1 | 7 | 2 |
| denovoLocus12624 | insignificant | insignificant | insignificant | not genotyped | significant | insignificant | not genotyped | insignificant | insignificant | insignificant | 1 | 7 | 2 |
| denovoLocus12732 | insignificant | insignificant | insignificant | insignificant | insignificant | not genotyped | significant | insignificant | not genotyped | insignificant | 1 | 7 | 2 |
| denovoLocus14368 | insignificant | insignificant | significant | not genotyped | insignificant | insignificant | insignificant | insignificant | not genotyped | insignificant | 1 | 7 | 2 |
| denovoLocus14408 | insignificant | insignificant | insignificant | insignificant | insignificant | insignificant | significant | not genotyped | not genotyped | insignificant | 1 | 7 | 2 |
| denovoLocus145 | insignificant | insignificant | insignificant | not genotyped | insignificant | significant | not genotyped | insignificant | insignificant | insignificant | 1 | 7 | 2 |
| denovoLocus14524 | insignificant | insignificant | insignificant | insignificant | insignificant | insignificant | insignificant | not genotyped | significant | not genotyped | 1 | 7 | 2 |
| denovoLocus14589 | insignificant | insignificant | not genotyped | insignificant | insignificant | not genotyped | insignificant | insignificant | insignificant | significant | 1 | 7 | 2 |
| denovoLocus14726 | significant | insignificant | insignificant | not genotyped | insignificant | insignificant | insignificant | insignificant | not genotyped | insignificant | 1 | 7 | 2 |
| denovoLocus15324 | insignificant | insignificant | significant | insignificant | not genotyped | insignificant | not genotyped | insignificant | insignificant | insignificant | 1 | 7 | 2 |
| denovoLocus15442 | not genotyped | insignificant | insignificant | insignificant | significant | insignificant | insignificant | not genotyped | insignificant | insignificant | 1 | 7 | 2 |
| denovoLocus16544 | not genotyped | insignificant | insignificant | insignificant | insignificant | insignificant | significant | not genotyped | insignificant | insignificant | 1 | 7 | 2 |
| denovoLocus17118 | insignificant | insignificant | insignificant | insignificant | insignificant | insignificant | insignificant | significant | not genotyped | not genotyped | 1 | 7 | 2 |
| denovoLocus17371 | insignificant | not genotyped | insignificant | not genotyped | insignificant | significant | insignificant | insignificant | insignificant | insignificant | 1 | 7 | 2 |
| denovoLocus17703 | insignificant | insignificant | insignificant | insignificant | not genotyped | insignificant | insignificant | not genotyped | significant | insignificant | 1 | 7 | 2 |
| denovoLocus18034 | significant | insignificant | insignificant | not genotyped | insignificant | insignificant | insignificant | insignificant | not genotyped | insignificant | 1 | 7 | 2 |
| denovoLocus1903 | insignificant | not genotyped | not genotyped | insignificant | significant | insignificant | insignificant | insignificant | insignificant | insignificant | 1 | 7 | 2 |
| denovoLocus19763 | insignificant | insignificant | insignificant | not genotyped | not genotyped | insignificant | significant | insignificant | insignificant | insignificant | 1 | 7 | 2 |
| denovoLocus20524 | significant | not genotyped | insignificant | insignificant | insignificant | insignificant | not genotyped | insignificant | insignificant | insignificant | 1 | 7 | 2 |
| denovoLocus2145 | insignificant | insignificant | insignificant | insignificant | significant | insignificant | not genotyped | not genotyped | insignificant | insignificant | 1 | 7 | 2 |
| denovoLocus21873 | not genotyped | insignificant | insignificant | insignificant | insignificant | insignificant | not genotyped | insignificant | significant | insignificant | 1 | 7 | 2 |
| denovoLocus24859 | insignificant | not genotyped | insignificant | insignificant | insignificant | insignificant | significant | not genotyped | insignificant | insignificant | 1 | 7 | 2 |
| denovoLocus26462 | insignificant | insignificant | not genotyped | insignificant | insignificant | insignificant | significant | insignificant | not genotyped | insignificant | 1 | 7 | 2 |
| denovoLocus27022 | insignificant | insignificant | insignificant | not genotyped | insignificant | insignificant | insignificant | insignificant | not genotyped | significant | 1 | 7 | 2 |
| denovoLocus27973 | not genotyped | insignificant | insignificant | insignificant | insignificant | insignificant | significant | insignificant | insignificant | not genotyped | 1 | 7 | 2 |
| denovoLocus2812 | insignificant | insignificant | insignificant | insignificant | not genotyped | not genotyped | significant | insignificant | insignificant | insignificant | 1 | 7 | 2 |
| denovoLocus28945 | insignificant | insignificant | insignificant | not genotyped | insignificant | insignificant | significant | insignificant | not genotyped | insignificant | 1 | 7 | 2 |
| denovoLocus30345 | not genotyped | insignificant | insignificant | insignificant | significant | insignificant | insignificant | not genotyped | insignificant | insignificant | 1 | 7 | 2 |
| denovoLocus308 | insignificant | insignificant | insignificant | insignificant | insignificant | not genotyped | not genotyped | insignificant | significant | insignificant | 1 | 7 | 2 |
| denovoLocus31362 | significant | insignificant | insignificant | insignificant | not genotyped | insignificant | insignificant | insignificant | not genotyped | insignificant | 1 | 7 | 2 |
| denovoLocus31374 | not genotyped | insignificant | not genotyped | insignificant | insignificant | insignificant | insignificant | insignificant | insignificant | significant | 1 | 7 | 2 |
| denovoLocus31514 | not genotyped | insignificant | not genotyped | insignificant | significant | insignificant | insignificant | insignificant | insignificant | insignificant | 1 | 7 | 2 |
| denovoLocus32015 | insignificant | insignificant | not genotyped | insignificant | insignificant | insignificant | significant | insignificant | insignificant | not genotyped | 1 | 7 | 2 |
| denovoLocus3262 | insignificant | insignificant | insignificant | insignificant | not genotyped | insignificant | significant | insignificant | insignificant | not genotyped | 1 | 7 | 2 |
| denovoLocus33732 | insignificant | insignificant | insignificant | insignificant | not genotyped | insignificant | significant | insignificant | not genotyped | insignificant | 1 | 7 | 2 |
| denovoLocus35591 | insignificant | insignificant | insignificant | insignificant | insignificant | insignificant | significant | not genotyped | not genotyped | insignificant | 1 | 7 | 2 |
| denovoLocus35681 | significant | insignificant | insignificant | not genotyped | insignificant | not genotyped | insignificant | insignificant | insignificant | insignificant | 1 | 7 | 2 |
| denovoLocus3670 | not genotyped | insignificant | insignificant | not genotyped | insignificant | insignificant | significant | insignificant | insignificant | insignificant | 1 | 7 | 2 |
| denovoLocus37152 | insignificant | insignificant | insignificant | insignificant | insignificant | insignificant | significant | not genotyped | not genotyped | insignificant | 1 | 7 | 2 |
| denovoLocus38134 | not genotyped | insignificant | insignificant | insignificant | insignificant | insignificant | insignificant | significant | insignificant | not genotyped | 1 | 7 | 2 |
| denovoLocus41280 | insignificant | insignificant | insignificant | insignificant | insignificant | insignificant | insignificant | not genotyped | not genotyped | significant | 1 | 7 | 2 |
| denovoLocus44459 | not genotyped | insignificant | insignificant | insignificant | not genotyped | insignificant | insignificant | insignificant | insignificant | significant | 1 | 7 | 2 |
| denovoLocus4518 | insignificant | insignificant | not genotyped | insignificant | insignificant | insignificant | insignificant | not genotyped | insignificant | significant | 1 | 7 | 2 |
| denovoLocus4717 | not genotyped | insignificant | insignificant | insignificant | insignificant | insignificant | insignificant | significant | not genotyped | insignificant | 1 | 7 | 2 |
| denovoLocus48195 | insignificant | insignificant | not genotyped | insignificant | insignificant | insignificant | significant | insignificant | insignificant | not genotyped | 1 | 7 | 2 |
| denovoLocus483 | significant | insignificant | insignificant | not genotyped | not genotyped | insignificant | insignificant | insignificant | insignificant | insignificant | 1 | 7 | 2 |
| denovoLocus49311 | insignificant | insignificant | insignificant | insignificant | insignificant | not genotyped | insignificant | insignificant | not genotyped | significant | 1 | 7 | 2 |
| denovoLocus4943 | insignificant | insignificant | not genotyped | insignificant | insignificant | insignificant | insignificant | not genotyped | insignificant | significant | 1 | 7 | 2 |
| denovoLocus49826 | insignificant | insignificant | not genotyped | insignificant | insignificant | insignificant | significant | not genotyped | insignificant | insignificant | 1 | 7 | 2 |
| denovoLocus5185 | insignificant | insignificant | significant | not genotyped | insignificant | insignificant | insignificant | insignificant | not genotyped | insignificant | 1 | 7 | 2 |
| denovoLocus53102 | insignificant | not genotyped | insignificant | insignificant | insignificant | insignificant | insignificant | not genotyped | insignificant | significant | 1 | 7 | 2 |
| denovoLocus53233 | insignificant | insignificant | insignificant | insignificant | insignificant | not genotyped | significant | insignificant | not genotyped | insignificant | 1 | 7 | 2 |
| denovoLocus53297 | insignificant | insignificant | insignificant | insignificant | insignificant | not genotyped | insignificant | significant | not genotyped | insignificant | 1 | 7 | 2 |
| denovoLocus55567 | insignificant | not genotyped | significant | insignificant | insignificant | insignificant | insignificant | not genotyped | insignificant | insignificant | 1 | 7 | 2 |
| denovoLocus58589 | significant | insignificant | not genotyped | not genotyped | insignificant | insignificant | insignificant | insignificant | insignificant | insignificant | 1 | 7 | 2 |
| denovoLocus59334 | insignificant | insignificant | not genotyped | insignificant | insignificant | insignificant | significant | not genotyped | insignificant | insignificant | 1 | 7 | 2 |
| denovoLocus60169 | insignificant | insignificant | not genotyped | not genotyped | insignificant | insignificant | insignificant | insignificant | insignificant | significant | 1 | 7 | 2 |
| denovoLocus60665 | not genotyped | not genotyped | insignificant | insignificant | insignificant | insignificant | significant | insignificant | insignificant | insignificant | 1 | 7 | 2 |
| denovoLocus6175 | insignificant | insignificant | significant | insignificant | not genotyped | insignificant | not genotyped | insignificant | insignificant | insignificant | 1 | 7 | 2 |
| denovoLocus62805 | insignificant | insignificant | insignificant | insignificant | insignificant | not genotyped | significant | insignificant | insignificant | not genotyped | 1 | 7 | 2 |
| denovoLocus63777 | insignificant | not genotyped | insignificant | not genotyped | insignificant | insignificant | significant | insignificant | insignificant | insignificant | 1 | 7 | 2 |
| denovoLocus6642 | not genotyped | insignificant | significant | insignificant | not genotyped | insignificant | insignificant | insignificant | insignificant | insignificant | 1 | 7 | 2 |
| denovoLocus6706 | insignificant | insignificant | insignificant | insignificant | insignificant | not genotyped | significant | insignificant | insignificant | not genotyped | 1 | 7 | 2 |
| denovoLocus692 | insignificant | insignificant | not genotyped | insignificant | not genotyped | insignificant | insignificant | significant | insignificant | insignificant | 1 | 7 | 2 |
| denovoLocus70390 | insignificant | insignificant | not genotyped | insignificant | insignificant | insignificant | insignificant | not genotyped | significant | insignificant | 1 | 7 | 2 |
| denovoLocus72257 | insignificant | insignificant | not genotyped | insignificant | insignificant | not genotyped | significant | insignificant | insignificant | insignificant | 1 | 7 | 2 |
| denovoLocus7951 | insignificant | insignificant | insignificant | insignificant | not genotyped | insignificant | insignificant | insignificant | not genotyped | significant | 1 | 7 | 2 |
| denovoLocus81958 | insignificant | insignificant | insignificant | insignificant | insignificant | not genotyped | not genotyped | insignificant | significant | insignificant | 1 | 7 | 2 |
| denovoLocus826 | insignificant | not genotyped | insignificant | not genotyped | insignificant | insignificant | insignificant | insignificant | insignificant | significant | 1 | 7 | 2 |
| denovoLocus8385 | not genotyped | insignificant | insignificant | insignificant | insignificant | not genotyped | insignificant | significant | insignificant | insignificant | 1 | 7 | 2 |
| denovoLocus8678 | insignificant | insignificant | not genotyped | insignificant | insignificant | insignificant | significant | not genotyped | insignificant | insignificant | 1 | 7 | 2 |
| denovoLocus9438 | insignificant | insignificant | insignificant | insignificant | insignificant | insignificant | significant | insignificant | not genotyped | not genotyped | 1 | 7 | 2 |
| denovoLocus9441 | insignificant | insignificant | significant | insignificant | insignificant | insignificant | not genotyped | insignificant | insignificant | not genotyped | 1 | 7 | 2 |
| denovoLocus9587 | insignificant | insignificant | insignificant | insignificant | insignificant | not genotyped | insignificant | insignificant | not genotyped | significant | 1 | 7 | 2 |
| denovoLocus9661 | insignificant | insignificant | not genotyped | insignificant | significant | insignificant | insignificant | insignificant | not genotyped | insignificant | 1 | 7 | 2 |
| denovoLocus10061 | insignificant | insignificant | not genotyped | insignificant | insignificant | insignificant | not genotyped | insignificant | significant | not genotyped | 1 | 6 | 3 |
| denovoLocus10391 | not genotyped | not genotyped | significant | not genotyped | insignificant | insignificant | insignificant | insignificant | insignificant | insignificant | 1 | 6 | 3 |
| denovoLocus13025 | insignificant | not genotyped | insignificant | not genotyped | insignificant | insignificant | significant | not genotyped | insignificant | insignificant | 1 | 6 | 3 |
| denovoLocus13645 | significant | not genotyped | insignificant | insignificant | not genotyped | insignificant | not genotyped | insignificant | insignificant | insignificant | 1 | 6 | 3 |
| denovoLocus13923 | not genotyped | insignificant | not genotyped | insignificant | insignificant | insignificant | significant | not genotyped | insignificant | insignificant | 1 | 6 | 3 |
| denovoLocus1441 | significant | not genotyped | not genotyped | not genotyped | insignificant | insignificant | insignificant | insignificant | insignificant | insignificant | 1 | 6 | 3 |
| denovoLocus157 | insignificant | not genotyped | insignificant | insignificant | not genotyped | insignificant | significant | insignificant | insignificant | not genotyped | 1 | 6 | 3 |
| denovoLocus15747 | insignificant | not genotyped | not genotyped | insignificant | insignificant | not genotyped | significant | insignificant | insignificant | insignificant | 1 | 6 | 3 |
| denovoLocus1625 | insignificant | not genotyped | insignificant | insignificant | insignificant | significant | insignificant | insignificant | not genotyped | not genotyped | 1 | 6 | 3 |
| denovoLocus1633 | insignificant | not genotyped | insignificant | not genotyped | not genotyped | insignificant | significant | insignificant | insignificant | insignificant | 1 | 6 | 3 |
| denovoLocus16706 | insignificant | not genotyped | insignificant | insignificant | significant | not genotyped | not genotyped | insignificant | insignificant | insignificant | 1 | 6 | 3 |
| denovoLocus16988 | insignificant | not genotyped | not genotyped | not genotyped | insignificant | insignificant | significant | insignificant | insignificant | insignificant | 1 | 6 | 3 |
| denovoLocus17068 | not genotyped | insignificant | significant | insignificant | insignificant | not genotyped | insignificant | insignificant | not genotyped | insignificant | 1 | 6 | 3 |
| denovoLocus17217 | insignificant | not genotyped | not genotyped | insignificant | insignificant | insignificant | insignificant | insignificant | not genotyped | significant | 1 | 6 | 3 |
| denovoLocus17285 | insignificant | insignificant | insignificant | not genotyped | insignificant | insignificant | insignificant | not genotyped | not genotyped | significant | 1 | 6 | 3 |
| denovoLocus17347 | insignificant | insignificant | not genotyped | not genotyped | not genotyped | insignificant | insignificant | insignificant | significant | insignificant | 1 | 6 | 3 |
| denovoLocus20722 | significant | insignificant | insignificant | insignificant | not genotyped | not genotyped | not genotyped | insignificant | insignificant | insignificant | 1 | 6 | 3 |
| denovoLocus21774 | insignificant | insignificant | significant | insignificant | insignificant | not genotyped | insignificant | not genotyped | insignificant | not genotyped | 1 | 6 | 3 |
| denovoLocus2205 | not genotyped | insignificant | significant | not genotyped | insignificant | insignificant | not genotyped | insignificant | insignificant | insignificant | 1 | 6 | 3 |
| denovoLocus22105 | not genotyped | insignificant | not genotyped | insignificant | insignificant | not genotyped | significant | insignificant | insignificant | insignificant | 1 | 6 | 3 |
| denovoLocus22729 | insignificant | not genotyped | significant | not genotyped | insignificant | insignificant | insignificant | not genotyped | insignificant | insignificant | 1 | 6 | 3 |
| denovoLocus23088 | insignificant | not genotyped | significant | not genotyped | insignificant | insignificant | insignificant | not genotyped | insignificant | insignificant | 1 | 6 | 3 |
| denovoLocus23375 | insignificant | insignificant | insignificant | not genotyped | insignificant | insignificant | significant | not genotyped | insignificant | not genotyped | 1 | 6 | 3 |
| denovoLocus24828 | insignificant | insignificant | significant | insignificant | insignificant | not genotyped | not genotyped | insignificant | not genotyped | insignificant | 1 | 6 | 3 |
| denovoLocus25136 | insignificant | insignificant | not genotyped | insignificant | not genotyped | insignificant | significant | insignificant | not genotyped | insignificant | 1 | 6 | 3 |
| denovoLocus25349 | significant | insignificant | not genotyped | insignificant | insignificant | not genotyped | not genotyped | insignificant | insignificant | insignificant | 1 | 6 | 3 |
| denovoLocus2565 | insignificant | not genotyped | not genotyped | insignificant | insignificant | insignificant | significant | insignificant | insignificant | not genotyped | 1 | 6 | 3 |
| denovoLocus25762 | insignificant | insignificant | not genotyped | insignificant | insignificant | insignificant | insignificant | not genotyped | not genotyped | significant | 1 | 6 | 3 |
| denovoLocus25799 | insignificant | not genotyped | insignificant | insignificant | not genotyped | insignificant | not genotyped | significant | insignificant | insignificant | 1 | 6 | 3 |
| denovoLocus25924 | insignificant | insignificant | not genotyped | insignificant | not genotyped | insignificant | insignificant | not genotyped | insignificant | significant | 1 | 6 | 3 |
| denovoLocus26140 | not genotyped | insignificant | not genotyped | not genotyped | insignificant | insignificant | significant | insignificant | insignificant | insignificant | 1 | 6 | 3 |
| denovoLocus2689 | not genotyped | insignificant | not genotyped | insignificant | insignificant | significant | insignificant | insignificant | not genotyped | insignificant | 1 | 6 | 3 |
| denovoLocus27102 | not genotyped | insignificant | insignificant | not genotyped | insignificant | insignificant | significant | insignificant | not genotyped | insignificant | 1 | 6 | 3 |
| denovoLocus27139 | insignificant | insignificant | significant | insignificant | not genotyped | insignificant | insignificant | insignificant | not genotyped | not genotyped | 1 | 6 | 3 |
| denovoLocus28341 | insignificant | insignificant | not genotyped | not genotyped | not genotyped | insignificant | significant | insignificant | insignificant | insignificant | 1 | 6 | 3 |
| denovoLocus2836 | not genotyped | insignificant | not genotyped | insignificant | insignificant | insignificant | insignificant | not genotyped | insignificant | significant | 1 | 6 | 3 |
| denovoLocus28899 | not genotyped | insignificant | insignificant | not genotyped | insignificant | insignificant | significant | insignificant | not genotyped | insignificant | 1 | 6 | 3 |
| denovoLocus28996 | insignificant | insignificant | not genotyped | insignificant | insignificant | insignificant | significant | not genotyped | not genotyped | insignificant | 1 | 6 | 3 |
| denovoLocus29382 | not genotyped | insignificant | insignificant | insignificant | insignificant | insignificant | significant | not genotyped | not genotyped | insignificant | 1 | 6 | 3 |
| denovoLocus29606 | insignificant | not genotyped | not genotyped | insignificant | insignificant | insignificant | significant | insignificant | not genotyped | insignificant | 1 | 6 | 3 |
| denovoLocus32148 | insignificant | not genotyped | significant | insignificant | insignificant | insignificant | insignificant | not genotyped | insignificant | not genotyped | 1 | 6 | 3 |
| denovoLocus35012 | insignificant | insignificant | insignificant | insignificant | insignificant | not genotyped | insignificant | not genotyped | significant | not genotyped | 1 | 6 | 3 |
| denovoLocus35329 | significant | insignificant | insignificant | not genotyped | insignificant | insignificant | insignificant | not genotyped | insignificant | not genotyped | 1 | 6 | 3 |
| denovoLocus36728 | insignificant | insignificant | insignificant | insignificant | not genotyped | not genotyped | significant | insignificant | insignificant | not genotyped | 1 | 6 | 3 |
| denovoLocus37557 | not genotyped | not genotyped | not genotyped | insignificant | insignificant | insignificant | significant | insignificant | insignificant | insignificant | 1 | 6 | 3 |
| denovoLocus37626 | insignificant | insignificant | insignificant | insignificant | not genotyped | insignificant | not genotyped | not genotyped | insignificant | significant | 1 | 6 | 3 |
| denovoLocus37948 | insignificant | insignificant | insignificant | insignificant | insignificant | insignificant | not genotyped | not genotyped | not genotyped | significant | 1 | 6 | 3 |
| denovoLocus40075 | insignificant | not genotyped | insignificant | insignificant | not genotyped | insignificant | not genotyped | significant | insignificant | insignificant | 1 | 6 | 3 |
| denovoLocus41519 | insignificant | insignificant | not genotyped | not genotyped | insignificant | insignificant | significant | insignificant | not genotyped | insignificant | 1 | 6 | 3 |
| denovoLocus41611 | insignificant | not genotyped | insignificant | not genotyped | insignificant | not genotyped | insignificant | insignificant | significant | insignificant | 1 | 6 | 3 |
| denovoLocus45659 | not genotyped | insignificant | insignificant | insignificant | insignificant | insignificant | not genotyped | not genotyped | insignificant | significant | 1 | 6 | 3 |
| denovoLocus48009 | insignificant | insignificant | not genotyped | insignificant | insignificant | not genotyped | significant | insignificant | not genotyped | insignificant | 1 | 6 | 3 |
| denovoLocus4921 | not genotyped | not genotyped | not genotyped | insignificant | insignificant | insignificant | significant | insignificant | insignificant | insignificant | 1 | 6 | 3 |
| denovoLocus51314 | insignificant | significant | insignificant | insignificant | not genotyped | not genotyped | insignificant | insignificant | insignificant | not genotyped | 1 | 6 | 3 |
| denovoLocus52108 | insignificant | insignificant | not genotyped | significant | insignificant | insignificant | insignificant | insignificant | not genotyped | not genotyped | 1 | 6 | 3 |
| denovoLocus53049 | not genotyped | insignificant | significant | insignificant | insignificant | not genotyped | insignificant | not genotyped | insignificant | insignificant | 1 | 6 | 3 |
| denovoLocus54426 | insignificant | insignificant | insignificant | not genotyped | not genotyped | not genotyped | significant | insignificant | insignificant | insignificant | 1 | 6 | 3 |
| denovoLocus55171 | not genotyped | insignificant | insignificant | insignificant | not genotyped | not genotyped | insignificant | insignificant | insignificant | significant | 1 | 6 | 3 |
| denovoLocus56939 | insignificant | not genotyped | not genotyped | significant | insignificant | insignificant | insignificant | insignificant | not genotyped | insignificant | 1 | 6 | 3 |
| denovoLocus57207 | insignificant | insignificant | not genotyped | insignificant | insignificant | not genotyped | insignificant | insignificant | significant | not genotyped | 1 | 6 | 3 |
| denovoLocus57329 | insignificant | insignificant | not genotyped | not genotyped | insignificant | significant | insignificant | not genotyped | insignificant | insignificant | 1 | 6 | 3 |
| denovoLocus58071 | not genotyped | insignificant | insignificant | insignificant | insignificant | insignificant | insignificant | not genotyped | not genotyped | significant | 1 | 6 | 3 |
| denovoLocus58206 | insignificant | not genotyped | insignificant | not genotyped | insignificant | insignificant | not genotyped | significant | insignificant | insignificant | 1 | 6 | 3 |
| denovoLocus5836 | not genotyped | insignificant | insignificant | not genotyped | insignificant | not genotyped | insignificant | insignificant | significant | insignificant | 1 | 6 | 3 |
| denovoLocus6039 | insignificant | not genotyped | insignificant | insignificant | not genotyped | insignificant | significant | not genotyped | insignificant | insignificant | 1 | 6 | 3 |
| denovoLocus61132 | not genotyped | insignificant | insignificant | insignificant | not genotyped | insignificant | significant | insignificant | insignificant | not genotyped | 1 | 6 | 3 |
| denovoLocus62783 | insignificant | not genotyped | insignificant | not genotyped | insignificant | insignificant | significant | not genotyped | insignificant | insignificant | 1 | 6 | 3 |
| denovoLocus67110 | not genotyped | insignificant | insignificant | insignificant | not genotyped | insignificant | not genotyped | insignificant | significant | insignificant | 1 | 6 | 3 |
| denovoLocus67419 | insignificant | insignificant | not genotyped | insignificant | insignificant | insignificant | significant | not genotyped | not genotyped | insignificant | 1 | 6 | 3 |
| denovoLocus73915 | not genotyped | insignificant | insignificant | insignificant | insignificant | insignificant | not genotyped | insignificant | significant | not genotyped | 1 | 6 | 3 |
| denovoLocus7769 | not genotyped | insignificant | insignificant | insignificant | significant | insignificant | not genotyped | not genotyped | insignificant | insignificant | 1 | 6 | 3 |
| denovoLocus77718 | insignificant | not genotyped | insignificant | insignificant | insignificant | insignificant | significant | not genotyped | insignificant | not genotyped | 1 | 6 | 3 |
| denovoLocus79795 | not genotyped | insignificant | insignificant | not genotyped | significant | insignificant | insignificant | insignificant | insignificant | not genotyped | 1 | 6 | 3 |
| denovoLocus79806 | insignificant | insignificant | significant | not genotyped | insignificant | insignificant | not genotyped | insignificant | insignificant | not genotyped | 1 | 6 | 3 |
| denovoLocus8065 | insignificant | insignificant | insignificant | not genotyped | not genotyped | insignificant | significant | not genotyped | insignificant | insignificant | 1 | 6 | 3 |
| denovoLocus8455 | not genotyped | insignificant | not genotyped | insignificant | insignificant | insignificant | not genotyped | insignificant | significant | insignificant | 1 | 6 | 3 |
| denovoLocus84611 | insignificant | not genotyped | not genotyped | significant | insignificant | insignificant | insignificant | insignificant | insignificant | not genotyped | 1 | 6 | 3 |
| denovoLocus8516 | insignificant | not genotyped | insignificant | insignificant | insignificant | insignificant | significant | insignificant | not genotyped | not genotyped | 1 | 6 | 3 |
| denovoLocus8767 | not genotyped | insignificant | not genotyped | significant | insignificant | insignificant | not genotyped | insignificant | insignificant | insignificant | 1 | 6 | 3 |
| denovoLocus93225 | insignificant | not genotyped | insignificant | insignificant | not genotyped | significant | insignificant | insignificant | not genotyped | insignificant | 1 | 6 | 3 |
| denovoLocus94112 | not genotyped | insignificant | insignificant | insignificant | insignificant | not genotyped | not genotyped | significant | insignificant | insignificant | 1 | 6 | 3 |
| denovoLocus94242 | insignificant | insignificant | insignificant | not genotyped | insignificant | significant | not genotyped | not genotyped | insignificant | insignificant | 1 | 6 | 3 |
| denovoLocus94269 | insignificant | insignificant | not genotyped | insignificant | not genotyped | not genotyped | significant | insignificant | insignificant | insignificant | 1 | 6 | 3 |
| denovoLocus9639 | insignificant | not genotyped | insignificant | not genotyped | insignificant | insignificant | insignificant | insignificant | significant | not genotyped | 1 | 6 | 3 |
| denovoLocus101807 | not genotyped | insignificant | not genotyped | insignificant | not genotyped | insignificant | significant | not genotyped | insignificant | insignificant | 1 | 5 | 4 |
| denovoLocus10491 | significant | not genotyped | not genotyped | not genotyped | insignificant | insignificant | not genotyped | insignificant | insignificant | insignificant | 1 | 5 | 4 |
| denovoLocus10759 | insignificant | insignificant | not genotyped | not genotyped | significant | not genotyped | not genotyped | insignificant | insignificant | insignificant | 1 | 5 | 4 |
| denovoLocus10984 | significant | insignificant | not genotyped | insignificant | insignificant | not genotyped | insignificant | not genotyped | insignificant | not genotyped | 1 | 5 | 4 |
| denovoLocus11363 | insignificant | insignificant | significant | insignificant | not genotyped | not genotyped | not genotyped | insignificant | not genotyped | insignificant | 1 | 5 | 4 |
| denovoLocus11455 | insignificant | not genotyped | not genotyped | not genotyped | insignificant | insignificant | significant | insignificant | not genotyped | insignificant | 1 | 5 | 4 |
| denovoLocus114643 | insignificant | not genotyped | insignificant | insignificant | not genotyped | significant | insignificant | not genotyped | insignificant | not genotyped | 1 | 5 | 4 |
| denovoLocus11465 | not genotyped | insignificant | not genotyped | not genotyped | insignificant | not genotyped | insignificant | insignificant | significant | insignificant | 1 | 5 | 4 |
| denovoLocus119596 | not genotyped | insignificant | not genotyped | not genotyped | insignificant | not genotyped | significant | insignificant | insignificant | insignificant | 1 | 5 | 4 |
| denovoLocus13171 | significant | not genotyped | not genotyped | insignificant | not genotyped | insignificant | insignificant | insignificant | not genotyped | insignificant | 1 | 5 | 4 |
| denovoLocus13874 | not genotyped | insignificant | not genotyped | insignificant | insignificant | insignificant | not genotyped | not genotyped | significant | insignificant | 1 | 5 | 4 |
| denovoLocus13992 | insignificant | not genotyped | insignificant | insignificant | insignificant | insignificant | significant | not genotyped | not genotyped | not genotyped | 1 | 5 | 4 |
| denovoLocus14155 | insignificant | not genotyped | insignificant | insignificant | not genotyped | insignificant | significant | insignificant | not genotyped | not genotyped | 1 | 5 | 4 |
| denovoLocus1442 | not genotyped | insignificant | not genotyped | not genotyped | insignificant | insignificant | significant | not genotyped | insignificant | insignificant | 1 | 5 | 4 |
| denovoLocus14839 | not genotyped | insignificant | insignificant | not genotyped | not genotyped | not genotyped | significant | insignificant | insignificant | insignificant | 1 | 5 | 4 |
| denovoLocus15156 | significant | not genotyped | not genotyped | insignificant | insignificant | insignificant | insignificant | insignificant | not genotyped | not genotyped | 1 | 5 | 4 |
| denovoLocus1528 | not genotyped | insignificant | significant | insignificant | insignificant | not genotyped | not genotyped | insignificant | insignificant | not genotyped | 1 | 5 | 4 |
| denovoLocus15651 | not genotyped | insignificant | significant | not genotyped | insignificant | insignificant | insignificant | insignificant | not genotyped | not genotyped | 1 | 5 | 4 |
| denovoLocus16008 | insignificant | not genotyped | insignificant | insignificant | insignificant | insignificant | not genotyped | not genotyped | not genotyped | significant | 1 | 5 | 4 |
| denovoLocus16905 | not genotyped | insignificant | not genotyped | insignificant | not genotyped | insignificant | not genotyped | insignificant | insignificant | significant | 1 | 5 | 4 |
| denovoLocus1950 | insignificant | insignificant | not genotyped | insignificant | not genotyped | insignificant | not genotyped | not genotyped | significant | insignificant | 1 | 5 | 4 |
| denovoLocus19541 | insignificant | not genotyped | not genotyped | not genotyped | insignificant | insignificant | significant | insignificant | insignificant | not genotyped | 1 | 5 | 4 |
| denovoLocus19643 | insignificant | not genotyped | insignificant | significant | insignificant | insignificant | not genotyped | not genotyped | not genotyped | insignificant | 1 | 5 | 4 |
| denovoLocus19743 | not genotyped | insignificant | not genotyped | insignificant | not genotyped | not genotyped | significant | insignificant | insignificant | insignificant | 1 | 5 | 4 |
| denovoLocus20310 | not genotyped | not genotyped | insignificant | insignificant | not genotyped | insignificant | not genotyped | insignificant | insignificant | significant | 1 | 5 | 4 |
| denovoLocus20415 | insignificant | insignificant | insignificant | not genotyped | not genotyped | not genotyped | significant | not genotyped | insignificant | insignificant | 1 | 5 | 4 |
| denovoLocus22672 | insignificant | not genotyped | insignificant | insignificant | insignificant | not genotyped | significant | insignificant | not genotyped | not genotyped | 1 | 5 | 4 |
| denovoLocus22746 | insignificant | insignificant | not genotyped | insignificant | insignificant | insignificant | significant | not genotyped | not genotyped | not genotyped | 1 | 5 | 4 |
| denovoLocus22865 | not genotyped | insignificant | not genotyped | not genotyped | insignificant | insignificant | significant | not genotyped | insignificant | insignificant | 1 | 5 | 4 |
| denovoLocus24177 | not genotyped | insignificant | insignificant | insignificant | insignificant | not genotyped | not genotyped | not genotyped | significant | insignificant | 1 | 5 | 4 |
| denovoLocus24265 | insignificant | not genotyped | insignificant | not genotyped | insignificant | not genotyped | significant | not genotyped | insignificant | insignificant | 1 | 5 | 4 |
| denovoLocus24611 | not genotyped | not genotyped | insignificant | insignificant | significant | not genotyped | insignificant | not genotyped | insignificant | insignificant | 1 | 5 | 4 |
| denovoLocus25987 | insignificant | insignificant | not genotyped | not genotyped | not genotyped | insignificant | significant | insignificant | not genotyped | insignificant | 1 | 5 | 4 |
| denovoLocus26826 | not genotyped | insignificant | insignificant | not genotyped | not genotyped | not genotyped | significant | insignificant | insignificant | insignificant | 1 | 5 | 4 |
| denovoLocus27070 | insignificant | insignificant | insignificant | not genotyped | not genotyped | not genotyped | insignificant | not genotyped | significant | insignificant | 1 | 5 | 4 |
| denovoLocus28821 | insignificant | not genotyped | insignificant | insignificant | not genotyped | insignificant | significant | insignificant | not genotyped | not genotyped | 1 | 5 | 4 |
| denovoLocus29891 | not genotyped | not genotyped | significant | insignificant | not genotyped | insignificant | insignificant | not genotyped | insignificant | insignificant | 1 | 5 | 4 |
| denovoLocus30064 | insignificant | not genotyped | insignificant | insignificant | not genotyped | insignificant | significant | not genotyped | not genotyped | insignificant | 1 | 5 | 4 |
| denovoLocus30419 | insignificant | insignificant | insignificant | not genotyped | not genotyped | insignificant | significant | not genotyped | insignificant | not genotyped | 1 | 5 | 4 |
| denovoLocus3233 | not genotyped | insignificant | insignificant | not genotyped | not genotyped | not genotyped | insignificant | insignificant | insignificant | significant | 1 | 5 | 4 |
| denovoLocus3334 | not genotyped | insignificant | not genotyped | significant | insignificant | not genotyped | insignificant | not genotyped | insignificant | insignificant | 1 | 5 | 4 |
| denovoLocus3335 | insignificant | insignificant | insignificant | not genotyped | not genotyped | insignificant | significant | not genotyped | not genotyped | insignificant | 1 | 5 | 4 |
| denovoLocus34088 | insignificant | not genotyped | insignificant | insignificant | not genotyped | insignificant | insignificant | significant | not genotyped | not genotyped | 1 | 5 | 4 |
| denovoLocus343 | insignificant | not genotyped | insignificant | insignificant | not genotyped | insignificant | significant | not genotyped | not genotyped | insignificant | 1 | 5 | 4 |
| denovoLocus34448 | insignificant | insignificant | insignificant | significant | insignificant | not genotyped | not genotyped | insignificant | not genotyped | not genotyped | 1 | 5 | 4 |
| denovoLocus35373 | insignificant | insignificant | insignificant | not genotyped | not genotyped | insignificant | significant | insignificant | not genotyped | not genotyped | 1 | 5 | 4 |
| denovoLocus35448 | not genotyped | insignificant | insignificant | insignificant | significant | not genotyped | insignificant | not genotyped | not genotyped | insignificant | 1 | 5 | 4 |
| denovoLocus3617 | not genotyped | insignificant | not genotyped | insignificant | insignificant | insignificant | significant | not genotyped | insignificant | not genotyped | 1 | 5 | 4 |
| denovoLocus36613 | insignificant | insignificant | insignificant | not genotyped | not genotyped | not genotyped | significant | insignificant | insignificant | not genotyped | 1 | 5 | 4 |
| denovoLocus37135 | not genotyped | insignificant | not genotyped | insignificant | insignificant | not genotyped | insignificant | significant | insignificant | not genotyped | 1 | 5 | 4 |
| denovoLocus40232 | insignificant | not genotyped | insignificant | not genotyped | not genotyped | not genotyped | insignificant | insignificant | significant | insignificant | 1 | 5 | 4 |
| denovoLocus40933 | not genotyped | insignificant | insignificant | not genotyped | not genotyped | insignificant | insignificant | insignificant | significant | not genotyped | 1 | 5 | 4 |
| denovoLocus4111 | not genotyped | insignificant | not genotyped | not genotyped | not genotyped | insignificant | significant | insignificant | insignificant | insignificant | 1 | 5 | 4 |
| denovoLocus41621 | insignificant | not genotyped | not genotyped | insignificant | insignificant | insignificant | significant | not genotyped | not genotyped | insignificant | 1 | 5 | 4 |
| denovoLocus41719 | not genotyped | insignificant | not genotyped | not genotyped | insignificant | not genotyped | significant | insignificant | insignificant | insignificant | 1 | 5 | 4 |
| denovoLocus42250 | not genotyped | insignificant | insignificant | not genotyped | insignificant | insignificant | significant | not genotyped | not genotyped | insignificant | 1 | 5 | 4 |
| denovoLocus42542 | insignificant | insignificant | insignificant | not genotyped | not genotyped | not genotyped | significant | not genotyped | insignificant | insignificant | 1 | 5 | 4 |
| denovoLocus4255 | not genotyped | not genotyped | not genotyped | not genotyped | insignificant | insignificant | insignificant | insignificant | insignificant | significant | 1 | 5 | 4 |
| denovoLocus43559 | insignificant | insignificant | not genotyped | not genotyped | insignificant | not genotyped | significant | insignificant | insignificant | not genotyped | 1 | 5 | 4 |
| denovoLocus43732 | not genotyped | insignificant | significant | not genotyped | insignificant | not genotyped | not genotyped | insignificant | insignificant | insignificant | 1 | 5 | 4 |
| denovoLocus44011 | insignificant | not genotyped | insignificant | not genotyped | insignificant | insignificant | significant | not genotyped | not genotyped | insignificant | 1 | 5 | 4 |
| denovoLocus45458 | not genotyped | not genotyped | insignificant | insignificant | insignificant | insignificant | significant | not genotyped | not genotyped | insignificant | 1 | 5 | 4 |
| denovoLocus49905 | insignificant | insignificant | insignificant | significant | not genotyped | not genotyped | insignificant | insignificant | not genotyped | not genotyped | 1 | 5 | 4 |
| denovoLocus50769 | insignificant | insignificant | significant | not genotyped | not genotyped | not genotyped | not genotyped | insignificant | insignificant | insignificant | 1 | 5 | 4 |
| denovoLocus51099 | insignificant | insignificant | not genotyped | insignificant | insignificant | not genotyped | not genotyped | insignificant | significant | not genotyped | 1 | 5 | 4 |
| denovoLocus51295 | not genotyped | not genotyped | not genotyped | insignificant | insignificant | insignificant | not genotyped | significant | insignificant | insignificant | 1 | 5 | 4 |
| denovoLocus52444 | significant | not genotyped | insignificant | not genotyped | not genotyped | insignificant | insignificant | insignificant | not genotyped | insignificant | 1 | 5 | 4 |
| denovoLocus53062 | not genotyped | insignificant | significant | insignificant | insignificant | insignificant | not genotyped | insignificant | not genotyped | not genotyped | 1 | 5 | 4 |
| denovoLocus5314 | not genotyped | not genotyped | insignificant | insignificant | insignificant | insignificant | insignificant | significant | not genotyped | not genotyped | 1 | 5 | 4 |
| denovoLocus54354 | insignificant | insignificant | not genotyped | insignificant | insignificant | not genotyped | insignificant | not genotyped | not genotyped | significant | 1 | 5 | 4 |
| denovoLocus54752 | significant | insignificant | insignificant | not genotyped | not genotyped | insignificant | insignificant | not genotyped | not genotyped | insignificant | 1 | 5 | 4 |
| denovoLocus55250 | significant | not genotyped | not genotyped | not genotyped | not genotyped | insignificant | insignificant | insignificant | insignificant | insignificant | 1 | 5 | 4 |
| denovoLocus5530 | significant | insignificant | not genotyped | insignificant | not genotyped | not genotyped | insignificant | not genotyped | insignificant | insignificant | 1 | 5 | 4 |
| denovoLocus55417 | insignificant | insignificant | significant | not genotyped | insignificant | not genotyped | not genotyped | insignificant | insignificant | not genotyped | 1 | 5 | 4 |
| denovoLocus57286 | insignificant | insignificant | insignificant | not genotyped | insignificant | not genotyped | significant | insignificant | not genotyped | not genotyped | 1 | 5 | 4 |
| denovoLocus57727 | not genotyped | insignificant | not genotyped | insignificant | significant | insignificant | insignificant | not genotyped | not genotyped | insignificant | 1 | 5 | 4 |
| denovoLocus59218 | not genotyped | not genotyped | significant | insignificant | insignificant | insignificant | insignificant | insignificant | not genotyped | not genotyped | 1 | 5 | 4 |
| denovoLocus5952 | not genotyped | not genotyped | significant | insignificant | insignificant | not genotyped | insignificant | insignificant | insignificant | not genotyped | 1 | 5 | 4 |
| denovoLocus6054 | insignificant | insignificant | insignificant | not genotyped | insignificant | not genotyped | not genotyped | significant | insignificant | not genotyped | 1 | 5 | 4 |
| denovoLocus61727 | insignificant | insignificant | significant | not genotyped | not genotyped | insignificant | not genotyped | not genotyped | insignificant | insignificant | 1 | 5 | 4 |
| denovoLocus63184 | not genotyped | not genotyped | insignificant | insignificant | not genotyped | insignificant | significant | not genotyped | insignificant | insignificant | 1 | 5 | 4 |
| denovoLocus6379 | not genotyped | insignificant | not genotyped | significant | insignificant | insignificant | not genotyped | not genotyped | insignificant | insignificant | 1 | 5 | 4 |
| denovoLocus6429 | insignificant | not genotyped | insignificant | not genotyped | insignificant | insignificant | significant | insignificant | not genotyped | not genotyped | 1 | 5 | 4 |
| denovoLocus6666 | not genotyped | not genotyped | insignificant | insignificant | not genotyped | not genotyped | significant | insignificant | insignificant | insignificant | 1 | 5 | 4 |
| denovoLocus7061 | not genotyped | insignificant | not genotyped | insignificant | not genotyped | insignificant | significant | insignificant | insignificant | not genotyped | 1 | 5 | 4 |
| denovoLocus72219 | not genotyped | not genotyped | insignificant | insignificant | not genotyped | insignificant | insignificant | insignificant | not genotyped | significant | 1 | 5 | 4 |
| denovoLocus74709 | not genotyped | not genotyped | insignificant | insignificant | not genotyped | insignificant | significant | insignificant | not genotyped | insignificant | 1 | 5 | 4 |
| denovoLocus74942 | insignificant | not genotyped | insignificant | not genotyped | not genotyped | insignificant | insignificant | not genotyped | insignificant | significant | 1 | 5 | 4 |
| denovoLocus75519 | not genotyped | not genotyped | insignificant | insignificant | not genotyped | insignificant | significant | insignificant | insignificant | not genotyped | 1 | 5 | 4 |
| denovoLocus7668 | insignificant | not genotyped | not genotyped | insignificant | insignificant | not genotyped | insignificant | not genotyped | significant | insignificant | 1 | 5 | 4 |
| denovoLocus77358 | insignificant | not genotyped | insignificant | not genotyped | not genotyped | not genotyped | insignificant | insignificant | significant | insignificant | 1 | 5 | 4 |
| denovoLocus7795 | not genotyped | insignificant | insignificant | insignificant | not genotyped | insignificant | not genotyped | insignificant | not genotyped | significant | 1 | 5 | 4 |
| denovoLocus79020 | not genotyped | not genotyped | not genotyped | insignificant | significant | not genotyped | insignificant | insignificant | insignificant | insignificant | 1 | 5 | 4 |
| denovoLocus80015 | insignificant | insignificant | not genotyped | insignificant | not genotyped | insignificant | not genotyped | significant | insignificant | not genotyped | 1 | 5 | 4 |
| denovoLocus80241 | insignificant | not genotyped | not genotyped | not genotyped | not genotyped | insignificant | insignificant | insignificant | insignificant | significant | 1 | 5 | 4 |
| denovoLocus8027 | insignificant | insignificant | insignificant | not genotyped | not genotyped | significant | not genotyped | not genotyped | insignificant | insignificant | 1 | 5 | 4 |
| denovoLocus81666 | insignificant | not genotyped | insignificant | not genotyped | not genotyped | insignificant | significant | not genotyped | insignificant | insignificant | 1 | 5 | 4 |
| denovoLocus84447 | not genotyped | insignificant | significant | insignificant | not genotyped | insignificant | not genotyped | insignificant | insignificant | not genotyped | 1 | 5 | 4 |
| denovoLocus84672 | insignificant | significant | not genotyped | insignificant | insignificant | insignificant | insignificant | not genotyped | not genotyped | not genotyped | 1 | 5 | 4 |
| denovoLocus8470 | insignificant | not genotyped | not genotyped | insignificant | not genotyped | insignificant | insignificant | insignificant | not genotyped | significant | 1 | 5 | 4 |
| denovoLocus85946 | not genotyped | not genotyped | insignificant | not genotyped | insignificant | insignificant | significant | insignificant | not genotyped | insignificant | 1 | 5 | 4 |
| denovoLocus87143 | insignificant | not genotyped | insignificant | not genotyped | not genotyped | not genotyped | insignificant | insignificant | insignificant | significant | 1 | 5 | 4 |
| denovoLocus8750 | not genotyped | insignificant | not genotyped | not genotyped | insignificant | not genotyped | insignificant | insignificant | significant | insignificant | 1 | 5 | 4 |
| denovoLocus88034 | not genotyped | not genotyped | not genotyped | insignificant | insignificant | insignificant | significant | insignificant | not genotyped | insignificant | 1 | 5 | 4 |
| denovoLocus8819 | not genotyped | insignificant | not genotyped | not genotyped | not genotyped | insignificant | significant | insignificant | insignificant | insignificant | 1 | 5 | 4 |
| denovoLocus89433 | not genotyped | not genotyped | not genotyped | insignificant | insignificant | not genotyped | significant | insignificant | insignificant | insignificant | 1 | 5 | 4 |
| denovoLocus9041 | not genotyped | insignificant | not genotyped | insignificant | insignificant | not genotyped | not genotyped | insignificant | significant | insignificant | 1 | 5 | 4 |
| denovoLocus955 | insignificant | not genotyped | insignificant | insignificant | not genotyped | not genotyped | significant | not genotyped | insignificant | insignificant | 1 | 5 | 4 |
| denovoLocus9798 | insignificant | insignificant | not genotyped | insignificant | insignificant | insignificant | not genotyped | not genotyped | not genotyped | significant | 1 | 5 | 4 |
| denovoLocus9909 | insignificant | insignificant | insignificant | not genotyped | not genotyped | not genotyped | significant | not genotyped | insignificant | insignificant | 1 | 5 | 4 |
| denovoLocus9932 | insignificant | not genotyped | not genotyped | insignificant | significant | insignificant | not genotyped | insignificant | not genotyped | insignificant | 1 | 5 | 4 |
| denovoLocus104535 | insignificant | insignificant | not genotyped | not genotyped | not genotyped | insignificant | not genotyped | significant | insignificant | not genotyped | 1 | 4 | 5 |
| denovoLocus10461 | insignificant | not genotyped | insignificant | not genotyped | significant | not genotyped | insignificant | insignificant | not genotyped | not genotyped | 1 | 4 | 5 |
| denovoLocus10488 | not genotyped | insignificant | insignificant | not genotyped | not genotyped | insignificant | not genotyped | not genotyped | insignificant | significant | 1 | 4 | 5 |
| denovoLocus10535 | insignificant | not genotyped | insignificant | insignificant | significant | not genotyped | not genotyped | insignificant | not genotyped | not genotyped | 1 | 4 | 5 |
| denovoLocus105634 | significant | insignificant | not genotyped | not genotyped | not genotyped | insignificant | not genotyped | not genotyped | insignificant | insignificant | 1 | 4 | 5 |
| denovoLocus11023 | not genotyped | insignificant | not genotyped | insignificant | not genotyped | significant | not genotyped | insignificant | insignificant | not genotyped | 1 | 4 | 5 |
| denovoLocus11039 | not genotyped | insignificant | not genotyped | not genotyped | insignificant | not genotyped | not genotyped | significant | insignificant | insignificant | 1 | 4 | 5 |
| denovoLocus112583 | not genotyped | not genotyped | significant | not genotyped | not genotyped | insignificant | not genotyped | insignificant | insignificant | insignificant | 1 | 4 | 5 |
| denovoLocus11285 | not genotyped | not genotyped | not genotyped | insignificant | insignificant | insignificant | not genotyped | significant | insignificant | not genotyped | 1 | 4 | 5 |
| denovoLocus11416 | not genotyped | not genotyped | not genotyped | insignificant | not genotyped | insignificant | significant | insignificant | not genotyped | insignificant | 1 | 4 | 5 |
| denovoLocus11418 | not genotyped | significant | not genotyped | insignificant | insignificant | insignificant | not genotyped | not genotyped | not genotyped | insignificant | 1 | 4 | 5 |
| denovoLocus11769 | insignificant | not genotyped | not genotyped | not genotyped | significant | insignificant | not genotyped | insignificant | not genotyped | insignificant | 1 | 4 | 5 |
| denovoLocus11806 | insignificant | insignificant | insignificant | not genotyped | not genotyped | not genotyped | insignificant | not genotyped | not genotyped | significant | 1 | 4 | 5 |
| denovoLocus12850 | not genotyped | not genotyped | insignificant | insignificant | significant | not genotyped | not genotyped | insignificant | insignificant | not genotyped | 1 | 4 | 5 |
| denovoLocus12916 | not genotyped | insignificant | significant | insignificant | not genotyped | insignificant | not genotyped | not genotyped | insignificant | not genotyped | 1 | 4 | 5 |
| denovoLocus13182 | insignificant | insignificant | not genotyped | insignificant | insignificant | not genotyped | not genotyped | significant | not genotyped | not genotyped | 1 | 4 | 5 |
| denovoLocus13237 | insignificant | not genotyped | not genotyped | insignificant | insignificant | not genotyped | significant | insignificant | not genotyped | not genotyped | 1 | 4 | 5 |
| denovoLocus14309 | insignificant | insignificant | not genotyped | not genotyped | insignificant | not genotyped | not genotyped | not genotyped | insignificant | significant | 1 | 4 | 5 |
| denovoLocus14432 | not genotyped | insignificant | not genotyped | not genotyped | insignificant | not genotyped | significant | insignificant | insignificant | not genotyped | 1 | 4 | 5 |
| denovoLocus14876 | insignificant | insignificant | not genotyped | significant | not genotyped | insignificant | not genotyped | insignificant | not genotyped | not genotyped | 1 | 4 | 5 |
| denovoLocus1531 | insignificant | insignificant | not genotyped | insignificant | insignificant | not genotyped | significant | not genotyped | not genotyped | not genotyped | 1 | 4 | 5 |
| denovoLocus16431 | insignificant | not genotyped | not genotyped | not genotyped | insignificant | not genotyped | not genotyped | insignificant | insignificant | significant | 1 | 4 | 5 |
| denovoLocus16667 | not genotyped | insignificant | significant | not genotyped | insignificant | insignificant | not genotyped | not genotyped | not genotyped | insignificant | 1 | 4 | 5 |
| denovoLocus17084 | insignificant | insignificant | not genotyped | insignificant | not genotyped | not genotyped | significant | not genotyped | insignificant | not genotyped | 1 | 4 | 5 |
| denovoLocus18137 | not genotyped | insignificant | not genotyped | not genotyped | insignificant | not genotyped | not genotyped | insignificant | insignificant | significant | 1 | 4 | 5 |
| denovoLocus18164 | insignificant | not genotyped | insignificant | not genotyped | significant | insignificant | insignificant | not genotyped | not genotyped | not genotyped | 1 | 4 | 5 |
| denovoLocus18936 | insignificant | not genotyped | not genotyped | insignificant | not genotyped | not genotyped | significant | not genotyped | insignificant | insignificant | 1 | 4 | 5 |
| denovoLocus19402 | insignificant | insignificant | not genotyped | significant | not genotyped | not genotyped | not genotyped | not genotyped | insignificant | insignificant | 1 | 4 | 5 |
| denovoLocus19618 | insignificant | insignificant | not genotyped | not genotyped | not genotyped | not genotyped | significant | insignificant | insignificant | not genotyped | 1 | 4 | 5 |
| denovoLocus1982 | not genotyped | not genotyped | insignificant | insignificant | not genotyped | insignificant | insignificant | not genotyped | not genotyped | significant | 1 | 4 | 5 |
| denovoLocus20347 | not genotyped | insignificant | not genotyped | not genotyped | not genotyped | not genotyped | insignificant | insignificant | significant | insignificant | 1 | 4 | 5 |
| denovoLocus20484 | significant | not genotyped | not genotyped | insignificant | insignificant | insignificant | not genotyped | not genotyped | insignificant | not genotyped | 1 | 4 | 5 |
| denovoLocus21207 | insignificant | not genotyped | insignificant | not genotyped | not genotyped | insignificant | not genotyped | insignificant | significant | not genotyped | 1 | 4 | 5 |
| denovoLocus2427 | insignificant | not genotyped | insignificant | insignificant | not genotyped | insignificant | significant | not genotyped | not genotyped | not genotyped | 1 | 4 | 5 |
| denovoLocus24476 | insignificant | not genotyped | insignificant | insignificant | not genotyped | insignificant | significant | not genotyped | not genotyped | not genotyped | 1 | 4 | 5 |
| denovoLocus24908 | insignificant | not genotyped | not genotyped | not genotyped | insignificant | insignificant | significant | insignificant | not genotyped | not genotyped | 1 | 4 | 5 |
| denovoLocus25290 | not genotyped | not genotyped | insignificant | insignificant | not genotyped | insignificant | significant | not genotyped | not genotyped | insignificant | 1 | 4 | 5 |
| denovoLocus2557 | not genotyped | not genotyped | insignificant | insignificant | insignificant | insignificant | significant | not genotyped | not genotyped | not genotyped | 1 | 4 | 5 |
| denovoLocus26012 | significant | insignificant | not genotyped | not genotyped | not genotyped | insignificant | insignificant | not genotyped | insignificant | not genotyped | 1 | 4 | 5 |
| denovoLocus26953 | insignificant | insignificant | insignificant | not genotyped | insignificant | not genotyped | not genotyped | not genotyped | not genotyped | significant | 1 | 4 | 5 |
| denovoLocus27429 | insignificant | insignificant | not genotyped | not genotyped | insignificant | not genotyped | not genotyped | not genotyped | insignificant | significant | 1 | 4 | 5 |
| denovoLocus27887 | insignificant | insignificant | not genotyped | not genotyped | not genotyped | significant | insignificant | insignificant | not genotyped | not genotyped | 1 | 4 | 5 |
| denovoLocus29540 | not genotyped | not genotyped | significant | not genotyped | not genotyped | insignificant | not genotyped | insignificant | insignificant | insignificant | 1 | 4 | 5 |
| denovoLocus29790 | insignificant | insignificant | not genotyped | not genotyped | insignificant | not genotyped | not genotyped | insignificant | not genotyped | significant | 1 | 4 | 5 |
| denovoLocus30195 | insignificant | not genotyped | not genotyped | insignificant | not genotyped | not genotyped | significant | not genotyped | insignificant | insignificant | 1 | 4 | 5 |
| denovoLocus32712 | not genotyped | not genotyped | insignificant | insignificant | insignificant | not genotyped | not genotyped | insignificant | not genotyped | significant | 1 | 4 | 5 |
| denovoLocus32783 | not genotyped | insignificant | not genotyped | insignificant | insignificant | not genotyped | insignificant | not genotyped | significant | not genotyped | 1 | 4 | 5 |
| denovoLocus32856 | insignificant | insignificant | insignificant | not genotyped | not genotyped | not genotyped | not genotyped | not genotyped | significant | insignificant | 1 | 4 | 5 |
| denovoLocus33055 | not genotyped | not genotyped | not genotyped | insignificant | insignificant | not genotyped | significant | insignificant | insignificant | not genotyped | 1 | 4 | 5 |
| denovoLocus34309 | not genotyped | insignificant | not genotyped | not genotyped | insignificant | not genotyped | significant | insignificant | insignificant | not genotyped | 1 | 4 | 5 |
| denovoLocus34714 | insignificant | insignificant | not genotyped | insignificant | significant | not genotyped | not genotyped | not genotyped | not genotyped | insignificant | 1 | 4 | 5 |
| denovoLocus3788 | insignificant | insignificant | not genotyped | not genotyped | not genotyped | not genotyped | significant | not genotyped | insignificant | insignificant | 1 | 4 | 5 |
| denovoLocus38305 | not genotyped | not genotyped | not genotyped | insignificant | not genotyped | insignificant | insignificant | significant | insignificant | not genotyped | 1 | 4 | 5 |
| denovoLocus38919 | insignificant | insignificant | not genotyped | not genotyped | insignificant | not genotyped | insignificant | significant | not genotyped | not genotyped | 1 | 4 | 5 |
| denovoLocus39040 | insignificant | not genotyped | insignificant | significant | insignificant | not genotyped | insignificant | not genotyped | not genotyped | not genotyped | 1 | 4 | 5 |
| denovoLocus39579 | significant | not genotyped | insignificant | insignificant | not genotyped | insignificant | not genotyped | insignificant | not genotyped | not genotyped | 1 | 4 | 5 |
| denovoLocus41135 | not genotyped | insignificant | not genotyped | not genotyped | not genotyped | insignificant | significant | not genotyped | insignificant | insignificant | 1 | 4 | 5 |
| denovoLocus42786 | insignificant | not genotyped | significant | not genotyped | insignificant | not genotyped | not genotyped | insignificant | not genotyped | insignificant | 1 | 4 | 5 |
| denovoLocus4290 | significant | not genotyped | insignificant | insignificant | not genotyped | insignificant | not genotyped | not genotyped | not genotyped | insignificant | 1 | 4 | 5 |
| denovoLocus43001 | insignificant | not genotyped | not genotyped | not genotyped | insignificant | not genotyped | significant | not genotyped | insignificant | insignificant | 1 | 4 | 5 |
| denovoLocus46627 | not genotyped | not genotyped | insignificant | significant | insignificant | not genotyped | insignificant | not genotyped | not genotyped | insignificant | 1 | 4 | 5 |
| denovoLocus47003 | insignificant | not genotyped | significant | not genotyped | insignificant | insignificant | not genotyped | not genotyped | insignificant | not genotyped | 1 | 4 | 5 |
| denovoLocus53089 | not genotyped | insignificant | not genotyped | significant | insignificant | not genotyped | not genotyped | insignificant | insignificant | not genotyped | 1 | 4 | 5 |
| denovoLocus55286 | not genotyped | not genotyped | not genotyped | insignificant | not genotyped | insignificant | significant | insignificant | insignificant | not genotyped | 1 | 4 | 5 |
| denovoLocus55370 | not genotyped | insignificant | not genotyped | insignificant | not genotyped | insignificant | significant | insignificant | not genotyped | not genotyped | 1 | 4 | 5 |
| denovoLocus56576 | not genotyped | not genotyped | not genotyped | insignificant | not genotyped | insignificant | significant | insignificant | insignificant | not genotyped | 1 | 4 | 5 |
| denovoLocus56873 | not genotyped | insignificant | significant | not genotyped | insignificant | not genotyped | not genotyped | insignificant | not genotyped | insignificant | 1 | 4 | 5 |
| denovoLocus57601 | insignificant | not genotyped | insignificant | not genotyped | not genotyped | insignificant | significant | not genotyped | not genotyped | insignificant | 1 | 4 | 5 |
| denovoLocus57639 | insignificant | not genotyped | not genotyped | not genotyped | not genotyped | not genotyped | significant | insignificant | insignificant | insignificant | 1 | 4 | 5 |
| denovoLocus5774 | significant | insignificant | insignificant | not genotyped | not genotyped | not genotyped | not genotyped | insignificant | insignificant | not genotyped | 1 | 4 | 5 |
| denovoLocus59191 | insignificant | not genotyped | insignificant | not genotyped | insignificant | not genotyped | significant | insignificant | not genotyped | not genotyped | 1 | 4 | 5 |
| denovoLocus59342 | insignificant | not genotyped | not genotyped | insignificant | not genotyped | not genotyped | significant | insignificant | insignificant | not genotyped | 1 | 4 | 5 |
| denovoLocus6378 | not genotyped | insignificant | insignificant | not genotyped | insignificant | not genotyped | significant | insignificant | not genotyped | not genotyped | 1 | 4 | 5 |
| denovoLocus67645 | insignificant | not genotyped | not genotyped | significant | insignificant | insignificant | not genotyped | not genotyped | not genotyped | insignificant | 1 | 4 | 5 |
| denovoLocus70549 | insignificant | not genotyped | significant | insignificant | insignificant | not genotyped | not genotyped | not genotyped | insignificant | not genotyped | 1 | 4 | 5 |
| denovoLocus71001 | not genotyped | not genotyped | significant | not genotyped | insignificant | not genotyped | insignificant | insignificant | insignificant | not genotyped | 1 | 4 | 5 |
| denovoLocus71641 | insignificant | not genotyped | insignificant | insignificant | not genotyped | not genotyped | insignificant | significant | not genotyped | not genotyped | 1 | 4 | 5 |
| denovoLocus7372 | insignificant | insignificant | not genotyped | not genotyped | insignificant | not genotyped | significant | not genotyped | not genotyped | insignificant | 1 | 4 | 5 |
| denovoLocus74 | not genotyped | insignificant | not genotyped | insignificant | not genotyped | not genotyped | insignificant | insignificant | not genotyped | significant | 1 | 4 | 5 |
| denovoLocus7429 | not genotyped | not genotyped | insignificant | insignificant | not genotyped | not genotyped | insignificant | insignificant | significant | not genotyped | 1 | 4 | 5 |
| denovoLocus74742 | not genotyped | insignificant | insignificant | not genotyped | insignificant | not genotyped | not genotyped | not genotyped | insignificant | significant | 1 | 4 | 5 |
| denovoLocus7487 | not genotyped | insignificant | not genotyped | insignificant | insignificant | insignificant | not genotyped | not genotyped | significant | not genotyped | 1 | 4 | 5 |
| denovoLocus76660 | insignificant | not genotyped | not genotyped | significant | insignificant | not genotyped | insignificant | insignificant | not genotyped | not genotyped | 1 | 4 | 5 |
| denovoLocus7677 | insignificant | not genotyped | not genotyped | insignificant | insignificant | not genotyped | significant | not genotyped | not genotyped | insignificant | 1 | 4 | 5 |
| denovoLocus7880 | not genotyped | insignificant | insignificant | insignificant | not genotyped | not genotyped | not genotyped | insignificant | not genotyped | significant | 1 | 4 | 5 |
| denovoLocus82712 | insignificant | not genotyped | insignificant | insignificant | significant | not genotyped | not genotyped | not genotyped | insignificant | not genotyped | 1 | 4 | 5 |
| denovoLocus84564 | insignificant | insignificant | insignificant | not genotyped | not genotyped | insignificant | significant | not genotyped | not genotyped | not genotyped | 1 | 4 | 5 |
| denovoLocus84762 | not genotyped | not genotyped | not genotyped | insignificant | insignificant | not genotyped | significant | insignificant | not genotyped | insignificant | 1 | 4 | 5 |
| denovoLocus85139 | insignificant | not genotyped | insignificant | not genotyped | not genotyped | insignificant | insignificant | significant | not genotyped | not genotyped | 1 | 4 | 5 |
| denovoLocus8587 | insignificant | insignificant | not genotyped | not genotyped | insignificant | not genotyped | significant | insignificant | not genotyped | not genotyped | 1 | 4 | 5 |
| denovoLocus86845 | insignificant | not genotyped | not genotyped | insignificant | insignificant | not genotyped | significant | not genotyped | not genotyped | insignificant | 1 | 4 | 5 |
| denovoLocus9073 | insignificant | not genotyped | insignificant | not genotyped | not genotyped | significant | insignificant | insignificant | not genotyped | not genotyped | 1 | 4 | 5 |
| denovoLocus9709 | insignificant | not genotyped | not genotyped | not genotyped | not genotyped | insignificant | significant | not genotyped | insignificant | insignificant | 1 | 4 | 5 |
| denovoLocus9757 | not genotyped | insignificant | insignificant | not genotyped | not genotyped | insignificant | significant | not genotyped | not genotyped | insignificant | 1 | 4 | 5 |
| denovoLocus9852 | not genotyped | insignificant | significant | not genotyped | insignificant | not genotyped | insignificant | insignificant | not genotyped | not genotyped | 1 | 4 | 5 |
| denovoLocus990 | not genotyped | insignificant | insignificant | not genotyped | not genotyped | not genotyped | insignificant | insignificant | significant | not genotyped | 1 | 4 | 5 |
| denovoLocus99667 | insignificant | not genotyped | not genotyped | not genotyped | insignificant | not genotyped | insignificant | not genotyped | significant | insignificant | 1 | 4 | 5 |
| denovoLocus10060 | not genotyped | not genotyped | not genotyped | insignificant | not genotyped | not genotyped | significant | not genotyped | insignificant | insignificant | 1 | 3 | 6 |
| denovoLocus10072 | insignificant | not genotyped | not genotyped | insignificant | insignificant | not genotyped | significant | not genotyped | not genotyped | not genotyped | 1 | 3 | 6 |
| denovoLocus107223 | insignificant | not genotyped | not genotyped | insignificant | not genotyped | not genotyped | significant | insignificant | not genotyped | not genotyped | 1 | 3 | 6 |
| denovoLocus107423 | insignificant | insignificant | significant | not genotyped | not genotyped | not genotyped | not genotyped | not genotyped | not genotyped | insignificant | 1 | 3 | 6 |
| denovoLocus107467 | not genotyped | insignificant | not genotyped | insignificant | not genotyped | insignificant | significant | not genotyped | not genotyped | not genotyped | 1 | 3 | 6 |
| denovoLocus12583 | not genotyped | not genotyped | insignificant | insignificant | not genotyped | not genotyped | significant | insignificant | not genotyped | not genotyped | 1 | 3 | 6 |
| denovoLocus12733 | not genotyped | not genotyped | not genotyped | not genotyped | insignificant | insignificant | significant | insignificant | not genotyped | not genotyped | 1 | 3 | 6 |
| denovoLocus13276 | not genotyped | not genotyped | not genotyped | insignificant | not genotyped | significant | insignificant | insignificant | not genotyped | not genotyped | 1 | 3 | 6 |
| denovoLocus14317 | not genotyped | not genotyped | not genotyped | insignificant | insignificant | not genotyped | insignificant | significant | not genotyped | not genotyped | 1 | 3 | 6 |
| denovoLocus14417 | significant | insignificant | not genotyped | not genotyped | not genotyped | not genotyped | not genotyped | insignificant | insignificant | not genotyped | 1 | 3 | 6 |
| denovoLocus14584 | not genotyped | insignificant | insignificant | not genotyped | significant | not genotyped | not genotyped | not genotyped | not genotyped | insignificant | 1 | 3 | 6 |
| denovoLocus1495 | not genotyped | not genotyped | not genotyped | not genotyped | insignificant | not genotyped | significant | not genotyped | insignificant | insignificant | 1 | 3 | 6 |
| denovoLocus15769 | significant | not genotyped | not genotyped | not genotyped | not genotyped | insignificant | not genotyped | not genotyped | insignificant | insignificant | 1 | 3 | 6 |
| denovoLocus15985 | insignificant | not genotyped | not genotyped | insignificant | not genotyped | not genotyped | insignificant | not genotyped | not genotyped | significant | 1 | 3 | 6 |
| denovoLocus16675 | insignificant | significant | not genotyped | not genotyped | insignificant | not genotyped | not genotyped | not genotyped | not genotyped | insignificant | 1 | 3 | 6 |
| denovoLocus16735 | not genotyped | insignificant | not genotyped | not genotyped | insignificant | not genotyped | significant | not genotyped | insignificant | not genotyped | 1 | 3 | 6 |
| denovoLocus17525 | not genotyped | not genotyped | not genotyped | insignificant | insignificant | insignificant | significant | not genotyped | not genotyped | not genotyped | 1 | 3 | 6 |
| denovoLocus17758 | not genotyped | not genotyped | not genotyped | insignificant | not genotyped | insignificant | significant | not genotyped | not genotyped | insignificant | 1 | 3 | 6 |
| denovoLocus17939 | not genotyped | not genotyped | not genotyped | insignificant | insignificant | insignificant | significant | not genotyped | not genotyped | not genotyped | 1 | 3 | 6 |
| denovoLocus18288 | not genotyped | not genotyped | not genotyped | insignificant | not genotyped | insignificant | significant | not genotyped | insignificant | not genotyped | 1 | 3 | 6 |
| denovoLocus18292 | insignificant | not genotyped | significant | not genotyped | insignificant | not genotyped | not genotyped | not genotyped | insignificant | not genotyped | 1 | 3 | 6 |
| denovoLocus18728 | insignificant | not genotyped | not genotyped | not genotyped | not genotyped | insignificant | significant | not genotyped | insignificant | not genotyped | 1 | 3 | 6 |
| denovoLocus1969 | not genotyped | not genotyped | not genotyped | significant | not genotyped | insignificant | insignificant | insignificant | not genotyped | not genotyped | 1 | 3 | 6 |
| denovoLocus19736 | insignificant | not genotyped | insignificant | not genotyped | not genotyped | not genotyped | significant | insignificant | not genotyped | not genotyped | 1 | 3 | 6 |
| denovoLocus23064 | insignificant | significant | insignificant | not genotyped | not genotyped | not genotyped | not genotyped | not genotyped | not genotyped | insignificant | 1 | 3 | 6 |
| denovoLocus23737 | insignificant | not genotyped | not genotyped | not genotyped | not genotyped | insignificant | significant | not genotyped | insignificant | not genotyped | 1 | 3 | 6 |
| denovoLocus24488 | not genotyped | not genotyped | not genotyped | insignificant | not genotyped | insignificant | significant | insignificant | not genotyped | not genotyped | 1 | 3 | 6 |
| denovoLocus2465 | not genotyped | insignificant | not genotyped | insignificant | not genotyped | insignificant | significant | not genotyped | not genotyped | not genotyped | 1 | 3 | 6 |
| denovoLocus2482 | not genotyped | insignificant | not genotyped | not genotyped | not genotyped | not genotyped | significant | insignificant | insignificant | not genotyped | 1 | 3 | 6 |
| denovoLocus24999 | significant | insignificant | not genotyped | not genotyped | insignificant | insignificant | not genotyped | not genotyped | not genotyped | not genotyped | 1 | 3 | 6 |
| denovoLocus25207 | insignificant | not genotyped | insignificant | not genotyped | significant | not genotyped | insignificant | not genotyped | not genotyped | not genotyped | 1 | 3 | 6 |
| denovoLocus2682 | not genotyped | insignificant | insignificant | not genotyped | significant | not genotyped | not genotyped | insignificant | not genotyped | not genotyped | 1 | 3 | 6 |
| denovoLocus28013 | insignificant | not genotyped | not genotyped | insignificant | not genotyped | insignificant | significant | not genotyped | not genotyped | not genotyped | 1 | 3 | 6 |
| denovoLocus2846 | not genotyped | not genotyped | not genotyped | not genotyped | insignificant | not genotyped | significant | insignificant | insignificant | not genotyped | 1 | 3 | 6 |
| denovoLocus28673 | not genotyped | not genotyped | insignificant | not genotyped | insignificant | insignificant | significant | not genotyped | not genotyped | not genotyped | 1 | 3 | 6 |
| denovoLocus2949 | insignificant | insignificant | not genotyped | not genotyped | insignificant | not genotyped | not genotyped | not genotyped | significant | not genotyped | 1 | 3 | 6 |
| denovoLocus29738 | insignificant | not genotyped | not genotyped | not genotyped | insignificant | not genotyped | insignificant | not genotyped | not genotyped | significant | 1 | 3 | 6 |
| denovoLocus29768 | not genotyped | not genotyped | insignificant | not genotyped | not genotyped | insignificant | significant | insignificant | not genotyped | not genotyped | 1 | 3 | 6 |
| denovoLocus2995 | not genotyped | not genotyped | not genotyped | insignificant | insignificant | not genotyped | not genotyped | significant | insignificant | not genotyped | 1 | 3 | 6 |
| denovoLocus31097 | significant | insignificant | not genotyped | insignificant | not genotyped | not genotyped | not genotyped | not genotyped | not genotyped | insignificant | 1 | 3 | 6 |
| denovoLocus31756 | not genotyped | not genotyped | not genotyped | not genotyped | insignificant | insignificant | significant | not genotyped | insignificant | not genotyped | 1 | 3 | 6 |
| denovoLocus33048 | not genotyped | not genotyped | not genotyped | insignificant | not genotyped | insignificant | significant | insignificant | not genotyped | not genotyped | 1 | 3 | 6 |
| denovoLocus35589 | not genotyped | insignificant | not genotyped | significant | insignificant | not genotyped | not genotyped | insignificant | not genotyped | not genotyped | 1 | 3 | 6 |
| denovoLocus35928 | not genotyped | insignificant | not genotyped | not genotyped | not genotyped | not genotyped | significant | not genotyped | insignificant | insignificant | 1 | 3 | 6 |
| denovoLocus36169 | insignificant | insignificant | not genotyped | not genotyped | insignificant | not genotyped | not genotyped | not genotyped | not genotyped | significant | 1 | 3 | 6 |
| denovoLocus3645 | insignificant | not genotyped | not genotyped | not genotyped | not genotyped | not genotyped | significant | insignificant | insignificant | not genotyped | 1 | 3 | 6 |
| denovoLocus37426 | not genotyped | insignificant | not genotyped | insignificant | not genotyped | significant | not genotyped | not genotyped | not genotyped | insignificant | 1 | 3 | 6 |
| denovoLocus38074 | not genotyped | not genotyped | not genotyped | not genotyped | not genotyped | not genotyped | insignificant | insignificant | insignificant | significant | 1 | 3 | 6 |
| denovoLocus39039 | insignificant | insignificant | not genotyped | not genotyped | not genotyped | not genotyped | significant | not genotyped | not genotyped | insignificant | 1 | 3 | 6 |
| denovoLocus39460 | insignificant | insignificant | not genotyped | not genotyped | not genotyped | insignificant | not genotyped | not genotyped | significant | not genotyped | 1 | 3 | 6 |
| denovoLocus395 | not genotyped | insignificant | not genotyped | significant | not genotyped | not genotyped | not genotyped | not genotyped | insignificant | insignificant | 1 | 3 | 6 |
| denovoLocus4023 | insignificant | not genotyped | insignificant | not genotyped | not genotyped | not genotyped | significant | insignificant | not genotyped | not genotyped | 1 | 3 | 6 |
| denovoLocus40319 | insignificant | not genotyped | not genotyped | not genotyped | insignificant | not genotyped | significant | insignificant | not genotyped | not genotyped | 1 | 3 | 6 |
| denovoLocus40591 | insignificant | not genotyped | not genotyped | not genotyped | insignificant | insignificant | significant | not genotyped | not genotyped | not genotyped | 1 | 3 | 6 |
| denovoLocus4264 | insignificant | insignificant | not genotyped | not genotyped | not genotyped | not genotyped | not genotyped | not genotyped | insignificant | significant | 1 | 3 | 6 |
| denovoLocus43023 | not genotyped | not genotyped | not genotyped | insignificant | insignificant | not genotyped | significant | not genotyped | insignificant | not genotyped | 1 | 3 | 6 |
| denovoLocus440 | significant | insignificant | not genotyped | not genotyped | not genotyped | not genotyped | not genotyped | not genotyped | insignificant | insignificant | 1 | 3 | 6 |
| denovoLocus451 | not genotyped | insignificant | not genotyped | insignificant | not genotyped | not genotyped | significant | not genotyped | insignificant | not genotyped | 1 | 3 | 6 |
| denovoLocus4639 | not genotyped | not genotyped | not genotyped | insignificant | insignificant | insignificant | significant | not genotyped | not genotyped | not genotyped | 1 | 3 | 6 |
| denovoLocus46419 | not genotyped | insignificant | not genotyped | not genotyped | not genotyped | significant | not genotyped | not genotyped | insignificant | insignificant | 1 | 3 | 6 |
| denovoLocus46438 | insignificant | not genotyped | not genotyped | not genotyped | significant | insignificant | not genotyped | not genotyped | insignificant | not genotyped | 1 | 3 | 6 |
| denovoLocus47770 | not genotyped | insignificant | not genotyped | not genotyped | insignificant | not genotyped | not genotyped | not genotyped | significant | insignificant | 1 | 3 | 6 |
| denovoLocus50438 | not genotyped | not genotyped | insignificant | not genotyped | insignificant | not genotyped | not genotyped | not genotyped | significant | insignificant | 1 | 3 | 6 |
| denovoLocus50816 | not genotyped | insignificant | insignificant | insignificant | not genotyped | not genotyped | significant | not genotyped | not genotyped | not genotyped | 1 | 3 | 6 |
| denovoLocus53541 | not genotyped | insignificant | not genotyped | not genotyped | insignificant | insignificant | significant | not genotyped | not genotyped | not genotyped | 1 | 3 | 6 |
| denovoLocus54245 | insignificant | not genotyped | not genotyped | not genotyped | insignificant | insignificant | significant | not genotyped | not genotyped | not genotyped | 1 | 3 | 6 |
| denovoLocus54301 | not genotyped | not genotyped | not genotyped | insignificant | insignificant | not genotyped | not genotyped | insignificant | significant | not genotyped | 1 | 3 | 6 |
| denovoLocus56139 | not genotyped | not genotyped | not genotyped | insignificant | significant | insignificant | not genotyped | not genotyped | not genotyped | insignificant | 1 | 3 | 6 |
| denovoLocus56552 | not genotyped | not genotyped | not genotyped | insignificant | insignificant | not genotyped | significant | insignificant | not genotyped | not genotyped | 1 | 3 | 6 |
| denovoLocus56759 | not genotyped | insignificant | not genotyped | not genotyped | insignificant | not genotyped | significant | not genotyped | insignificant | not genotyped | 1 | 3 | 6 |
| denovoLocus58 | not genotyped | insignificant | not genotyped | not genotyped | not genotyped | insignificant | not genotyped | not genotyped | insignificant | significant | 1 | 3 | 6 |
| denovoLocus58940 | not genotyped | not genotyped | insignificant | not genotyped | not genotyped | insignificant | not genotyped | insignificant | significant | not genotyped | 1 | 3 | 6 |
| denovoLocus6020 | not genotyped | insignificant | not genotyped | insignificant | not genotyped | not genotyped | significant | insignificant | not genotyped | not genotyped | 1 | 3 | 6 |
| denovoLocus60275 | not genotyped | not genotyped | not genotyped | insignificant | not genotyped | insignificant | significant | insignificant | not genotyped | not genotyped | 1 | 3 | 6 |
| denovoLocus60661 | insignificant | not genotyped | not genotyped | insignificant | insignificant | not genotyped | significant | not genotyped | not genotyped | not genotyped | 1 | 3 | 6 |
| denovoLocus62287 | not genotyped | insignificant | insignificant | not genotyped | insignificant | not genotyped | significant | not genotyped | not genotyped | not genotyped | 1 | 3 | 6 |
| denovoLocus64024 | not genotyped | insignificant | insignificant | not genotyped | not genotyped | not genotyped | not genotyped | insignificant | significant | not genotyped | 1 | 3 | 6 |
| denovoLocus6435 | not genotyped | not genotyped | not genotyped | insignificant | not genotyped | insignificant | significant | insignificant | not genotyped | not genotyped | 1 | 3 | 6 |
| denovoLocus6469 | not genotyped | not genotyped | not genotyped | not genotyped | insignificant | insignificant | not genotyped | not genotyped | significant | insignificant | 1 | 3 | 6 |
| denovoLocus6550 | not genotyped | not genotyped | not genotyped | insignificant | insignificant | not genotyped | significant | not genotyped | not genotyped | insignificant | 1 | 3 | 6 |
| denovoLocus66657 | not genotyped | not genotyped | not genotyped | not genotyped | insignificant | insignificant | not genotyped | insignificant | significant | not genotyped | 1 | 3 | 6 |
| denovoLocus67483 | insignificant | not genotyped | not genotyped | not genotyped | insignificant | not genotyped | significant | not genotyped | not genotyped | insignificant | 1 | 3 | 6 |
| denovoLocus68124 | not genotyped | not genotyped | not genotyped | not genotyped | not genotyped | insignificant | significant | insignificant | insignificant | not genotyped | 1 | 3 | 6 |
| denovoLocus68825 | not genotyped | insignificant | insignificant | not genotyped | not genotyped | not genotyped | insignificant | not genotyped | significant | not genotyped | 1 | 3 | 6 |
| denovoLocus69290 | insignificant | insignificant | insignificant | not genotyped | not genotyped | significant | not genotyped | not genotyped | not genotyped | not genotyped | 1 | 3 | 6 |
| denovoLocus69605 | not genotyped | insignificant | not genotyped | not genotyped | significant | not genotyped | insignificant | not genotyped | insignificant | not genotyped | 1 | 3 | 6 |
| denovoLocus70856 | not genotyped | insignificant | not genotyped | not genotyped | significant | not genotyped | insignificant | not genotyped | not genotyped | insignificant | 1 | 3 | 6 |
| denovoLocus71205 | not genotyped | not genotyped | not genotyped | not genotyped | not genotyped | insignificant | insignificant | significant | insignificant | not genotyped | 1 | 3 | 6 |
| denovoLocus71226 | insignificant | not genotyped | not genotyped | insignificant | not genotyped | not genotyped | not genotyped | insignificant | significant | not genotyped | 1 | 3 | 6 |
| denovoLocus71971 | insignificant | not genotyped | not genotyped | insignificant | not genotyped | insignificant | significant | not genotyped | not genotyped | not genotyped | 1 | 3 | 6 |
| denovoLocus72125 | not genotyped | not genotyped | not genotyped | insignificant | insignificant | not genotyped | not genotyped | insignificant | significant | not genotyped | 1 | 3 | 6 |
| denovoLocus72918 | not genotyped | insignificant | not genotyped | not genotyped | insignificant | insignificant | significant | not genotyped | not genotyped | not genotyped | 1 | 3 | 6 |
| denovoLocus74282 | not genotyped | insignificant | insignificant | not genotyped | not genotyped | insignificant | significant | not genotyped | not genotyped | not genotyped | 1 | 3 | 6 |
| denovoLocus7600 | not genotyped | not genotyped | not genotyped | not genotyped | insignificant | insignificant | significant | insignificant | not genotyped | not genotyped | 1 | 3 | 6 |
| denovoLocus8523 | insignificant | insignificant | significant | not genotyped | not genotyped | not genotyped | not genotyped | not genotyped | insignificant | not genotyped | 1 | 3 | 6 |
| denovoLocus8922 | insignificant | not genotyped | not genotyped | not genotyped | not genotyped | insignificant | significant | not genotyped | insignificant | not genotyped | 1 | 3 | 6 |
| denovoLocus100702 | insignificant | not genotyped | not genotyped | not genotyped | not genotyped | not genotyped | not genotyped | insignificant | not genotyped | significant | 1 | 2 | 7 |
| denovoLocus10175 | not genotyped | not genotyped | not genotyped | significant | not genotyped | insignificant | not genotyped | not genotyped | insignificant | not genotyped | 1 | 2 | 7 |
| denovoLocus10619 | significant | not genotyped | insignificant | not genotyped | not genotyped | insignificant | not genotyped | not genotyped | not genotyped | not genotyped | 1 | 2 | 7 |
| denovoLocus10906 | not genotyped | not genotyped | not genotyped | insignificant | not genotyped | not genotyped | significant | not genotyped | insignificant | not genotyped | 1 | 2 | 7 |
| denovoLocus1118 | not genotyped | not genotyped | not genotyped | not genotyped | insignificant | not genotyped | insignificant | significant | not genotyped | not genotyped | 1 | 2 | 7 |
| denovoLocus115068 | not genotyped | insignificant | not genotyped | insignificant | not genotyped | not genotyped | not genotyped | not genotyped | not genotyped | significant | 1 | 2 | 7 |
| denovoLocus116503 | not genotyped | not genotyped | not genotyped | insignificant | insignificant | not genotyped | significant | not genotyped | not genotyped | not genotyped | 1 | 2 | 7 |
| denovoLocus12657 | insignificant | not genotyped | not genotyped | not genotyped | not genotyped | insignificant | not genotyped | not genotyped | not genotyped | significant | 1 | 2 | 7 |
| denovoLocus14363 | not genotyped | not genotyped | not genotyped | not genotyped | not genotyped | not genotyped | insignificant | insignificant | significant | not genotyped | 1 | 2 | 7 |
| denovoLocus15341 | insignificant | insignificant | not genotyped | not genotyped | not genotyped | not genotyped | not genotyped | significant | not genotyped | not genotyped | 1 | 2 | 7 |
| denovoLocus15964 | insignificant | not genotyped | insignificant | not genotyped | not genotyped | not genotyped | not genotyped | not genotyped | significant | not genotyped | 1 | 2 | 7 |
| denovoLocus16117 | insignificant | not genotyped | significant | not genotyped | not genotyped | insignificant | not genotyped | not genotyped | not genotyped | not genotyped | 1 | 2 | 7 |
| denovoLocus1630 | not genotyped | insignificant | not genotyped | not genotyped | not genotyped | not genotyped | not genotyped | not genotyped | significant | insignificant | 1 | 2 | 7 |
| denovoLocus17262 | insignificant | not genotyped | insignificant | significant | not genotyped | not genotyped | not genotyped | not genotyped | not genotyped | not genotyped | 1 | 2 | 7 |
| denovoLocus17631 | not genotyped | not genotyped | significant | not genotyped | not genotyped | insignificant | not genotyped | insignificant | not genotyped | not genotyped | 1 | 2 | 7 |
| denovoLocus19387 | not genotyped | insignificant | not genotyped | not genotyped | not genotyped | not genotyped | not genotyped | not genotyped | insignificant | significant | 1 | 2 | 7 |
| denovoLocus20069 | significant | not genotyped | insignificant | not genotyped | insignificant | not genotyped | not genotyped | not genotyped | not genotyped | not genotyped | 1 | 2 | 7 |
| denovoLocus20288 | not genotyped | insignificant | significant | not genotyped | insignificant | not genotyped | not genotyped | not genotyped | not genotyped | not genotyped | 1 | 2 | 7 |
| denovoLocus20501 | not genotyped | insignificant | not genotyped | insignificant | not genotyped | significant | not genotyped | not genotyped | not genotyped | not genotyped | 1 | 2 | 7 |
| denovoLocus20604 | not genotyped | not genotyped | not genotyped | not genotyped | not genotyped | significant | not genotyped | not genotyped | insignificant | insignificant | 1 | 2 | 7 |
| denovoLocus21171 | not genotyped | not genotyped | not genotyped | not genotyped | not genotyped | insignificant | significant | insignificant | not genotyped | not genotyped | 1 | 2 | 7 |
| denovoLocus2130 | insignificant | not genotyped | not genotyped | not genotyped | not genotyped | not genotyped | significant | not genotyped | not genotyped | insignificant | 1 | 2 | 7 |
| denovoLocus25158 | not genotyped | not genotyped | not genotyped | not genotyped | not genotyped | insignificant | significant | not genotyped | insignificant | not genotyped | 1 | 2 | 7 |
| denovoLocus25921 | not genotyped | significant | not genotyped | not genotyped | insignificant | not genotyped | not genotyped | not genotyped | not genotyped | insignificant | 1 | 2 | 7 |
| denovoLocus2721 | not genotyped | not genotyped | not genotyped | not genotyped | not genotyped | insignificant | significant | not genotyped | insignificant | not genotyped | 1 | 2 | 7 |
| denovoLocus29979 | not genotyped | insignificant | not genotyped | not genotyped | not genotyped | not genotyped | significant | not genotyped | insignificant | not genotyped | 1 | 2 | 7 |
| denovoLocus30686 | not genotyped | not genotyped | not genotyped | insignificant | not genotyped | not genotyped | significant | not genotyped | not genotyped | insignificant | 1 | 2 | 7 |
| denovoLocus31123 | not genotyped | not genotyped | not genotyped | not genotyped | not genotyped | significant | not genotyped | insignificant | insignificant | not genotyped | 1 | 2 | 7 |
| denovoLocus316 | not genotyped | not genotyped | not genotyped | not genotyped | insignificant | not genotyped | not genotyped | not genotyped | insignificant | significant | 1 | 2 | 7 |
| denovoLocus3229 | insignificant | insignificant | not genotyped | not genotyped | significant | not genotyped | not genotyped | not genotyped | not genotyped | not genotyped | 1 | 2 | 7 |
| denovoLocus32306 | not genotyped | not genotyped | not genotyped | not genotyped | not genotyped | not genotyped | significant | insignificant | insignificant | not genotyped | 1 | 2 | 7 |
| denovoLocus32937 | not genotyped | not genotyped | insignificant | not genotyped | not genotyped | insignificant | not genotyped | significant | not genotyped | not genotyped | 1 | 2 | 7 |
| denovoLocus33215 | insignificant | insignificant | not genotyped | significant | not genotyped | not genotyped | not genotyped | not genotyped | not genotyped | not genotyped | 1 | 2 | 7 |
| denovoLocus35011 | insignificant | not genotyped | not genotyped | not genotyped | significant | not genotyped | not genotyped | insignificant | not genotyped | not genotyped | 1 | 2 | 7 |
| denovoLocus35689 | insignificant | not genotyped | not genotyped | not genotyped | not genotyped | not genotyped | significant | not genotyped | not genotyped | insignificant | 1 | 2 | 7 |
| denovoLocus35770 | not genotyped | not genotyped | not genotyped | insignificant | not genotyped | not genotyped | significant | not genotyped | insignificant | not genotyped | 1 | 2 | 7 |
| denovoLocus36670 | insignificant | not genotyped | insignificant | not genotyped | not genotyped | not genotyped | not genotyped | not genotyped | not genotyped | significant | 1 | 2 | 7 |
| denovoLocus38271 | insignificant | not genotyped | not genotyped | not genotyped | significant | not genotyped | not genotyped | insignificant | not genotyped | not genotyped | 1 | 2 | 7 |
| denovoLocus3902 | significant | not genotyped | not genotyped | not genotyped | insignificant | not genotyped | not genotyped | insignificant | not genotyped | not genotyped | 1 | 2 | 7 |
| denovoLocus42180 | not genotyped | not genotyped | insignificant | not genotyped | not genotyped | not genotyped | significant | insignificant | not genotyped | not genotyped | 1 | 2 | 7 |
| denovoLocus42732 | not genotyped | not genotyped | not genotyped | not genotyped | insignificant | significant | not genotyped | not genotyped | insignificant | not genotyped | 1 | 2 | 7 |
| denovoLocus43191 | not genotyped | not genotyped | not genotyped | insignificant | not genotyped | insignificant | significant | not genotyped | not genotyped | not genotyped | 1 | 2 | 7 |
| denovoLocus43948 | not genotyped | not genotyped | not genotyped | not genotyped | not genotyped | insignificant | not genotyped | insignificant | significant | not genotyped | 1 | 2 | 7 |
| denovoLocus4494 | insignificant | insignificant | not genotyped | not genotyped | not genotyped | not genotyped | not genotyped | not genotyped | not genotyped | significant | 1 | 2 | 7 |
| denovoLocus45728 | not genotyped | insignificant | not genotyped | not genotyped | not genotyped | significant | not genotyped | insignificant | not genotyped | not genotyped | 1 | 2 | 7 |
| denovoLocus46973 | not genotyped | not genotyped | insignificant | significant | not genotyped | not genotyped | not genotyped | insignificant | not genotyped | not genotyped | 1 | 2 | 7 |
| denovoLocus48292 | not genotyped | not genotyped | insignificant | not genotyped | not genotyped | insignificant | not genotyped | not genotyped | not genotyped | significant | 1 | 2 | 7 |
| denovoLocus49500 | not genotyped | not genotyped | not genotyped | insignificant | not genotyped | not genotyped | insignificant | significant | not genotyped | not genotyped | 1 | 2 | 7 |
| denovoLocus51347 | not genotyped | not genotyped | not genotyped | not genotyped | insignificant | not genotyped | not genotyped | not genotyped | insignificant | significant | 1 | 2 | 7 |
| denovoLocus5165 | not genotyped | insignificant | not genotyped | not genotyped | not genotyped | not genotyped | significant | not genotyped | insignificant | not genotyped | 1 | 2 | 7 |
| denovoLocus52641 | not genotyped | insignificant | not genotyped | significant | not genotyped | insignificant | not genotyped | not genotyped | not genotyped | not genotyped | 1 | 2 | 7 |
| denovoLocus5451 | insignificant | not genotyped | not genotyped | significant | insignificant | not genotyped | not genotyped | not genotyped | not genotyped | not genotyped | 1 | 2 | 7 |
| denovoLocus55351 | not genotyped | insignificant | not genotyped | not genotyped | not genotyped | not genotyped | not genotyped | insignificant | significant | not genotyped | 1 | 2 | 7 |
| denovoLocus55935 | insignificant | not genotyped | not genotyped | not genotyped | not genotyped | insignificant | not genotyped | not genotyped | not genotyped | significant | 1 | 2 | 7 |
| denovoLocus58618 | insignificant | not genotyped | not genotyped | not genotyped | significant | not genotyped | not genotyped | insignificant | not genotyped | not genotyped | 1 | 2 | 7 |
| denovoLocus6075 | not genotyped | not genotyped | not genotyped | insignificant | not genotyped | not genotyped | significant | insignificant | not genotyped | not genotyped | 1 | 2 | 7 |
| denovoLocus60891 | not genotyped | not genotyped | not genotyped | not genotyped | insignificant | insignificant | not genotyped | not genotyped | significant | not genotyped | 1 | 2 | 7 |
| denovoLocus62540 | not genotyped | not genotyped | not genotyped | insignificant | insignificant | not genotyped | significant | not genotyped | not genotyped | not genotyped | 1 | 2 | 7 |
| denovoLocus6289 | not genotyped | not genotyped | not genotyped | not genotyped | insignificant | not genotyped | significant | not genotyped | insignificant | not genotyped | 1 | 2 | 7 |
| denovoLocus6412 | not genotyped | not genotyped | not genotyped | insignificant | not genotyped | not genotyped | significant | insignificant | not genotyped | not genotyped | 1 | 2 | 7 |
| denovoLocus67787 | insignificant | not genotyped | not genotyped | not genotyped | not genotyped | not genotyped | not genotyped | insignificant | not genotyped | significant | 1 | 2 | 7 |
| denovoLocus69287 | not genotyped | insignificant | not genotyped | not genotyped | not genotyped | insignificant | significant | not genotyped | not genotyped | not genotyped | 1 | 2 | 7 |
| denovoLocus69799 | not genotyped | not genotyped | not genotyped | not genotyped | insignificant | insignificant | not genotyped | not genotyped | not genotyped | significant | 1 | 2 | 7 |
| denovoLocus7124 | not genotyped | not genotyped | not genotyped | not genotyped | not genotyped | not genotyped | significant | insignificant | not genotyped | insignificant | 1 | 2 | 7 |
| denovoLocus72764 | insignificant | not genotyped | not genotyped | significant | not genotyped | not genotyped | not genotyped | not genotyped | not genotyped | insignificant | 1 | 2 | 7 |
| denovoLocus7870 | not genotyped | not genotyped | not genotyped | not genotyped | not genotyped | significant | insignificant | insignificant | not genotyped | not genotyped | 1 | 2 | 7 |
| denovoLocus78720 | insignificant | not genotyped | not genotyped | not genotyped | significant | insignificant | not genotyped | not genotyped | not genotyped | not genotyped | 1 | 2 | 7 |
| denovoLocus8163 | not genotyped | insignificant | not genotyped | not genotyped | significant | not genotyped | not genotyped | insignificant | not genotyped | not genotyped | 1 | 2 | 7 |
| denovoLocus81633 | not genotyped | not genotyped | not genotyped | significant | not genotyped | not genotyped | not genotyped | insignificant | insignificant | not genotyped | 1 | 2 | 7 |
| denovoLocus83367 | not genotyped | not genotyped | insignificant | not genotyped | not genotyped | significant | not genotyped | not genotyped | not genotyped | insignificant | 1 | 2 | 7 |
| denovoLocus93054 | not genotyped | not genotyped | not genotyped | not genotyped | insignificant | not genotyped | significant | not genotyped | not genotyped | insignificant | 1 | 2 | 7 |
| denovoLocus931 | not genotyped | not genotyped | not genotyped | significant | not genotyped | not genotyped | not genotyped | insignificant | insignificant | not genotyped | 1 | 2 | 7 |
| denovoLocus9331 | not genotyped | not genotyped | not genotyped | not genotyped | insignificant | insignificant | significant | not genotyped | not genotyped | not genotyped | 1 | 2 | 7 |
| denovoLocus9726 | not genotyped | not genotyped | not genotyped | insignificant | not genotyped | not genotyped | significant | not genotyped | insignificant | not genotyped | 1 | 2 | 7 |
| denovoLocus99820 | insignificant | not genotyped | not genotyped | insignificant | not genotyped | not genotyped | significant | not genotyped | not genotyped | not genotyped | 1 | 2 | 7 |
| denovoLocus10036 | not genotyped | not genotyped | not genotyped | significant | not genotyped | not genotyped | not genotyped | not genotyped | insignificant | not genotyped | 1 | 1 | 8 |
| denovoLocus10522 | not genotyped | not genotyped | not genotyped | not genotyped | not genotyped | not genotyped | insignificant | not genotyped | significant | not genotyped | 1 | 1 | 8 |
| denovoLocus10990 | not genotyped | not genotyped | insignificant | not genotyped | not genotyped | not genotyped | significant | not genotyped | not genotyped | not genotyped | 1 | 1 | 8 |
| denovoLocus109919 | not genotyped | insignificant | not genotyped | not genotyped | not genotyped | not genotyped | not genotyped | significant | not genotyped | not genotyped | 1 | 1 | 8 |
| denovoLocus11601 | not genotyped | not genotyped | not genotyped | not genotyped | not genotyped | not genotyped | insignificant | not genotyped | not genotyped | significant | 1 | 1 | 8 |
| denovoLocus12254 | not genotyped | not genotyped | not genotyped | significant | not genotyped | not genotyped | not genotyped | not genotyped | insignificant | not genotyped | 1 | 1 | 8 |
| denovoLocus13057 | not genotyped | not genotyped | not genotyped | not genotyped | not genotyped | not genotyped | significant | insignificant | not genotyped | not genotyped | 1 | 1 | 8 |
| denovoLocus13546 | not genotyped | not genotyped | not genotyped | not genotyped | not genotyped | insignificant | not genotyped | not genotyped | significant | not genotyped | 1 | 1 | 8 |
| denovoLocus14112 | not genotyped | insignificant | not genotyped | not genotyped | not genotyped | not genotyped | significant | not genotyped | not genotyped | not genotyped | 1 | 1 | 8 |
| denovoLocus17085 | not genotyped | not genotyped | not genotyped | not genotyped | not genotyped | insignificant | significant | not genotyped | not genotyped | not genotyped | 1 | 1 | 8 |
| denovoLocus1714 | not genotyped | insignificant | not genotyped | not genotyped | not genotyped | not genotyped | not genotyped | not genotyped | significant | not genotyped | 1 | 1 | 8 |
| denovoLocus1726 | not genotyped | not genotyped | significant | not genotyped | not genotyped | insignificant | not genotyped | not genotyped | not genotyped | not genotyped | 1 | 1 | 8 |
| denovoLocus18331 | not genotyped | not genotyped | not genotyped | not genotyped | not genotyped | insignificant | significant | not genotyped | not genotyped | not genotyped | 1 | 1 | 8 |
| denovoLocus1946 | not genotyped | not genotyped | not genotyped | not genotyped | not genotyped | not genotyped | not genotyped | insignificant | significant | not genotyped | 1 | 1 | 8 |
| denovoLocus19806 | not genotyped | not genotyped | not genotyped | not genotyped | not genotyped | insignificant | not genotyped | not genotyped | significant | not genotyped | 1 | 1 | 8 |
| denovoLocus21883 | not genotyped | not genotyped | not genotyped | not genotyped | not genotyped | insignificant | significant | not genotyped | not genotyped | not genotyped | 1 | 1 | 8 |
| denovoLocus2216 | not genotyped | not genotyped | not genotyped | not genotyped | insignificant | not genotyped | significant | not genotyped | not genotyped | not genotyped | 1 | 1 | 8 |
| denovoLocus22238 | not genotyped | not genotyped | not genotyped | not genotyped | not genotyped | not genotyped | insignificant | significant | not genotyped | not genotyped | 1 | 1 | 8 |
| denovoLocus22764 | not genotyped | insignificant | not genotyped | not genotyped | not genotyped | significant | not genotyped | not genotyped | not genotyped | not genotyped | 1 | 1 | 8 |
| denovoLocus2663 | not genotyped | significant | not genotyped | not genotyped | not genotyped | not genotyped | not genotyped | not genotyped | not genotyped | insignificant | 1 | 1 | 8 |
| denovoLocus27270 | not genotyped | not genotyped | not genotyped | significant | not genotyped | not genotyped | not genotyped | not genotyped | insignificant | not genotyped | 1 | 1 | 8 |
| denovoLocus3005 | not genotyped | not genotyped | not genotyped | significant | not genotyped | insignificant | not genotyped | not genotyped | not genotyped | not genotyped | 1 | 1 | 8 |
| denovoLocus3047 | not genotyped | not genotyped | not genotyped | not genotyped | not genotyped | insignificant | significant | not genotyped | not genotyped | not genotyped | 1 | 1 | 8 |
| denovoLocus3048 | not genotyped | not genotyped | not genotyped | insignificant | not genotyped | not genotyped | significant | not genotyped | not genotyped | not genotyped | 1 | 1 | 8 |
| denovoLocus3057 | not genotyped | not genotyped | not genotyped | significant | not genotyped | not genotyped | not genotyped | insignificant | not genotyped | not genotyped | 1 | 1 | 8 |
| denovoLocus33862 | not genotyped | insignificant | not genotyped | not genotyped | not genotyped | not genotyped | not genotyped | not genotyped | not genotyped | significant | 1 | 1 | 8 |
| denovoLocus36794 | not genotyped | not genotyped | significant | insignificant | not genotyped | not genotyped | not genotyped | not genotyped | not genotyped | not genotyped | 1 | 1 | 8 |
| denovoLocus4368 | insignificant | not genotyped | not genotyped | not genotyped | not genotyped | not genotyped | significant | not genotyped | not genotyped | not genotyped | 1 | 1 | 8 |
| denovoLocus44226 | not genotyped | not genotyped | not genotyped | not genotyped | not genotyped | insignificant | significant | not genotyped | not genotyped | not genotyped | 1 | 1 | 8 |
| denovoLocus4463 | not genotyped | not genotyped | not genotyped | not genotyped | insignificant | not genotyped | not genotyped | not genotyped | significant | not genotyped | 1 | 1 | 8 |
| denovoLocus44821 | not genotyped | not genotyped | not genotyped | insignificant | not genotyped | significant | not genotyped | not genotyped | not genotyped | not genotyped | 1 | 1 | 8 |
| denovoLocus46507 | not genotyped | insignificant | not genotyped | significant | not genotyped | not genotyped | not genotyped | not genotyped | not genotyped | not genotyped | 1 | 1 | 8 |
| denovoLocus516 | not genotyped | not genotyped | not genotyped | not genotyped | not genotyped | not genotyped | significant | not genotyped | insignificant | not genotyped | 1 | 1 | 8 |
| denovoLocus53060 | not genotyped | significant | not genotyped | not genotyped | not genotyped | not genotyped | not genotyped | insignificant | not genotyped | not genotyped | 1 | 1 | 8 |
| denovoLocus53379 | significant | insignificant | not genotyped | not genotyped | not genotyped | not genotyped | not genotyped | not genotyped | not genotyped | not genotyped | 1 | 1 | 8 |
| denovoLocus53663 | not genotyped | not genotyped | not genotyped | not genotyped | not genotyped | insignificant | not genotyped | not genotyped | not genotyped | significant | 1 | 1 | 8 |
| denovoLocus5413 | not genotyped | insignificant | not genotyped | not genotyped | not genotyped | not genotyped | significant | not genotyped | not genotyped | not genotyped | 1 | 1 | 8 |
| denovoLocus55778 | not genotyped | not genotyped | not genotyped | not genotyped | not genotyped | not genotyped | significant | insignificant | not genotyped | not genotyped | 1 | 1 | 8 |
| denovoLocus57324 | not genotyped | not genotyped | significant | not genotyped | not genotyped | insignificant | not genotyped | not genotyped | not genotyped | not genotyped | 1 | 1 | 8 |
| denovoLocus61545 | significant | insignificant | not genotyped | not genotyped | not genotyped | not genotyped | not genotyped | not genotyped | not genotyped | not genotyped | 1 | 1 | 8 |
| denovoLocus61946 | not genotyped | not genotyped | not genotyped | significant | not genotyped | not genotyped | not genotyped | insignificant | not genotyped | not genotyped | 1 | 1 | 8 |
| denovoLocus63114 | not genotyped | insignificant | significant | not genotyped | not genotyped | not genotyped | not genotyped | not genotyped | not genotyped | not genotyped | 1 | 1 | 8 |
| denovoLocus65281 | not genotyped | not genotyped | not genotyped | not genotyped | not genotyped | not genotyped | significant | not genotyped | not genotyped | insignificant | 1 | 1 | 8 |
| denovoLocus67582 | not genotyped | not genotyped | not genotyped | not genotyped | not genotyped | insignificant | not genotyped | not genotyped | significant | not genotyped | 1 | 1 | 8 |
| denovoLocus68991 | significant | insignificant | not genotyped | not genotyped | not genotyped | not genotyped | not genotyped | not genotyped | not genotyped | not genotyped | 1 | 1 | 8 |
| denovoLocus69894 | not genotyped | not genotyped | not genotyped | insignificant | not genotyped | not genotyped | not genotyped | significant | not genotyped | not genotyped | 1 | 1 | 8 |
| denovoLocus70437 | not genotyped | not genotyped | not genotyped | not genotyped | not genotyped | not genotyped | significant | not genotyped | insignificant | not genotyped | 1 | 1 | 8 |
| denovoLocus72953 | not genotyped | not genotyped | not genotyped | not genotyped | not genotyped | not genotyped | not genotyped | insignificant | significant | not genotyped | 1 | 1 | 8 |
| denovoLocus75591 | not genotyped | not genotyped | not genotyped | not genotyped | not genotyped | not genotyped | significant | not genotyped | insignificant | not genotyped | 1 | 1 | 8 |
| denovoLocus76722 | not genotyped | not genotyped | insignificant | not genotyped | not genotyped | significant | not genotyped | not genotyped | not genotyped | not genotyped | 1 | 1 | 8 |
| denovoLocus79736 | not genotyped | not genotyped | insignificant | significant | not genotyped | not genotyped | not genotyped | not genotyped | not genotyped | not genotyped | 1 | 1 | 8 |
| denovoLocus82613 | not genotyped | not genotyped | not genotyped | not genotyped | not genotyped | insignificant | significant | not genotyped | not genotyped | not genotyped | 1 | 1 | 8 |
| denovoLocus82746 | not genotyped | not genotyped | not genotyped | not genotyped | insignificant | not genotyped | significant | not genotyped | not genotyped | not genotyped | 1 | 1 | 8 |
| denovoLocus83814 | not genotyped | insignificant | not genotyped | not genotyped | not genotyped | not genotyped | not genotyped | not genotyped | not genotyped | significant | 1 | 1 | 8 |
| denovoLocus843 | significant | not genotyped | insignificant | not genotyped | not genotyped | not genotyped | not genotyped | not genotyped | not genotyped | not genotyped | 1 | 1 | 8 |
| denovoLocus91789 | not genotyped | not genotyped | not genotyped | not genotyped | not genotyped | not genotyped | not genotyped | not genotyped | significant | insignificant | 1 | 1 | 8 |
| denovoLocus93162 | not genotyped | not genotyped | not genotyped | not genotyped | not genotyped | not genotyped | not genotyped | insignificant | significant | not genotyped | 1 | 1 | 8 |
| denovoLocus96767 | significant | not genotyped | insignificant | not genotyped | not genotyped | not genotyped | not genotyped | not genotyped | not genotyped | not genotyped | 1 | 1 | 8 |
| denovoLocus9772 | not genotyped | not genotyped | insignificant | not genotyped | not genotyped | not genotyped | significant | not genotyped | not genotyped | not genotyped | 1 | 1 | 8 |
| denovoLocus10439 | not genotyped | not genotyped | not genotyped | not genotyped | not genotyped | not genotyped | significant | not genotyped | not genotyped | not genotyped | 1 | 0 | 9 |
| denovoLocus11086 | not genotyped | not genotyped | significant | not genotyped | not genotyped | not genotyped | not genotyped | not genotyped | not genotyped | not genotyped | 1 | 0 | 9 |
| denovoLocus11212 | not genotyped | not genotyped | not genotyped | not genotyped | significant | not genotyped | not genotyped | not genotyped | not genotyped | not genotyped | 1 | 0 | 9 |
| denovoLocus11967 | not genotyped | not genotyped | not genotyped | not genotyped | not genotyped | not genotyped | not genotyped | not genotyped | not genotyped | significant | 1 | 0 | 9 |
| denovoLocus13048 | not genotyped | not genotyped | not genotyped | not genotyped | not genotyped | not genotyped | significant | not genotyped | not genotyped | not genotyped | 1 | 0 | 9 |
| denovoLocus13686 | not genotyped | not genotyped | not genotyped | not genotyped | not genotyped | not genotyped | not genotyped | not genotyped | significant | not genotyped | 1 | 0 | 9 |
| denovoLocus14859 | not genotyped | not genotyped | not genotyped | not genotyped | not genotyped | not genotyped | significant | not genotyped | not genotyped | not genotyped | 1 | 0 | 9 |
| denovoLocus14860 | not genotyped | not genotyped | not genotyped | not genotyped | not genotyped | not genotyped | not genotyped | not genotyped | significant | not genotyped | 1 | 0 | 9 |
| denovoLocus15635 | not genotyped | not genotyped | not genotyped | not genotyped | not genotyped | not genotyped | significant | not genotyped | not genotyped | not genotyped | 1 | 0 | 9 |
| denovoLocus15827 | not genotyped | not genotyped | not genotyped | not genotyped | not genotyped | not genotyped | not genotyped | not genotyped | significant | not genotyped | 1 | 0 | 9 |
| denovoLocus16722 | not genotyped | not genotyped | not genotyped | not genotyped | not genotyped | significant | not genotyped | not genotyped | not genotyped | not genotyped | 1 | 0 | 9 |
| denovoLocus17876 | not genotyped | not genotyped | not genotyped | not genotyped | not genotyped | not genotyped | not genotyped | not genotyped | significant | not genotyped | 1 | 0 | 9 |
| denovoLocus1788 | not genotyped | not genotyped | not genotyped | not genotyped | not genotyped | not genotyped | significant | not genotyped | not genotyped | not genotyped | 1 | 0 | 9 |
| denovoLocus18385 | not genotyped | not genotyped | not genotyped | not genotyped | not genotyped | not genotyped | not genotyped | not genotyped | not genotyped | significant | 1 | 0 | 9 |
| denovoLocus19953 | not genotyped | not genotyped | not genotyped | not genotyped | not genotyped | not genotyped | significant | not genotyped | not genotyped | not genotyped | 1 | 0 | 9 |
| denovoLocus2162 | not genotyped | not genotyped | not genotyped | not genotyped | not genotyped | not genotyped | significant | not genotyped | not genotyped | not genotyped | 1 | 0 | 9 |
| denovoLocus26344 | significant | not genotyped | not genotyped | not genotyped | not genotyped | not genotyped | not genotyped | not genotyped | not genotyped | not genotyped | 1 | 0 | 9 |
| denovoLocus26426 | not genotyped | not genotyped | not genotyped | not genotyped | significant | not genotyped | not genotyped | not genotyped | not genotyped | not genotyped | 1 | 0 | 9 |
| denovoLocus27216 | not genotyped | not genotyped | not genotyped | not genotyped | not genotyped | not genotyped | significant | not genotyped | not genotyped | not genotyped | 1 | 0 | 9 |
| denovoLocus29401 | not genotyped | not genotyped | not genotyped | not genotyped | not genotyped | not genotyped | not genotyped | significant | not genotyped | not genotyped | 1 | 0 | 9 |
| denovoLocus31740 | not genotyped | not genotyped | not genotyped | not genotyped | not genotyped | not genotyped | significant | not genotyped | not genotyped | not genotyped | 1 | 0 | 9 |
| denovoLocus34585 | not genotyped | not genotyped | not genotyped | not genotyped | not genotyped | not genotyped | significant | not genotyped | not genotyped | not genotyped | 1 | 0 | 9 |
| denovoLocus35478 | not genotyped | not genotyped | not genotyped | not genotyped | not genotyped | not genotyped | not genotyped | not genotyped | significant | not genotyped | 1 | 0 | 9 |
| denovoLocus3726 | not genotyped | not genotyped | not genotyped | not genotyped | not genotyped | not genotyped | significant | not genotyped | not genotyped | not genotyped | 1 | 0 | 9 |
| denovoLocus38389 | not genotyped | not genotyped | not genotyped | not genotyped | not genotyped | significant | not genotyped | not genotyped | not genotyped | not genotyped | 1 | 0 | 9 |
| denovoLocus4554 | not genotyped | not genotyped | not genotyped | not genotyped | not genotyped | significant | not genotyped | not genotyped | not genotyped | not genotyped | 1 | 0 | 9 |
| denovoLocus4568 | not genotyped | not genotyped | not genotyped | not genotyped | not genotyped | not genotyped | not genotyped | significant | not genotyped | not genotyped | 1 | 0 | 9 |
| denovoLocus46246 | not genotyped | not genotyped | not genotyped | not genotyped | not genotyped | not genotyped | not genotyped | not genotyped | not genotyped | significant | 1 | 0 | 9 |
| denovoLocus51020 | not genotyped | not genotyped | not genotyped | significant | not genotyped | not genotyped | not genotyped | not genotyped | not genotyped | not genotyped | 1 | 0 | 9 |
| denovoLocus53405 | not genotyped | not genotyped | not genotyped | not genotyped | not genotyped | not genotyped | not genotyped | significant | not genotyped | not genotyped | 1 | 0 | 9 |
| denovoLocus57558 | not genotyped | not genotyped | not genotyped | significant | not genotyped | not genotyped | not genotyped | not genotyped | not genotyped | not genotyped | 1 | 0 | 9 |
| denovoLocus61081 | not genotyped | not genotyped | not genotyped | significant | not genotyped | not genotyped | not genotyped | not genotyped | not genotyped | not genotyped | 1 | 0 | 9 |
| denovoLocus62877 | not genotyped | not genotyped | not genotyped | not genotyped | not genotyped | significant | not genotyped | not genotyped | not genotyped | not genotyped | 1 | 0 | 9 |
| denovoLocus65455 | not genotyped | not genotyped | not genotyped | not genotyped | not genotyped | significant | not genotyped | not genotyped | not genotyped | not genotyped | 1 | 0 | 9 |
| denovoLocus67766 | not genotyped | not genotyped | not genotyped | not genotyped | not genotyped | not genotyped | significant | not genotyped | not genotyped | not genotyped | 1 | 0 | 9 |
| denovoLocus68579 | significant | not genotyped | not genotyped | not genotyped | not genotyped | not genotyped | not genotyped | not genotyped | not genotyped | not genotyped | 1 | 0 | 9 |
| denovoLocus69090 | not genotyped | not genotyped | not genotyped | not genotyped | not genotyped | not genotyped | significant | not genotyped | not genotyped | not genotyped | 1 | 0 | 9 |
| denovoLocus69253 | not genotyped | not genotyped | not genotyped | not genotyped | not genotyped | not genotyped | not genotyped | not genotyped | significant | not genotyped | 1 | 0 | 9 |
| denovoLocus70763 | not genotyped | not genotyped | not genotyped | not genotyped | not genotyped | not genotyped | not genotyped | not genotyped | not genotyped | significant | 1 | 0 | 9 |
| denovoLocus72942 | not genotyped | not genotyped | not genotyped | not genotyped | not genotyped | not genotyped | significant | not genotyped | not genotyped | not genotyped | 1 | 0 | 9 |
| denovoLocus75905 | not genotyped | not genotyped | significant | not genotyped | not genotyped | not genotyped | not genotyped | not genotyped | not genotyped | not genotyped | 1 | 0 | 9 |
| denovoLocus78662 | significant | not genotyped | not genotyped | not genotyped | not genotyped | not genotyped | not genotyped | not genotyped | not genotyped | not genotyped | 1 | 0 | 9 |
| denovoLocus80287 | not genotyped | not genotyped | not genotyped | not genotyped | not genotyped | not genotyped | not genotyped | not genotyped | significant | not genotyped | 1 | 0 | 9 |
| denovoLocus8116 | not genotyped | not genotyped | not genotyped | not genotyped | not genotyped | not genotyped | not genotyped | not genotyped | not genotyped | significant | 1 | 0 | 9 |
| denovoLocus84634 | not genotyped | not genotyped | not genotyped | not genotyped | not genotyped | not genotyped | significant | not genotyped | not genotyped | not genotyped | 1 | 0 | 9 |
| denovoLocus86232 | not genotyped | not genotyped | not genotyped | significant | not genotyped | not genotyped | not genotyped | not genotyped | not genotyped | not genotyped | 1 | 0 | 9 |
| denovoLocus87847 | not genotyped | not genotyped | not genotyped | not genotyped | not genotyped | not genotyped | significant | not genotyped | not genotyped | not genotyped | 1 | 0 | 9 |
| denovoLocus88248 | significant | not genotyped | not genotyped | not genotyped | not genotyped | not genotyped | not genotyped | not genotyped | not genotyped | not genotyped | 1 | 0 | 9 |
|  |  |  |  |  |  |  |  |  |  |  |  | median not genotyped | 5.5 |
|  |  |  |  |  |  |  |  |  |  |  |  | percent of tags found in less than half of crosses | 35.64705882 |
|  |  |  |  |  |  |  |  |  |  |  |  | percent of tags genotyped in 50% of crosses | 64.35294118 |
