## SupplementalFigure1 for "Genetic variation in heat tolerance of the coral *Platygyra daedalea* indicates potential for adaptation to ocean warming"

**Supplementary Material: Graph of the Temperature at Esk Reef during the study taken from the Australian Institute of Marine Science (AIMS).**

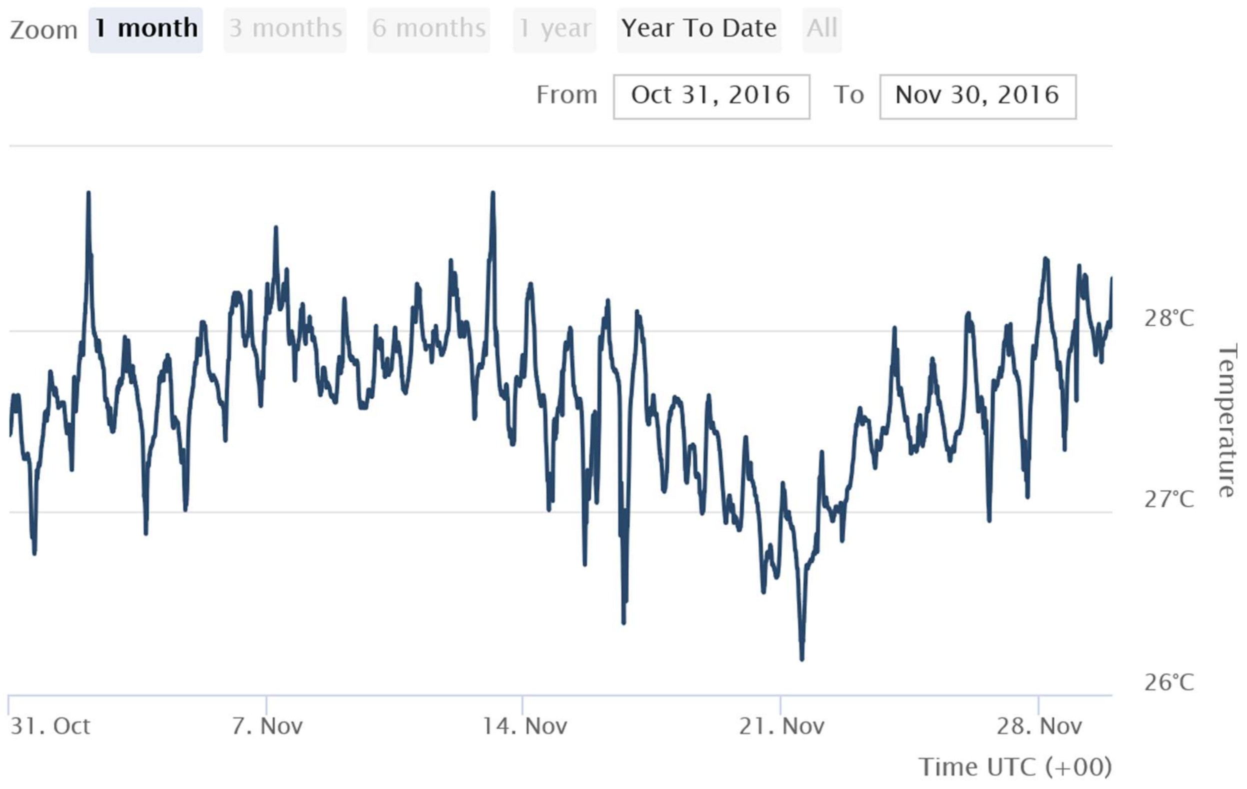


Supplemental Figure 1: Temperature log for Esk Reef. This plot shows the water temperature for the reef from which the parent coral colonies were collected. The mean temperature is 27˚C for this reef, which why we chose this temperature for our control and maintenance temperature for the coral larvae used in this study. Taken directly from the AIMS data access website: https://apps.aims.gov.au/ts-explorer/?series=HAVFL1&fromDate=2016-10-31T00:00:00&thruDate=2016-11-30T00:00:00
